# Molecular features of RNA silencing against phloem-restricted polerovirus TuYV enable amplification of silencing signal from host transcripts

**DOI:** 10.1101/2021.03.19.436175

**Authors:** Marion Clavel, Esther Lechner, Marco Incarbone, Timothée Vincent, Valerie Cognat, Ekaterina Smirnova, Maxime Lecorbeiller, Véronique Brault, Véronique Ziegler-Graff, Pascal Genschik

**Author notes:** For correspondence (MC), (PG).

## Abstract

In plants and some animal lineages, RNA silencing is an efficient and adaptable defense mechanism against viruses. To counter it, viruses encode suppressor proteins that interfere with RNA silencing. Phloem-restricted viruses are spreading at an alarming rate and cause substantial reduction of crop yield, but how they interact with their hosts at the molecular level is still insufficiently understood. Here, we investigate the antiviral response against phloem-restricted turnip yellows virus (TuYV) in the model plant *Arabidopsis thaliana*. Using a combination of genetics, deep sequencing, and mechanical vasculature enrichment, we show that the main axis of silencing active against TuYV involves 22-nt vsiRNA production by DCL2, and their preferential loading into AGO1. Unexpectedly, and despite the viral encoded VSR P0 previously shown to mediate degradation of AGO proteins, vascular AGO1 undergoes specific post-translational stabilization during TuYV infection. We also identify vascular novel secondary siRNA produced from conserved plant transcripts and initiated by DCL2-processed AGO1-loaded vsiRNA, supporting a viral strategy to modulate host response. Collectively, our work uncovers the complexity of antiviral RNA silencing against phloem-restricted TuYV and prompts a re-assessment of the role of its suppressor of silencing P0 during genuine infection.

## Introduction

To defend themselves against pathogens, plants have developed a molecular arsenal allowing them to detect and resist the incoming threat. In turn, pathogens have adopted numerous evasions strategies and can exploit plant defenses with the resulting arms race leading to complex and ever-changing host-microbe interactions. One such focal point of plant defense and viral counter-defense is RNA silencing, with traces of such interactions evident across both plant and virus diversity (Pumplin & Voinnet, 2013; Yang & Li, 2018). All RNA silencing pathways rest on the action of small RNA (sRNA) whose production depends on the enzymatic activity of RNAse III proteins called Dicer-like (DCLs). In the case of RNA viruses, production of viral small interfering (vsi)RNA is triggered by double-stranded (ds)RNA replication intermediates or intramolecular foldback structures in the viral genome mainly by the action of DCL4 and DCL2, generating 21- and 22-nt vsiRNA duplexes respectively (Blevins *et al*, 2006; Bouché *et al*, 2006; Deleris *et al*, 2006). This first layer of detection and degradation is reinforced by specialized effector proteins called ARGONAUTE (AGO) that associate with the vsiRNA to form the antiviral RNA-inducted silencing complex (RISC) (Carbonell & Carrington, 2015). The RISC complex can target RNA in a sequence-specific manner, leading to endonucleolytic cleavage (slicing) catalyzed by the AGO and/or via translational repression coupled with mRNA decay (Poulsen *et al*, 2013). The silencing signal can further be amplified through the conversion of single stranded (ss)RNA targets into dsRNA thanks to host-encoded RNA-dependent RNA polymerase (RDR) proteins, providing new template for secondary vsiRNA production by DCLs, that are important to achieve optimal silencing for some viruses (Qu *et al*, 2008; Donaire *et al*, 2008; Wang *et al*, 2010; Garcia-Ruiz *et al*, 2010). Its layered, self-reinforcing and sequence specific mechanism makes RNA silencing a particularly potent immune system.

In order to foil this mechanism most, if not all, plant viruses deploy specialized proteins that are known as viral suppressor of RNA silencing, or VSRs. VSRs across different virus families that are highly diverse have adopted various strategies to impair different steps of RNA silencing (Incarbone & Dunoyer, 2013; Pumplin & Voinnet, 2013). Their deployment usually results in the abrogation of the movement of vsiRNA and therefore of plant immunization (Guo & Ding, 2002; Schott *et al*, 2012; Incarbone *et al*, 2017), resulting in the accumulation of high amounts of the viral genome often accompanied by strong symptoms. Consequently, viruses for which VSR activity has been inactivated are strongly affected in their ability to move long distances and achieve systemic infection (Havelda *et al*, 2003; Bayne *et al*, 2005; Wang *et al*, 2011; Chiba *et al*, 2013; Deleris *et al*, 2006; Garcia-Ruiz *et al*, 2010; Incarbone *et al*, 2017). This observation has been instrumental in deciphering the silencing components involved in plant defense, since their mutation leads to a rescue of viral movement. However, a causal link between VSR and movement has only been established for comparatively few viruses, while many more known VSRs await *in vivo* characterization in the context of infection.

The P0 VSR of phloem-restricted poleroviruses presents an interesting case: while the intricacies of its mode of action are understood, they do not necessarily reconcile with observations made in the context of infection nor with the behaviour of natural variants of the protein. By hijacking the S phase kinase associated protein 1 (SKP1) - Cullin 1 – F-box (SCF) E3 ubiquitin ligase complex and enforcing degradation of most AGO proteins, P0 impedes the formation of vsiRNA-RISC (Pazhouhandeh *et al*, 2006; Baumberger *et al*, 2007; Bortolamiol *et al*, 2007). In the case of the AGO1 protein, it has been shown that it’s interaction with P0 leads to its ubiquitination and vacuolar degradation (Derrien *et al*, 2012; Michaeli *et al*, 2019). This strategy seems to be particularly effective, as in heterologous assays in *N. benthamiana* (patch assays) P0 of turnip yellows virus (TuYV) is able to suppress the potent RNA silencing reaction to transgenes, enabling strong and persistent expression of GFP. Accordingly, strong dosage of P0 leads to developmental phenotypes reminiscent of *AGO1* knockout plants (Bohmert *et al*, 1998) because P0 also disables miRNA-RISC assembly (Bortolamiol *et al*, 2007; Fusaro *et al*, 2012).

On the opposite spectrum, studies employing a TuYY that is unable to produce P0 show that it is dispensable for systemic infection (Ziegler-Graff *et al*, 1996) while the resulting systemic infection by the wild type (WT) virus is asymptomatic in Arabidopsis (Bortolamiol-Bécet *et al*, 2018). Furthermore, P0 proteins from polerovirus isolates collected throughout the world display varying degrees of silencing efficiency in patch assay, ranging from strong (Mangwende *et al*, 2009; Han *et al*, 2010; Liu *et al*, 2012; Chen *et al*, 2016; Fusaro *et al*, 2012) and moderate (Delfosse *et al*, 2014; Cascardo *et al*, 2015; Almasi *et al*, 2015) to non-existent (Kozlowska-Makulska *et al*, 2010; Han *et al*, 2010). Intriguingly, the start codon of ORF0 encoding P0 of TuYV has a poor initiation context (Mayo & Ziegler-Graff, 1996) which results in leaky scanning by the ribosomes and therefore reduced initiation. Optimization of the 5’ context decreases viral RNA accumulation and leads to second-site mutations of the start codon that restore low translation initiation in the systemic progeny (Pfeffer *et al*, 2002). Similarly, potato leafroll virus (PLRV) genomic leader sequence exerts an inhibitory effect on translation of the downstream ORF0 and ORF1 (Juszczuk *et al*, 2000). These observations suggest that neither high accumulation nor strong suppression activity are traits that have been selected for during plant-polerovirus coevolution. These contrasting observations raise the question of the role of P0 during infection of the plant host, and of its interaction with the RNA silencing machinery.

By means of a genetic screen, we have recently uncovered a suppressor of the P0-dependent developmental phenotype typically associated with AGO1 degradation (Derrien *et al*, 2018). The single Gly to Asp substitution in the DUF1785 of AGO1 leads to the loss of the SCF^P0^-AGO1 interaction and therefore renders the mutant AGO1 protein non-degradable by P0, which could have potential practical applications. We also showed that this mutation hinders sRNA duplex “unwinding” by AGO1 itself, particularly for perfect duplexes, a hallmark of siRNA rather than miRNA. Accordingly, miRNA-programmed RISC activity is mostly unperturbed by the mutation, leading to a mild developmental phenotype, while endogenous siRNA-programmed RISC is strongly affected, which results in the near-complete loss of secondary siRNA. Thus, this unique allele of *AGO1* allows for unprecedented decoupling of miRNA-guided pathways from siRNA guided ones, which are relied upon to perform antiviral RNA silencing.

Here, we take advantage of this unique allele, *ago1-57*, to investigate the silencing components required for efficient antiviral immunity against TuYV. We show that TuYV-derived vsiRNA are largely channelled toward AGO1, and that its antiviral importance is only evident in some missense mutants. *ago1-57* causes a 50% reduction in systemic TuYV infection by delaying the establishment of the infection in agroinoculated leaves but fails to provide any protection in aphid-mediated infection of the plant vascular system. We further show that TuYV RNA is mainly cleaved into 22-nt vsiRNA by vascular DCL2 in a mechanism that is distinct from that described for turnip crinckle virus (TCV) infection. While transgenic vascular P0 can recapitulate AGO1 degradation in the phloem, AGO1 undergoes vascular post-translational stabilization during TuYV infection in a P0-independent manner. Finally, we show that both DCL2 processing and AGO1 stabilization concur to vsiRNA-AGO1 mediated targeting of a set of conserved endogenous messenger RNA to produce secondary siRNA, highlighting a potential mechanism for viruses to modulate their host’s transcriptome by manipulating antiviral silencing pathways.

## Results

### siRNA-RISC-deficient point mutation in AGO1 highlights its crucial role in antiviral defense against TuYV

We have previously demonstrated that the G371D mutation, carried by the *ago1-57* allele of AGO1, allows evasion from the P0 VSR while simultaneously causing retention of perfect siRNA duplexes, leading to the inhibition of secondary siRNA production (Derrien *et al*, 2018). We therefore addressed the implications of this dual phenotype during TuYV infection in the model plant *Arabidopsis thaliana* (hereafter Arabidopsis), using an infectious clone of TuYV containing an 81 bp insertion of the *AtCHLI1* gene in the 3’ non-coding region of the genome (Bortolamiol-Bécet *et al*, 2018), referred to as TuYVs81. Since the systemic viral propagation is limited to the phloem and can act as a trigger to produce vsiRNA that move cell-to-cell, silencing of the *CHLI1* gene in the neighbouring cells through virus-induced gene silencing (VIGS) typically leads to yellowing around the veins. To assess the importance of AGO1 during TuYV infection and for VIGS, WT (Col-0) and different *ago1* mutant alleles were inoculated with either a WT TuYVs81 or the P0-deficient clone TuYVs81P0^-^ and viral RNA accumulation was assessed in systemic tissues. Intriguingly, while absence of P0 consistently resulted in less viral RNA accumulation relative to the WT TuYVs81, none of the tested *ago1* alleles exhibited the same phenotype. *ago1-27* plants accumulated a similar amount of TuYVs81 RNA to the WT plants, while the *ago1-57* and *ago1-38* plants accumulated moderately more viral RNA (**Figure 1A**). While Col-0 and *ago1-27* plants display clear vein yellowing, *ago1-57* plants presented reduced yellowing despite elevated viral RNA levels, indicating this mutant allele of AGO1 is impaired in its ability to establish VIGS (**Figure 1B**). None of the *ago1-38* plants displayed any vein yellowing, despite the presence of the viral RNA in the systemic leaves. Any VIGS defect observed in presence of the WT virus was exacerbated when infected with TuYVs81P0^-^, resulting in even less vein yellowing in *ago1-57*. We conclude that TuYVs81-based VIGS is sensitive to the amount of trigger RNA, relies on AGO1 for its completion, and that different AGO1 point mutations result in varying VIGS as well as antiviral RNA silencing efficacy.

**Figure 1:**
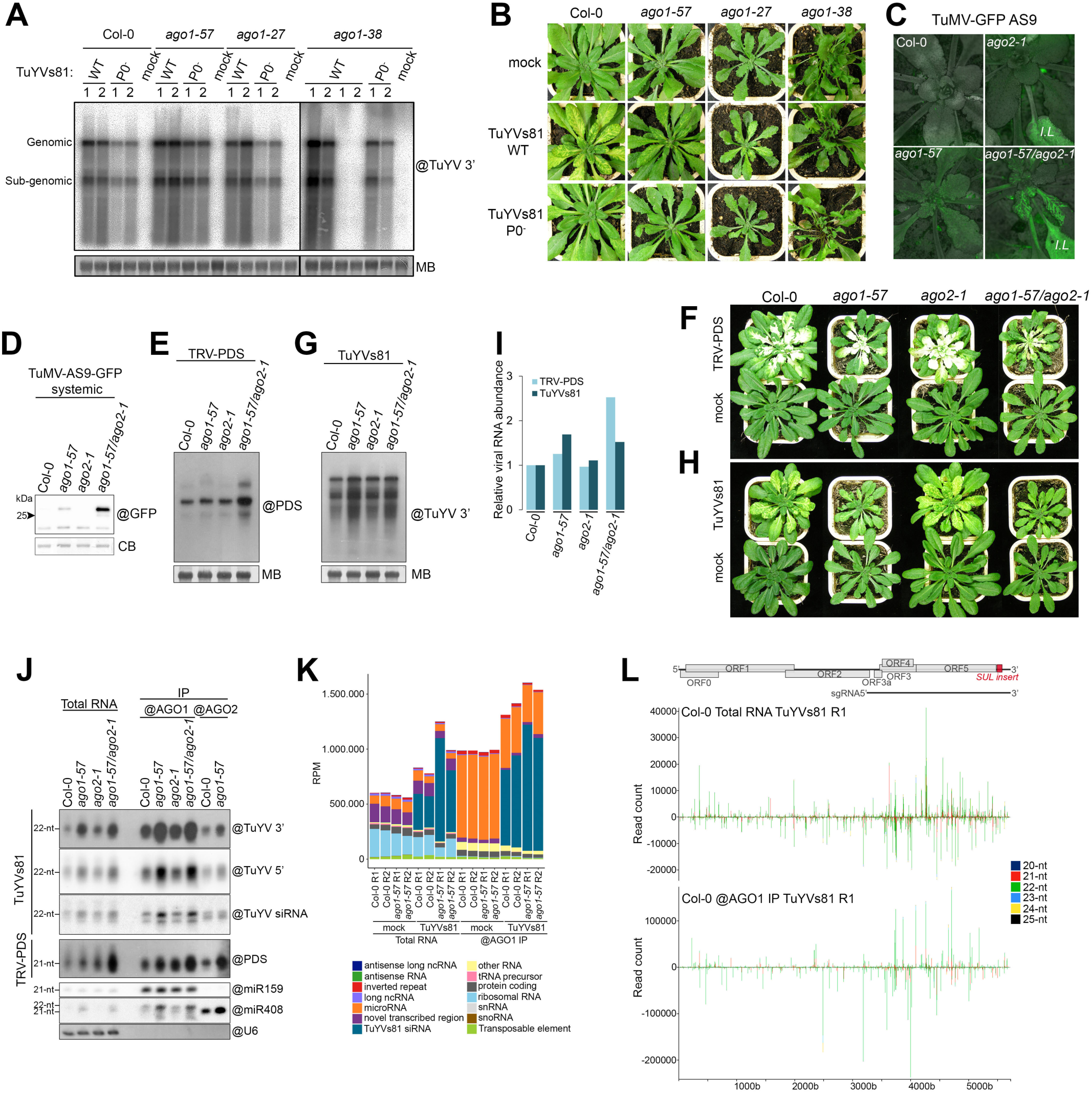
AGO1 is the main Argonaute protein involved in defense against turnip yellows virus. **(A.)** Accumulation of TuYVs81 RNA in systemic leaves of Col-0, *ago1-57*, *ago1-27* and *ago1-38* plants at 19 days post-infiltration (dpi) with either WT or P0-less (P0^-^) virus. Mock stands for mock-inoculated plants. Each lane represents a pool of four to seven individuals from which RNA was extracted from either the youngest rosette leaves (**1**) or older rosette leaves with yellow veins (**2**). For *ago1-38* that did not display any vein yellowing, leaves were sampled in a similar fashion from plants displaying reddened leaves (left) or not (right, no virus accumulation). Viral RNA abundance was measured by RNA gel blot; loading control is obtained by staining the membrane with methylene blue (MB). “@” indicates hybridization with DNA probe against the 3’ part of the TuYV genome. **(B.)** Representative image of the infected genotypes analyzed in **A**. **(C.)** Representative image of the TuMV-AS9-GFP infected plants at 9 dpi. Successful systemic movement is achieved only in *ago1-57* and the double mutant while expression of viral-derived GFP in the inoculated leaves (I.L) is clearly visible. **(D.)** Rescue of the TuMV-AS9-GFP systemic movement in the single *ago1-57* and the double *ago1-57/ago2-1* genetic backgrounds. Systemic leaves of TuMV-AS9-GFP inoculated plants were harvested at 13 dpi (n=5 plants), and GFP protein content was measured by immunoblot. “@” indicates hybridization with GFP antibody, loading control is obtained by post-staining the membrane with coomassie blue (CB). **(E.)** Accumulation of TRV-PDS RNA in systemic leaves in the indicated genotypes at 20 dpi (n=5 plants) measured by RNA gel blot. “@” indicates hybridization with DNA probe against the PDS insert and loading control is obtained by staining the membrane with methylene blue (MB). **(F.)** Representative individuals infected with TRV-PDS displaying systemic leaf whitening due to the silencing of the PDS gene. **(G.)** Accumulation of TuYVs81 RNA in systemic leaves in the indicated genotypes at 20 dpi (n=5 plants) measured by RNA gel blot. “@” indicates hybridization with DNA probe and loading control is obtained by staining the membrane with methylene blue (MB). **(H.)** Representative individuals infected with TuYVs81 displaying systemic vein yellowing. **(I.)** Quantification of viral RNA signal in **E** and **G** relative to Col-0 and normalized to MB signal. **(J.)** vsiRNA abundance in total RNA, AGO1 and AGO2 immunoprecipitates in TuYVs81 and TRV-PDS infected leaves. “@” indicates hybridization with DNA probe or use of a specific antibody for immunoprecipitation. **(K.)** Global quantification of 21-nt to 24-nt small RNA reads aligned to the reference Arabidopsis genome and TuYVs81 genome per functional categories (araport11), expressed as (RPM) reads per million ([category count * 1.000.000]/library size). Libraries were obtained from total RNA and AGO1 IP in mock-inoculated (mock) or infected (TuYVs81) systemic leaves at 16 dpi (n= 7 or 8 individual plants per replicate). R1 = replicate 1, R2 = replicate 2. **(L.)** Distribution of TuYVs81-derived sRNA reads (20-nt to 25-nt) along the TuYVs81 genome in Col-0 total RNA and AGO1 IP replicate 1 (R1), with MISIS. Bars indicate the position of the 5’ (+ strand) and 3’ (-strand) extremity of each mapped sRNA. Y-axis represents read counts, and each size category is represented in the indicated color.

Other AGO proteins have been shown to be involved in the silencing of several RNA viruses, sometimes in combinations (Carbonell & Carrington, 2015). We therefore tested the contribution of different AGOs to TuYV silencing by infecting single or multiple knock out mutants for *AGO5*, *AGO10*, *AGO2* and *AGO7*, in addition to *ago1-27*, which is a missense allele. Quantification of the viral RNA in systemic leaves revealed that only the *ago1-57* mutation caused enhanced accumulation of TuYV (**Figure S1A**), further demonstrating the importance of AGO1 in TuYV antiviral defense. Here too, the *ago1-27* mutant did not exhibit any difference, neither did the combinations that contained that allele.

Because AGO2 has been described in several studies as the main antiviral AGO (Harvey *et al*, 2011; Jaubert *et al*, 2011; Garcia-Ruiz *et al*, 2015) or to function in tandem with AGO1 (Ma *et al*, 2015; Wang *et al*, 2011), we introgressed the *ago2-1* mutation into the *ago1-57* background and tested for antiviral performance against several well-studied RNA viruses. Plants were inoculated with an infectious clone of turnip mosaic virus (TuMV) containing a GFP reporter and a point mutation inactivating the VSR activity of HC-Pro, referred to as TuMV-AS9-GFP (Garcia-Ruiz *et al*, 2010). While GFP was clearly visible in inoculated leaves of *ago2-1* plants, systemic spread of the virus did not occur (**Figure 1C and D**). By contrast, *ago1-57* plants allowed limited spread of TuMV-AS9-GFP into the systemic leaves while the *ago1-57/ago2-1* combination allowed for strong GFP signal in the vasculature with elevated amounts of GFP protein. This reveals a previously unreported role for AGO1 in defense against TuMV and shows that both fully functional AGO1 and AGO2 are needed to mount efficient defense against TuMV-AS9-GFP. We then infected the same genotypes with tobacco rattle virus (TRV) carrying a *PHYTOENE DESATURASE* (PDS) fragment, referred to as TRV-PDS (Ratcliff *et al*, 2001), which triggers potent VIGS against this gene resulting in a bleached leaf phenotype. Both AGO1 and AGO2 have been reported as necessary for efficient antiviral silencing during TRV infection (Ma *et al*, 2015). Accordingly, we observed increased viral RNA and impaired PDS silencing in the double *ago1-57/ago2-1* mutant, while single mutants behaved identically to Col-0 (**Figure 1E and F**). In contrast, *ago2-1* single mutation did not affect TuYVs81 RNA accumulation, while the single *ago1-57* and the double mutant showed enhanced accumulation and an identical impairment in VIGS (**Figure 1G and I**). This suggests that, as opposed to TuMV and TRV, efficient antiviral silencing of TuYV by the plant does not rely on the coordinated action of AGO1 and AGO2, but rather that AGO1 alone is necessary to mount an efficient response.

Since AGO proteins can bind overlapping cohorts of sRNA and act redundantly on an RNA target, impairment of a particular AGO can result in compensatory loading by another AGO. We reasoned that if AGO2 can act as a surrogate to AGO1, then compromising *AGO1* function should lead to an increase in AGO2 loading with vsiRNA. We therefore performed immunoprecipitations of both AGO1 and AGO2 (**Figure S1B**) and analyzed the associated sRNA in plants systemically infected by TRV-PDS and TuYVs81. TRV-PDS-derived vsiRNA where loaded into AGO1 and AGO2, and in the *ago1- 57* background we observed increased loading into AGO2 (**Figure 1J**). TuYVs81 vsiRNA were found to be predominantly associated with AGO1, with increased production of vsiRNA in *ago1-57* plants likely leading to increased loading of AGO1-57. More importantly, little or no compensatory loading of AGO2 was observed, further establishing the pivotal role of AGO1 in TuYVs81 antiviral defense. We also compared loading abilities of AGO1-57 and AGO1-27 (**Figure S1C**) and found increased AGO2 loading with vsiRNA only in *ago1-27*, concomitant with impaired loading of the AGO1-27 protein of both vsiRNA and miRNA. This can be explained by the increased AGO2 protein level observed in *ago1-27* (**Figure 1SD**) caused by lessened miR403-AGO1 complex assembly (**Figure S1C**), a feature not observed in the *ago1-57* mutant. Like for TuYVs81, we also found that loading of AGO1 and AGO2 with *SUL* siRNA was identical in WT and *ago1-57* plants (**Figure S1E and F**), indicating that the mutant protein is not affected in siRNA loading, as we have previously shown (Derrien *et al*, 2018). Taken together, these results suggest that *ago1-57* presents a unique opportunity for the study of the antiviral role of AGO1 *in planta*, since it selectively impacts siRNA guided pathways and only marginally affects miRNA-RISC regulation.

Next, we performed sRNA deep sequencing of total RNA and AGO1 IPs from mock-inoculated and TuYVs81 infected Col-0 and *ago1-57* plants (**Figure S2A**). After verifying successful immunoprecipitation of AGO1 in all replicates and samples (**Figure S2B**), sRNA reads of 18-26nt were mapped to the reference Arabidopsis genome (Cheng *et al*, 2017) and the TuYVs81 genome and normalized 21-24nt reads mapping to the different categories are shown in **Figure 1K**. As expected, AGO1 IPs of mock samples contained abundant reads mapping to miRNA, that were mostly of 21-nt (**Figure S2C**). In infected samples, a large amount of vsiRNA were recovered from both the total RNA and the AGO1 IP, indicating that a large amount of the produced vsiRNA are channelled towards AGO1, as observed by northern blot. These vsiRNA were found to originate from both strands and to completely cover the TuYVs81 genome, with some hotspots found across the replicates and samples (**Figure 1L** **and S2D**). TuYVs81-mappers were enriched in AGO1 IP with some reaching a read count of over 400.000 in AGO1-57 IPs. Intriguingly, most reads mapping to the TuYVs81 genome were of 22-nt (**Figure 1L** **and S2C**) suggesting that DCL2, among the four Arabidopsis DCL proteins, is predominantly responsible for TuYVs81 processing. We also observed an impaired 5’U bias in AGO1 IPs from the *ago1-57* plants (**Figure S2E**). This is explained by the molecular phenotype of the allele, which retains both the guide strand and the passenger strand (Derrien *et al*, 2018), the latter having a 5’ extremity that can be any nucleotide, artificially raising the amount of 5’ C, G and A reads. Altogether our results show that AGO1 plays a key role in orchestrating antiviral defense against TuYV, by loading a vast population of vsiRNA which are mostly 22nt-long.

### Non-degradable AGO1-57 impairs establishment of systemic TuYVs81 infection

After numerous TuYVs81 infections using agro-inoculation we noticed that about only half of the initially inoculated *ago1-57* plants developed VIGS, in contrast to Col-0 inoculated plants that showed up to 100% systemic infection rate (**Figure S3A**). Since *ago1-57* is impaired in VIGS, we chose to monitor the presence of the viral readthrough protein (RT) in young leaves of the inoculated plant population during the progression of infection. This showed a similar kinetic to that of VIGS, with only 43,3% of the *ago1-57* plants exhibiting infection, while infection of *ago1-57* with TuMV-GFP and TRV-PDS reached 100% (**Figure S3A**). We hypothesized that this observation could be explained by the evasion of AGO1-57 from P0-mediated degradation or alternatively, that the mutation in *AGO1* leads to constitutive activation of defense-related genes leading to enhanced resistance, as observed for TRV-infected *ago1-27* (Ma *et al*, 2015), and regulated by miRNA and phasiRNA in multiple plant species (Li *et al*, 2012; Shivaprasad *et al*, 2012; Deng *et al*, 2018; López-Márquez *et al*, 2020). To discriminate between these two hypotheses, we performed similar infection kinetics in the three *ago1* hypomorphic mutants used previously. We found that only the *ago1-57* plant population exhibited a 50% loss of systemic infection by TuYVs81, while *ago1-27* and *ago1-38* both reached 100% infection, albeit with very dissimilar kinetics (**Figure 2A and B**). Importantly, this loss was not observed during establishment of TuMV-GFP infection (**Figure 2C and D**), indicating that it is unique to the *ago1-57*- TuYVs81 interaction, and likely results from the undegradable nature of the mutant protein.

**Figure 2:**
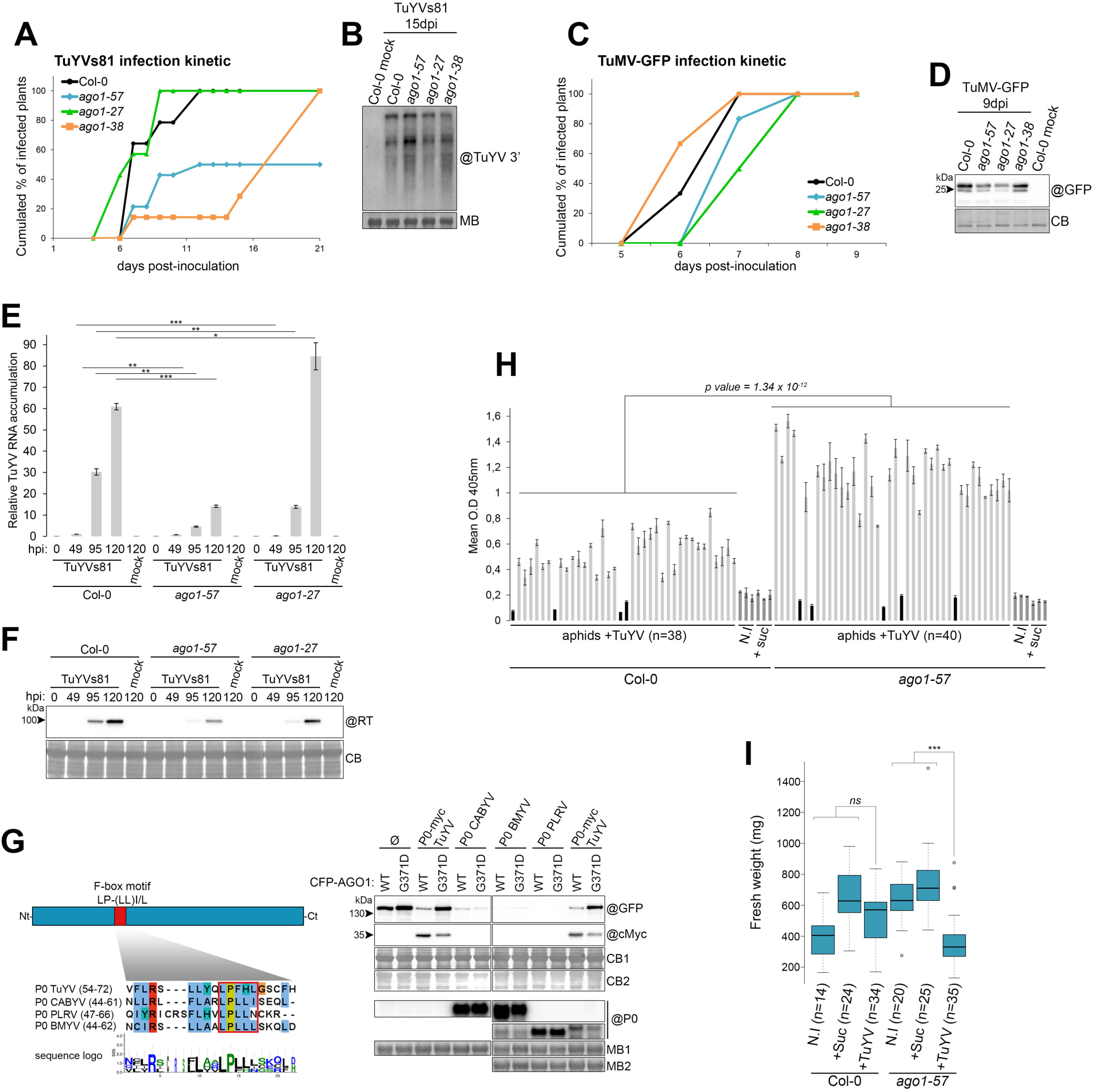
*ago1-57* uniquely affects systemic movement of TuYVs81 due to delayed viral RNA accumulation after agrobacterium-mediated inoculation but fails to provide any protective effect in vector-mediated infection. **(A.)** Kinetic of systemic TuYVs81 infection in Col-0, *ago1-57*, *ago1-27* and *ago1-38* represented as the cumulated percentage of infected plants in the inoculated population (n=14 individuals per genotype). To avoid any confounding effect introduced by VIGS deficiency in the mutants, infected individuals were scored by the detection of the TuYVs81 readthrough protein (RT, ORF5) in leaf patch from young systemic leaves at 6, 7, 9, 12, 15 and 21 dpi using either western or dot blot. Plants that exhibit systemic VIGS in between these sampling times are also counted as infected at the time point at which the VIGS was first observed. **(B.)** TuYVs81 viral RNA abundance in systemic leaves of the indicated mutants at 15 dpi measured by RNA blot. Each sample represents a mix of leaf patches for all infected individuals at that time point in the kinetic. “@” indicates hybridization with DNA probe and loading control is obtained by staining the membrane with methylene blue (MB). **(C.)** Kinetic of systemic TuMV-GFP infection in Col-0, *ago1*-*57*, *ago1-27* and *ago1-38* represented as the cumulated percentage of infected plants in the inoculated population (n=6 individuals per genotype). Infected individuals were scored by the detection of GFP in systemic leaves at the indicated day. **(D.)** Detection of TuMV-GFP in systemic leaves of the indicated mutants at 9 dpi measured by immunoblot. “@” indicates hybridization with GFP antibody, loading control is obtained by post-staining the membrane with Coomassie blue (CB). (E.) *ago1-57* displays delayed viral RNA accumulation after inoculation. Measurement of TuYVs81 RNA in inoculated leaves at 0, 49, 95, and 120 hours post infiltration (hpi) measured by RT-qPCR in the indicated genetic backgrounds represented as bar graph relative to Col-0 49hpi. Represented values are means of technical triplicates, error bars represent the SEM. For each time point, 5 infiltrated leaves from 4 individuals were harvested for each genotype. *p<0.05, **p<0.01, ***p<0.001 with Student’s t-test, one-tailed, paired. **(F.)** Accumulation of TuYVs81 readthrough protein (RT) at 0, 49, 95, and 120 hours post infiltration (hpi) measured by immunoblot. Samples are from the same tissues as in **E**. “@” indicates hybridization with RT antibody, loading control is obtained by post-staining the membrane with Coomassie blue (CB). **(G.)** G371D mutation does not confer undegradability to AGO1 in presence of P0 from diverse polerovirus species. Left panel: Schematic representation of the P0 protein, with the region containing the F-box motif highlighted in red and the corresponding alignment of P0 from turnip yellows virus (TuYV), cucurbit aphid-borne yellows virus (CABYV), beet mild yellowing virus (BMYV) and potato leafroll virus (PLRV) shown below. The minimal F-box motif is boxed in red, and the sequence logo for the alignment is shown below. Right panel: AGO1 degradation test in *N. benthamiana*. Both versions of CFP-AGO1 (WT or G371D) were expressed either without (ø) or with the indicated P0 proteins in two separate leaves and an equivalent amount of leaf patches were collected at 4 dpi. All infiltrated patches contain P19. Fusion protein levels were assessed by immunoblot on two different membranes (GFP corresponds to Coomassie stain CB1 and cMyc to CB2) and “@” indicates hybridization with the corresponding specific antibody. Expression of the untagged P0 constructs was verified by RNA blot using DNA probes specific for each sequence (@ P0) and equal loading was assessed by staining the corresponding membrane with methylene blue (MB1 and MB2). **(H.)** Detection and quantification of aphid-transmitted WT TuYV virions in systemic leaves of Col-0 or *ago1-57* individual plants. Eighteen-day-old plants were individually challenged with two *Myzus persicae* fed on either 20% sucrose solution (+Suc) or 20% sucrose solution containing 67mg/ml TuYV virions (+TuYV) or alternatively were left untreated (N.I). Each bar represents the mean O.D at 405nm (technical triplicate measurements) for a single individual within the considered category, and error bars represent SD. Individuals for which the O.D was ≤ to those of the N.I and +Suc control plants are considered as non-infected and are colored in black. Difference between the Col-0 and *ago1-57* populations is statistically different (Kruskal and Wallis test). **(I.)** Fresh weight measurement of all the analyzed plants in Figure 2H after removal of the non-infected plants, expressed in mg. ***p < 0.001 (two-way ANOVA followed by Tukey honest significant differences test to compare both genotypes and treatments).

P0-less mutants of TuYV show lower viral replication in protoplasts (Ziegler-Graff *et al*, 1996), and PLRV P0 is essential for viral multiplication in inoculated tissues (Sadowy *et al*, 2001), suggesting that suppression of RNA silencing is important during early steps of infections. Since AGO1-57 is resistant to P0-mediated degradation, it should exert a similar effect to the loss of viral P0 during the local establishment of infection, before systemic movement can be achieved. We therefore assayed viral RNA abundance in Arabidopsis leaves agro-inoculated with TuYVs81 in *ago1-57* compared to Col-0 and *ago1-27*. In all experiments, *ago1-57* leaves contained significantly less viral RNA than Col-0 at any given time (**Figure 2E** **and S3B**) and showed lower RT protein accumulation (**Figure 2F**). This slower build-up of infection in *ago1-57* was not shared by the *ago1-27* mutant, that is sensitive to P0-mediated degradation (Derrien *et al*, 2018), and is thus the consequence of resistance to P0.

Because P0 proteins from different poleroviruses have low sequence identity (Mayo & Ziegler-Graff, 1996) but seem to all contain the minimal F-box consensus motif (LPxxL/I), we tested if the G371D mutation enables AGO1 evasion from a range of P0 proteins. WT and G371D CFP-AGO1 were co-infiltrated with P0 proteins from TuYV, PLRV, cucurbit aphid-borne yellows virus (CABYV) and beet mild yellowing virus (BMYV), that all contain the F-box motif, and CFP-AGO1 degradation was assayed as previously described (Baumberger *et al*, 2007). Surprisingly, while the G371D mutation conferred protection from P0^Tu^ degradation to AGO1, the mutation failed to prevent degradation by the three other P0 proteins tested (**Figure 2G**). This highlights the existence of different modes of AGO1 recognition employed by viral P0s and indicates that the strategy employed by TuYV to target AGO1 is not a conserved feature of poleroviruses.

Finally, we assayed the performance of *ago1-57* when TuYV is delivered by its natural aphid vector, that directly injects virions into phloem cells. When infection was monitored 25 days after transmission using DAS-ELISA for each individual, we found that the number of plants that scored positive for the presence of the virus was comparable between Col-0 (89,5%) and *ago1-57* (87,5%) (**Figure 2H**). Not only did the *ago1-57* plants not display the reduced systemic infection as observed for agro-inoculation, but all the plants contained significantly more virion than their WT counterparts. Intriguingly, while TuYV infection normally leads to a symptomless infection in Arabidopsis, we observed a statistically significant reduction in fresh weight for the TuYV-infected *ago1-57* plants (**Figure 2I**), suggesting that impairment of AGO1 function leads to symptomatic infection. Taken together, our analysis shows that evasion of degradation by the mutant AGO1 causes delay in the establishment of local infection, leading to a decrease in systemic infection. This apparent resistance phenotype is overruled by direct inoculation into phloem cells.

### TuYV RNA is mostly processed by DCL2, but both DCL2 and DCL4 are necessary to mount an effective antiviral defense

To test the contribution of DCLs to vsiRNA production, we performed northern blots from tissues systemically infected by TuYV, in single and multiple mutants containing the *dcl2-1* or *dcl2-5*, *dcl4-2*, *dcl3-1* and *ago1-57* alleles. In *dcl2-1* infected plants, the main 22-nt TuYV vsiRNA band present in wild-type was replaced by a 21-nt band, indicating that the bulk of vsiRNA is indeed the product of DCL2, while loss of DCL4 did not affect the main vsiRNA population (**Figure 3A**). Loss of both DCL2 and DCL4 resulted in the sole accumulation of a 24-nt signal that is otherwise minimal in WT plants, while the *dcl2-5/dcl3-1* combination led to a single 21-nt signal. Loss of 22-nt vsiRNA in *dcl2-1* was consistently accompanied by lesser silencing signal spread, a feature that was not observed in the *dcl4-2* plant (**Figure 3B** **and S4A**), mirroring their relative contribution to vsiRNA production in the phloem cells. VIGS signal was abolished in *dcl2-1/dcl4-2* plants, while *dcl2-5/dcl3-1* were identical to single *dcl2-1*, indicating that the residual 24-nt do not constitute a mobile silencing signal sufficient to initiate VIGS. Identical vsiRNA patterns were observed when the *ago1-57* mutation was added to each combination (**Figure 3A**). Combining *dcl2-1* an *ago1-57*, that both exhibit weak VIGS, led to a complete loss of cell-to-cell silencing spread, which was not the case in *dcl4-2/ago1-57*, that produces abundant vsiRNA. We measured TuYV RNA content in the same tissues and found that, like *ago1*-*57*, single *dcl2-1* and *dcl4-2* mutations lead to a two-fold increase of TuYV RNA in systemic tissues, that was not increased when introduced in the *ago1-57* background (**Figure 3C**). Similarly, the loss of both *DCL2* and *DCL3* resulted in a two-fold increase while the double *dcl2-1/dcl4-2* lead to a dramatic increase in the amount of TuYV RNA accompanied by severe symptoms (**Figure 3B**). Intriguingly, the triple mutant combination lead to an even greater viral RNA accumulation, supporting that the 24-nt vsiRNA are able to exert antiviral activity through AGO1.

**Figure 3:**
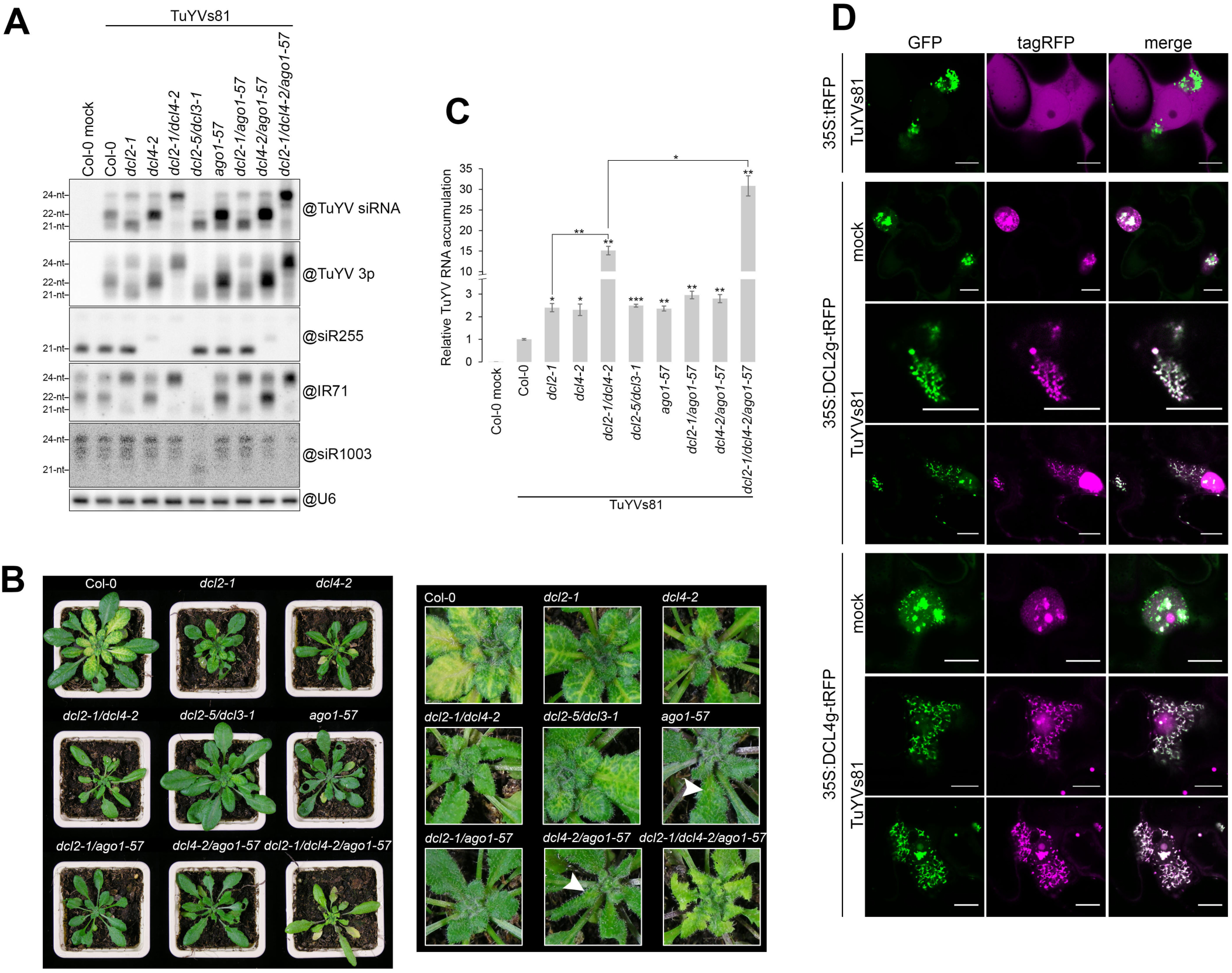
TuYvs81 RNA is mainly processed by DCL2 into 22-nt vsiRNA, yet both DCL2 and DCL4 are necessary to efficiently silence the virus. **(A.)** Analysis of vsiRNA size in infected TuYVs81 infected plants at 17 dpi using RNA gel blot. Most of the vsiRNA populations is of 22-nt and is lost in a *dcl2-1* single or combination mutant. “@” indicates hybridization with the indicated DNA probe. Endogenous siRNA are used to control for the proper identity of the *dcl* mutants and U6 signal is the loading control. **(B.)** Representative image of the infected genotypes analyzed in **A**. Left panel: whole plant view. Right panel: inset of young systemic leaves. White arrow indicate very feint vein yellowing. **(C.)** TuYVs81 viral RNA abundance in systemic leaves of the indicated mutants at 17 dpi measured by RT-qPCR. Levels are displayed relative to infected Col-0. Each sample represents a pool of several individuals from the indicated genotype. Only individuals that scored positive for the presence of systemic TuYVs81 (via detection of the RT protein in leaf patches) were harvested. p value above each sample is for pairwise comparison of the sample to Col-0. *p<0.05, **p<0.01, ***p<0.001 with Student’s t-test, two-tailed, unequal variance. **(D.)** Both DCL2 and DCL4 are redirected to cytosolic viral replication complexes (VRC) during TuYVs81 infection and colocalize with viral double stranded RNA. Representative single plane confocal images of transiently expressed 35S:tRFP, 35S:DCL2genomic-tRFP and 35S:DCL4genomic-tRFP with (TuYVs81) or without (mock) the virus in leaves of transgenic *N. benthamiana* stably expressing the double-stranded RNA-binding B2-GFP protein. Observations are from leaf discs of 3 to 5 days post-infiltration. Inset scale bar is 10 µm. See also **Figure S5** for additional images.

In order to test the contribution of secondary siRNA to antiviral silencing of TuYVs81, we infected loss of function mutants for *RDR6* and *SGS3*, that are both required for production of dsRNA and silencing signal amplification from AGO1 cleavage products, a mechanism that participates in antiviral silencing for some plant RNA viruses (Mourrain *et al*, 2000; Qu *et al*, 2008; Wang *et al*, 2010; Garcia-Ruiz *et al*, 2010). This revealed that neither VIGS (**Figure S4A**), vsiRNA profiles (**Figure S4B**) nor TuYV RNA accumulation (**Figure S4C**) were affected in those mutants. The same mutants presented systemic infection kinetics that were comparable to those of WT and *dcl* mutant plants (**Figure S4D**). This shows that silencing signal amplification via RDR6/SGS3 is dispensable for antiviral silencing during infection by TuYVs81. Note that this does not rule out production of secondary vsiRNA from the viral RNA after AGO1 slicing.

Since both DCL2 and DCL4 have antiviral activity against TuYVs81, but DCL2 rather than DCL4 is the major contributor of vsiRNA, we sought to establish their localization relative to viral replication complexes (VRCs). We took advantage of the recently established stable *N. benthamiana* line expressing the eGFP-tagged dsRNA binding domain of flock house virus (FHV) B2 protein (B2-GFP) that allows *in vivo* visualization of dsRNA, the minimal component of the VRCs (Monsion *et al*, 2018). Consistent with their ability to produce vsiRNA from double stranded TuYV RNA, both AtDCL2-tRFP and AtDCL4-tRFP were found to equally re-localize from the nucleus to the B2-GFP labelled VRCs (**Figure 3D** **and S5**), which was not the case for the nucleo-cytoplasmic tRFP control. In summary, our results show that both DCL2 and DCL4 are directed to VRCs upon infection to initiate antiviral defense against TuYVs81, but DCL2 appears to be the major source of vsiRNA in systemically infected tissues.

### DCL4 and DCL2 operate normally in TuYV-infected vasculature

To characterize more specifically the relative roles of DCL2 and DCL4 in infected vasculature, we employed MeSelect (mechanical separation of leaf compound tissues), a technique which enables isolation of vascular bundle cells (Svozil *et al*, 2015). As expected, TuYVs81 RNA was enriched when compared to whole leaf extracts (**Figure 4A** **and S6A**), and the loss of *DCL2* lead to an increased accumulation of TuYVs81 RNA, as observed in whole leaves (**Figure 3C**). Vascular vsiRNA were 22-nt in size (**Figure 4B** **and S6B**), confirming that their presence in whole leaf arises from the processing of TuYVs81 RNA by DCL2 in the vasculature. We also observed a smearing pattern above the 22-nt signal only in the enriched vasculature (**Figure S6B**) suggesting persistence of precursor RNA species/replication intermediates.

**Figure 4:**
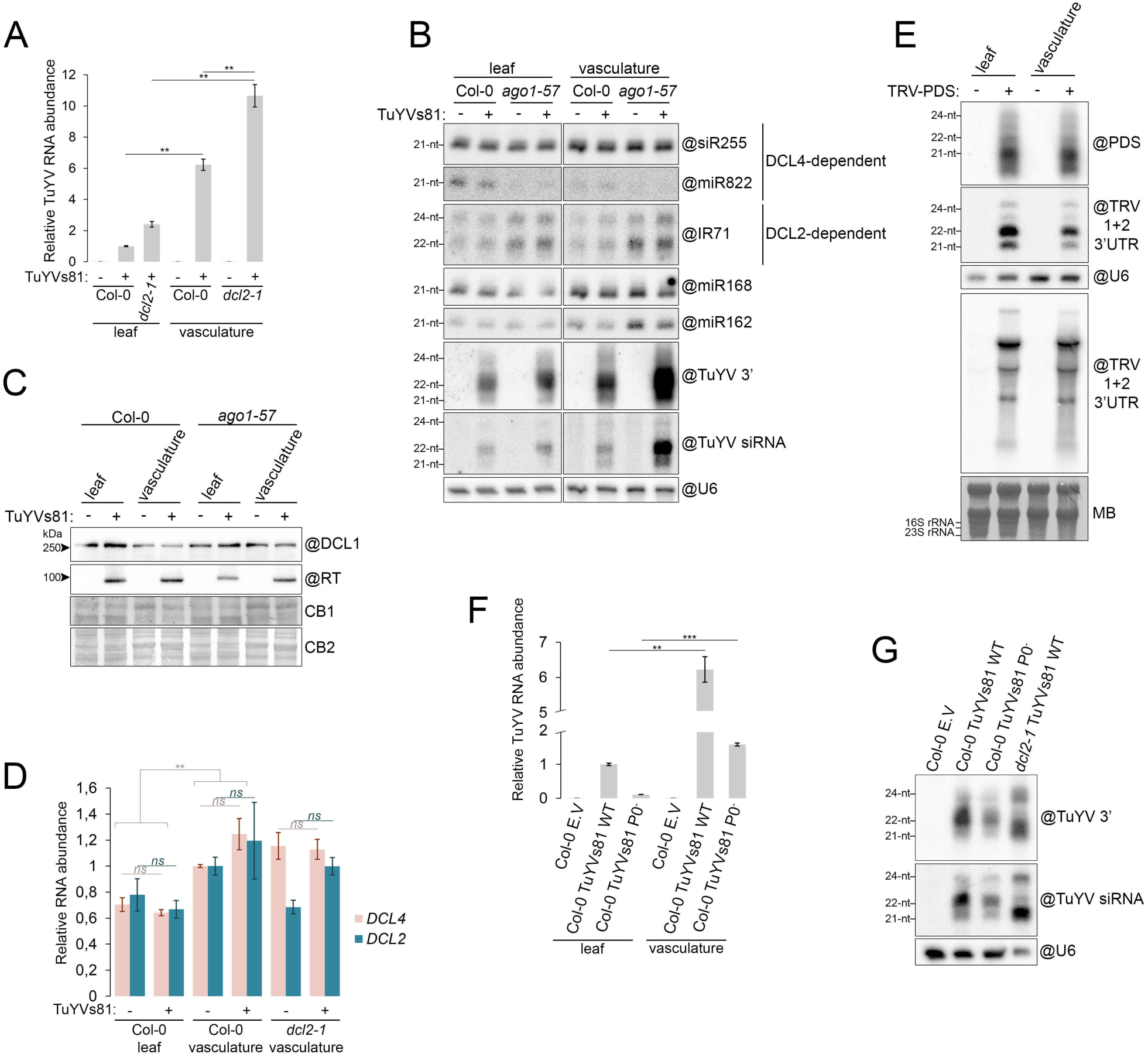
DCL2 processing is not a consequence of detectable viral manipulation in the infected vasculature. **(A.)** Measurement of TuYVs81 RNA in systemic whole leaves of Col-0 and *dcl2-1* plants (n=8 to 10 individuals) or enriched vascular bundles of the equivalent plants (n=20 leaves) at 17 dpi by RT-qPCR represented as bar graph relative to infected Col-0 leaves. Represented values are means of technical triplicates, error bars represent the SEM. **p<0.01, with Student’s t-test, one-tailed, paired. **(B.)** Analysis of sRNA abundance in total RNA extracted from the whole leaf or the vasculature of mock (-) or TuYVs81 infected (+) Col-0 and *ago1-57* plants by RNA gel blot. For leaf tissue, a mixture of infected leaves from several individual was used (n=8) and enriched vascular bundles (n=24 leaves) were obtained from the equivalent plants at 16 dpi. “@” indicates hybridization with DNA probe and U6 signal is the loading control. **(C.)**. Detection of the DCL1 and viral RT proteins in total protein extracted from the whole leaf or the vasculature of mock (-) or TuYVs81 infected (+) Col-0 and *ago1-57* plants by immunoblot. Samples were prepared from the same material as in **B**. “@” indicates hybridization antibodies, loading control is obtained by post-staining the membrane with Coomassie blue (DCL1 corresponds to Coomassie stain CB1 and RT to CB2). **(D.)** Measurement of *DCL2* and *DCL4* mRNA abundance in the same samples as **A** by RT-qPCR, represented as bar graph relative to - Col-0 vasculatures. Note that loss-of-function allele *dcl2-1* contains WT level of DCL2 messenger RNA. Represented values are means of technical triplicates, error bars represent the SEM. **p<0.01, *ns* p>0.05 with Student’s t-test, two-tailed, paired. See **Figure S6C** for an additional experiment. **(E.)** Analysis of TRV-PDS (+) infected whole leaves and vasculature of Col-0 plants (n=9 individuals, 8 leaves for vasculature) at 16 dpi by RNA gel blot. Top panel: sRNA blot shows equivalent profiles of vsiRNA made against the PDS insert that are mostly processed by DCL4 into 21-nt. An oligoprobe recognizing the conserved 3’ of RNA1 and 2 of TRV shows that this portion of the viral RNA is mostly processed by DCL2 into 22-nt, irrespective of the tissue. Bottom panel: TRV RNA is present in both tissue types and is not enriched in the vasculature. “@” indicates hybridization with DNA probe, U6 signal and methylene blue staining are the loading control. Note that chloroplastic 16S and 23S rRNA are only visible in whole leaf. **(F.)** Measurement of TuYVs81 RNA in systemic Col-0 whole leaves (n=8 to 10 individuals) or enriched vascular bundles of the equivalent plants (n=20 leaves) at 17 dpi by RT-qPCR represented as bar graph relative to infected leaves. Plants were inoculated either with the empty vector (mock), with the WT TuYVs81 or with a P0-less mutant. Represented values are means of technical triplicates, error bars represent the SEM. **p<0.01, ***p<0.001 with Student’s t-test, one-tailed, paired. **(G.)** Analysis of vsiRNA profile from whole leaf of Col-0 plants infected with either WT TuYVs81 or the P0-less TuYVs81 by RNA gel blot. Samples are the same as in **F**, infected *dcl2-1* is used a control for the absence of 22-nt. “@” indicates hybridization with DNA probe and U6 signal is the loading control.

Similarly to TuYV, turnip crinkle virus (TCV) is also processed mainly into 22-nt-long vsiRNA in Arabidopsis, its VSR P38 inhibiting the AGO1-miR162 regulation of DCL1 (Azevedo *et al*, 2010) which itself represses DCL4 and DCL3 via an unknown mechanism (Qu *et al*, 2008). To assess whether this is the case during TuYV infection, we monitored the abundance of miR162 and DCL1 in infected vasculature. We observed a mild decrease in the miR162 signal in both Col-0 and *ago1-57* infected vasculature (**Figure 4B**), but no appreciable difference in the abundance of the DCL1 protein in the same tissue (**Figure 4C**), ruling out a mechanism similar to TCV regulation of DCL1 in the case of TuYV. Measurement of *DCL2* and *DCL4* mRNA abundance showed that neither are affected by the presence of the virus in the vasculature (**Figure 4D** **and S6C**). To test if the activity rather than the expression of DCL4 is impaired in the presence of TuYVs81, we monitored the accumulation of DCL4-dependent endogenous sRNA. While miR822 is barely produced in the vasculature, siR255 (TAS1) abundance was unperturbed by the presence of the virus in the vasculature (**Figure 4B**), suggesting that DCL4 functionality is intact. Similarly, IR71 22-nt siRNA are unaffected by the virus, demonstrating that DCL2 is functioning normally. Because DCL4 and DCL2 usually exhibit hierarchical activity towards transgenes and RNA viruses, with DCL4 being the prime source of siRNA over DCL2, we wondered whether in the vasculature this hierarchy could be inverted. We thus compared sRNA profiles in whole leaf to those in vasculature recovered from TRV-PDS infected plants, an RNA virus that is primarily processed into 21-nt by DCL4 in WT plants (Deleris *et al*, 2006) and found that the vsiRNA profiles were identical both for the PDS reporter and the 3’ of RNA1 and RNA2 (**Figure 4E**), demonstrating that the vascular DCL hierarchy is identical to that of the leaves.

Collectively, these results show that processing of TuYVs81 by DCL2 is not a consequence of attenuated DCL4 expression and/or activity, nor of increased expression or overall activity of DCL2 in infected vasculature. Neither DCL2 nor DCL4 show vasculature-specific antiviral activity when challenged with an unrelated RNA virus, suggesting in turn manipulation of the DCL balance by the TuYV. To test whether P0 could affect the balance between DCL2 and DCL4 in TuYV processing, we compared the vsiRNA profiles in systemic tissues infected with either WT TuYVs81 or TuYVs81P0^-^.

As observed previously in whole leaves (**Figure 1A**), TuYVs81P0^-^ was less abundant than its WT counterpart in the isolated vasculatures (**Figure 4F**) but was nonetheless enriched comparatively to the leaf. Although the signal was weaker, TuYVs81P0^-^ infection still resulted in a major 22-nt signal (**Figure 4G**), indicating that P0 is not responsible for preponderant processing of TuYV by DCL2. The factor responsible for this phenomenon remains to be identified.

### Vascular AGO1 is sensitive to P0 but is unexpectedly stabilized in presence of TuYV

Next, we took advantage of the MeSelect method to observe viral P0-driven degradation of AGO1, a feature that is obscured in whole leaf tissues, and habitually observed with over-expression of P0 in Arabidopsis (Bortolamiol *et al*, 2007; Derrien *et al*, 2018; Michaeli *et al*, 2019) or in transient co-expression experiments (**Figure 2G**). We therefore separated vasculature from Col-0 and *ago1-57* infected plants, with both genotypes presenting a significant enrichment of TuYVs81 RNA in the vasculature, and a 3.5-fold enrichment in *ago1-57* compared to Col-0, as expected (**Figure 5A**). Strikingly, rather than a depletion, we observed an increased accumulation of vascular AGO1 protein in the presence of the virus, irrespective of the background used (**Figure 5B, S7A and S7B**). This enrichment relative to the mock inoculated vasculature was very robust and observed in vasculatures collected at different days post inoculation as quantified in **Figure 5C** (n=5 separate infection experiments), and was not observed for either AGO2 and AGO4 (**Figure 5B** **and S7B**).

**Figure 5:**
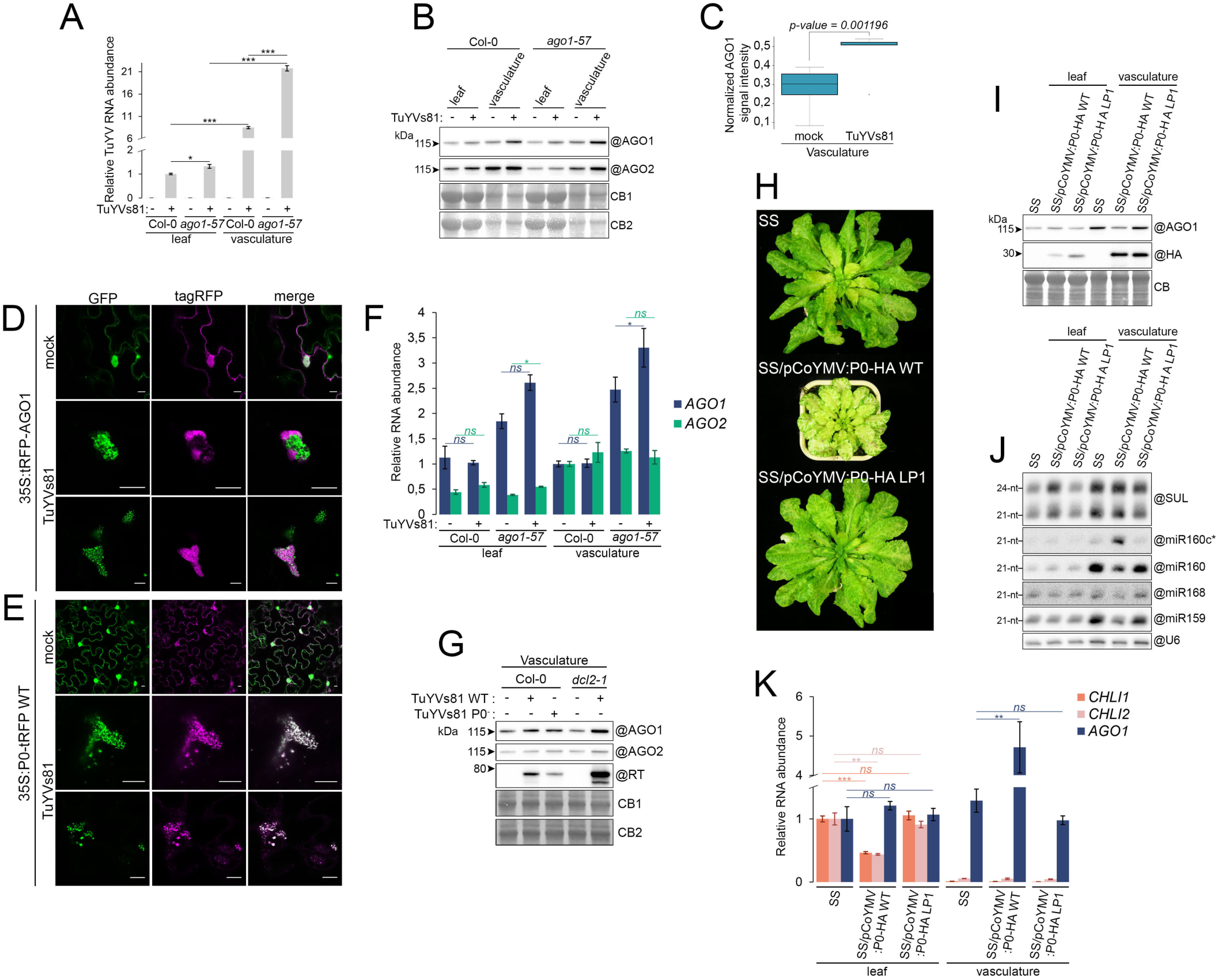
Vascular AGO1 is post-translationally stabilized in presence of TuYVs81, despite the ability of a transgenic, companion cell restricted P0 to degrade AGO1. **(A.)** Measurement of TuYVs81 RNA in systemic whole leaves of Col-0 and *ago1-57* plants (n=8) or enriched vascular bundles of the equivalent plants (n=24 leaves) at 16 dpi by RT-qPCR represented as bar graph relative to infected Col-0 leaves. Represented values are means of technical triplicates, error bars represent the SEM. *p<0.05, ***p<0.001 with Student’s t-test, one-tailed, paired between tissues, unequal variance between genotypes. **(B.)** Representative immunoblot of AGO1 and AGO2 accumulation in systemic whole leaves (n=12 individuals) and in enriched vascular bundles of the equivalent plants (n=18 leaves) at 21 dpi in Col-0 and *ago1-57*, in the absence (-) or presence (+) of TuYVs81. “@” indicates hybridization with the indicated antibodies, and loading control is obtained by post-staining the membrane with Coomassie blue (AGO1 on CB1 and AGO2 on CB2). **(C.)** Quantification of the AGO1 signal normalized to total protein signal (Coomassie blue stain, whole lane) in Col-0 vasculatures without (mock) or in the presence of TuYVs81. Collected values represent 5 biological replicates in which vasculatures were enriched at either 15, 16, 17 or 21 dpi from different infection experiments. p-value = 0.001196 with Student’s t-test, one-tailed, paired. **(D)** Representative single plane confocal images of transiently expressed 35S:tRFP-AGO1 with (TuYVs81) or without (mock) the virus in leaves of transgenic *N. benthamiana* stably expressing the double-stranded RNA-binding B2-GFP protein. Observations are from leaf discs at 3 dpi. Inset scale bar is 10 µm. See also **Figure S8A** for additional images. **(E.)** P0 colocalizes with viral double stranded RNA. Representative confocal images of transiently expressed 35S:P0-tRFPwith (TuYVs81) or without (mock) the virus in leaves of transgenic *N. benthamiana* stably expressing the double-stranded RNA-binding B2-GFP protein. Observations are from leaf discs at 3 dpi. Inset scale bar is 10 µm. See also **Figure S8B** for additional images. **(F.)** Levels of *AGO1* and *AGO2* mRNA are not significantly affected by the presence of TuYVs81 in the plant vasculature. mRNA abundance in the same samples as **A** by RT-qPCR, represented as bar graph relative to Col-0 vasculatures. Represented values are means of technical triplicates, error bars represent the SEM. *p<0.05, *ns* p>0.05 with Student’s t-test, two-tailed, paired. See **Figure S8G** for an additional experiment. **(G.)**Analysis of AGO1 and AGO2 protein level in enriched vascular bundles of Col-0 or *dcl2-1* plants (n=20 leaves from 8 to 10 individuals per genotype and treatment) at 17 dpi after inoculation with E.V (-), TuYVs81 WT or TuYVs81 P0^-^. “@” indicates hybridization with the indicated antibodies, and loading control is obtained by post-staining the membrane with Coomassie blue (AGO1 on CB1 and AGO2 on CB2). **(H.)** Persistent companion cell expression of P0 enhances spreading of *SUL* siRNA. Image of representative individual adult plants of WT, pCoYMV:P0-HA WT and LP1 in the SS background. **(I.)** Vascular AGO1 is degraded in presence of WT P0-HA. Immunoblot of AGO1 and P0-HA accumulation in whole leaves (n=4 individuals) and in enriched vascular bundles of the equivalent plants (n=10 to 16 leaves) in the indicated genotypes. Plants are the same as in **H**. “@” indicates hybridization with the indicated antibodies, and loading control is obtained by post-staining the membrane with Coomassie blue (CB). **(J.)** Analysis of sRNA abundance in total RNA extracted from the whole leaf or vasculature by RNA gel blot. RNA were obtained from the same samples as in **I**. “@” indicates hybridization with DNA probe and U6 signal is the loading control **(K.)** Degradation of AGO1 by vascular P0 leads to increased abundance of the *AGO1* mRNA in the vasculature and decrease of the total amount of *CHLI1* and *CHLI2* mRNA. mRNA abundance in the same samples as **L** by RT-qPCR, represented as bar graph relative to SS leaf. Represented values are means of technical triplicates, error bars represent the SEM. ***p<0.001, **p<0.01, *ns* p>0.05 with Student’s t-test, two-tailed, equal variance.

Because this observation was unexpected, given that SCF^P0^ enables vacuolar degradation of ARGONAUTE proteins (Derrien *et al*, 2012; Michaeli *et al*, 2019), we explored their localization relative to TuYVs81 VRCs using the B2-GFP assay in *N. benthamiana*. In contrast to what was observed for DCL proteins, tRFP-AGO1 did not fully colocalize with the dsRNA but rather re-localized to the immediate vicinity of TuYVs81 VRCs, with which it appeared to be enmeshed (**Figure 5D** **and S8A**). This localization was reminiscent of that of the ssRNA and CP of potato virus X (PVX) relative to that of B2-GFP labelled dsRNA, that are partitioned within larger “viral factories” (Monsion *et al*, 2018) also referred to as X-bodies (Tilsner *et al*, 2012). This suggests that AGO1 does not fully access TuYV VRCs, which contain dsRNA, but rather peripheral viral structures that could be rich in viral ssRNA. On the contrary WT P0-tRFP, which presented bright punctate cytosolic structures in non-inoculated leaves, colocalized with the VRCs in infected leaves (**Figure 5E** **and S8B**). Thus, AGO1 and P0 localizations only partially overlap during TuYVs81 infection, suggesting that only a fraction of AGO1 would be available for ubiquitination by viral P0 at any given time.

Since *AGO1* mRNA level is under tight control by the AGO1-miR168 feedback loop (Mallory & Vaucheret, 2009), we monitored miR168 abundance in TuYVs81-infected Arabidopsis vasculature (**Figure 4B** **and S7C**) and found it to be unchanged by TuYV. Accordingly, *AGO1* and *AGO2* mRNA abundance was identical in mock and TuYVs81-infected vasculature (**Figure 5F, S7F and S7G**), ruling out a transcriptional or post-transcriptional regulation of AGO1 caused by the virus. We also observed stabilization of a vascular-restricted Flag-AGO1 protein (pSUC:Flag-AGO1, two independent lines) in TuYVs81 infected whole leaves (**Figure S7D**) that itself did not result from transcriptional activation of the transgene (**Figure S7E**). We also tested the possible involvement of the viral P0 in AGO1 stabilization, or a correlation between this phenomenon and TuYVs81 processing by DCL2 by analyzing AGO1 abundance in vasculature isolated from TuYVs81P0^-^ infected plants, as well as in *dcl2-1* plants. In both cases, AGO1 overaccumulation was still evident, ruling out involvement of P0 and DCL2 (**Figure 5G**).

Finally, we tested the ability of P0 to degrade AGO1 and its consequences on silencing in vascular cells by engineering a P0-HA stable line under the control of the Commelina yellow mottle virus promoter (pCoYMV) that drives specific expression in the companion cells of leaves, stems and roots (Matsuda *et al*, 2002). This transgene was introduced into the SUC-SUL background (Himber *et al*, 2003) to monitor silencing. Lines expressing the transgene were confirmed by RT-qPCR in *in vitro* grown seedlings and exhibited vein yellowing (**Figure S9**), in contrast to a pSUC:P15-FHA lines that prevented movement of *SUL* siRNA to the neighbouring cells (Incarbone *et al*, 2017). Strikingly, older plants containing the WT P0-HA exhibited a strong enhancement of leaf yellowing that was absent from plants of the same age containing the LP1 version of P0-HA (**Figure 5H**), suggesting that P0 facilitates cell-to-cell movement of *SUL* siRNA. Isolation of vasculature revealed the enrichment of the P0-HA protein compared to whole leaves and the clear degradation of vascular AGO1 in the presence of P0-HA, that was dependent on an intact F-box motif (**Figure 5I**), demonstrating that vascular cells are a suitable environment for P0-mediated degradation. Degradation of AGO1 was accompanied by stabilization of miR160c*, a hallmark of miR160 duplex accumulation, as well as destabilization of guide miRNA (**Figure 5J**). *SUL* siRNA production was not changed by P0-HA presence in the vasculature, indicating that movement rather than biogenesis was impaired. Overall *CHLI1* and *CHLI2* mRNA were decreased in the pCoYMV:P0-HA-WT plants, due to *SUL* siRNA accessing transcripts far removed from the vascular initiation site (**Figure 5K**). Vascular *AGO1* homeostasis was perturbed (**Figure 5K**), most likely as a consequence of disturbance in the miR168-AGO1 feedback loop. All in all, we demonstrate that vascular P0, when produced by a transgene, is competent in degrading AGO1, and the resulting depletion of AGO1 potentiates the cell-to-cell movement of siRNA to the surrounding tissues. Surprisingly, TuYVs81 infection results in stabilization of vascular AGO1 via an unknown mechanism that is not caused by a post-transcriptional regulation and depends neither on P0 nor DCL2.

### AGO1-loaded 22-nt vsiRNA promote production of secondary siRNA from Arabidopsis transcripts

Given that 22-nt sRNA can initiate secondary siRNA cascade from AGO-targeted RNA (Chen *et al*, 2010; Cuperus *et al*, 2010), we wondered if the abundant TuYV-derived 22-nt vsiRNA could lead to production of secondary siRNA from Arabidopsis transcripts. We therefore performed pairwise differential enrichment analysis of sRNA reads mapping to the Arabidopsis genome between infected and mock samples and found very little change apart from a few loci. **Figure 6A** shows the twenty most deregulated loci across all IP datasets, with the two bottom-most clusters corresponding to *MIR* genes and phasiRNA producing loci that we have previously shown to be affected by the *ago1-57* mutation (Derrien *et al*, 2018). The top-most cluster contains nine loci that accumulate sRNA reads only in the presence of TuYVs81, and analysis of the total RNA datasets shows a similar behaviour for five out of nine loci and uncovers an additional gene following this pattern (**Figure S10A**). Additional quantitative analysis of the AGO1 IPs revealed very few loci presenting sRNA enrichment with an adjusted p.value < 0.05 that did not overlap with preexisting sRNA, namely AT1G18480, AT5G45930, AT5G11000 and AT4G31540 (**Figure S10B**). The fact that these sRNA cohorts are bound to AGO1 and less abundant in AGO1-57 IPs suggests that they are part of a functional RISC, and that their biogenesis is partly affected by the *ago1-57* passenger strand retention phenotype.

**Figure 6:**
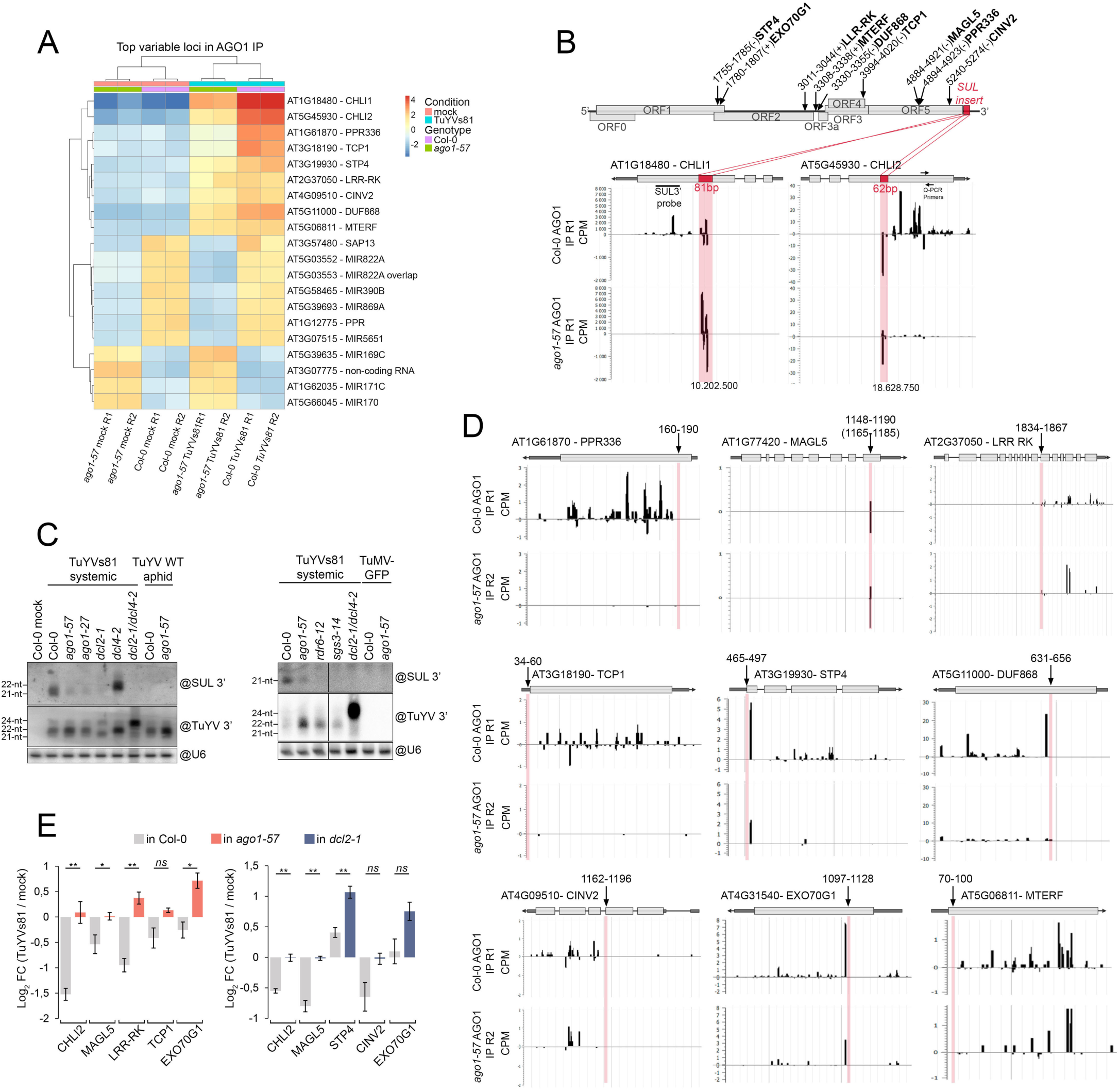
AGO1-loaded viral 22-nt siRNA promote production of secondary siRNA from Arabidopsis transcripts. **(A.)** Heatmap of annotation units showing the most variation in small RNA abundance across the eight AGO1 IP libraries. **(B.)** Top panel: Schematic representation of the TuYVs81 genome. Coordinates and strandedness of predicted discrete TuYV regions spawning sRNA for the indicated genes (psRNATarget 2017 release, expectation ≤ 3.5). *SUL* insert is represented as a red square in 3’ of the viral sequence. Bottom panel: Browser view of normalized sRNA reads (CPM count per million of mapped reads) mapping on both strands of *CHLI1* and *CHLI2* genes (0mm). Red squares represent the regions in the transcript that are identical to the *SUL* insert of TuYVs81. sRNA reads present within the pink highlighted area are directly produced from the viral RNA and trigger the production of the secondary siRNA population found in 3’ of both transcripts. Production of secondary siRNA is impaired in *ago1-57* plants. **(C.)** Production of 21-nt secondary siRNA produced from CHLI1 depends on the canonical AGO1/SGS3/RDR6/DCL2/4-mediated amplification pathway required for S-PTGS. Left panel: Analysis of sRNA abundance in total RNA extracted from the whole leaf of the indicated genotypes in either non-infected (E.V), TuYVs81 infected and WT TuYV infected aphid transmitted by RNA gel blot. For E.V and TuYVs81, a mixture of infected leaves from several individual was used (n=4) at 20 dpi. For aphid-infected samples, RNA from representative individuals in Figure 2H was used. Right panel: Analysis of sRNA abundance in total RNA extracted from the whole leaf of the indicated genotypes in either TuYVs81 or TuMV-GFP infected plants at 14 dpi. “@” indicates hybridization with indicated DNA probe and U6 signal is the loading control. Position of the SUL 3p probe is indicated in **B**. **(D.)** Browser view of normalized sRNA reads (CPM count per million of mapped reads) in AGO1 IP libraries from Col-0 and *ago1-57* plants for the nine vsiRNA-targeted Arabidopsis genes. The position of the predicted region directly targeted by primary vsiRNA is indicated above the scheme and highlighted in pink. Production of secondary endogenous siRNA from these genes is partially affected by the *ago1-57* mutation. **(E.)** mRNA abundance of the indicated target genes in isolated vascular tissues measured by RT-qPCR and expressed as Log_2_ fold change in TuYVs81-infected over mock.. Data from both panels are from separate experiments. Left panel: enriched vascular bundles of Col-0 and *ago1-57* plants (n=24 leaves) at 16 dpi. Right panel: enriched vascular bundles of Col-0 and *dcl2-1* plants (n=20 leaves) at 17 dpi. Represented values are means of technical triplicates, error bars represent the SEM. **p<0.01, *p<0.05, *ns* p>0.05 with Student’s t-test, two-tailed, equal variance.

The two top variable loci in all cases are *CHLI1* (AT1G18480) and *CHLI2* (AT5G45930). Since TuYVs81 contains the 81bp reporter fragment of *CHLI1* and that 62bp of that insert present an almost 100% sequence identity with *CHLI2*, we checked the siRNA distribution along these two loci. As expected, the regions overlapping with the s81 insert contained abundant siRNA, but an additional population of siRNA was found in 3’ of the two transcripts, that cannot be directly attributed to TuYVs81 processing, and was mostly lost in AGO1-57 IP (**Figure 6B**). We interpret this observation as a primary source of abundant 22-nt vsiRNA originating from the viral genome loaded into AGO1 and initiating production of secondary siRNA downstream of the targeting site. Since these primary vsiRNA are made from a largely double stranded RNA precursor, they form perfectly matched duplexes that are retained into AGO1-57 (Derrien *et al*, 2018), thus disabling production of transitive siRNA in the mutant. To ascertain this hypothesis, we analyzed *CHLI1*-derived siRNA (SUL 3’) by northern blot (**Figure 6C**). This revealed that TuYVs81-triggered secondary siRNA are 21-nt in length, are greatly diminished in both *ago1-57* and *ago1-27*, are entirely lost in *dcl2-1*, *rdr6-12* and *sgs3-14* mutants, and that DCL2 can replace DCL4 for their biogenesis. Their production is specific to TuYV containing the s81 insert, as neither WT TuYV nor the unrelated TuMV-GFP elicited their production. Thus, DCL2 processing of TuYVs81 is necessary and sufficient to induce transitivity from Arabidopsis transcripts through loading into AGO1, production of dsRNA by RDR6/SGS3 and subsequent processing by DCL4/DCL2.

We next wondered what could be the cause of siRNA production from the other unrelated loci. We hypothesized the existence of discrete TuYV regions that present complementary base-pairing with the candidate genes and that support production of 22-nt vsiRNA that are sufficient to guide *de novo* transitivity. We employed psRNATarget V2 (Dai *et al*, 2018) to predict which cohort of vsiRNA could match the mRNA sequence of the nine candidates. We defined a target region with the following criteria: existence in 5’ of the observed secondary siRNA and expectation score ≤ 3.5. This produced nine predicted TuYV-target paired regions of an average 31bp (±5bp) for the plant transcripts and 30bp (±4bp) for TuYVs81, that contained overlapping vsiRNA, the majority of which were 22-nt in length. Their positions on the TuYVs81 genome are indicated in **Figure 6B**, with the corresponding matched sequence indicated above all target genes in **Figure 6D**. For eight of these targets, some secondary siRNA were observed in infected leaves with perturbed patterns in *ago1-57* suggesting that their biogenesis is affected when unwinding is impaired. In only one instance, MAGL5, a single peak was observed that overlapped with the predicted targeting region. This peak corresponds to a single 21-nt vsiRNA originating from the minus strand of the virus that has 100% identity to 20bp of the sense *MAGL5* messenger. Accordingly, no signal amplification was observed from *MAGL5*. Functional annotation clustering (Huang *et al*, 2009) only revealed a moderate enrichment (cluster1 – score 0.86) for GO terms related to membrane and cell periphery for six out of nine genes (**Figure S10C**). Because targeting by vsiRNA implies downregulation in TuYVs81-infected vasculature, we measured the mRNA abundance of each target in infected and mock vasculatures by RT-qPCR, using *CHLI2* as benchmark (**Figure 6E**). For six out of nine genes, we observed a decrease in infected plants in one or two independent infection experiments. This decrease was lost, or an increase was observed, in vasculatures of both *ago1-57* and *dcl2-1* infected plants, supporting this phenomenon as a direct consequence of the siRNA production and amplification loop. All in all, we show that host transcript stability can be controlled by viral encoded, host produced vsiRNA, and these transcripts can directly lead to the production of vascular secondary siRNA. This “silencing cascade” strategy could be commonly employed by phloem-restricted poleroviruses and potentially others, to fine tune host gene expression in this key tissue.

## Discussion

### Antiviral RNA silencing mechanism against phloem-restricted TuYV

In this study we dissected the molecular machinery mediating antiviral silencing of the phloem-restricted polerovirus, TuYV, which revealed several unusual characteristics. First, we find that DCL2, rather than DCL4, is the major supplier of antiviral sRNA, which results in accumulation of large quantities of 22-nt vsiRNA produced in the vasculature, in accordance with the tropism of poleroviruses. This is in stark contrast with most viruses that have been described and form compatible interactions with Arabidopsis, which usually relies on DCL4 to produce the bulk of antiviral siRNA (Bouché *et al*, 2006; Deleris *et al*, 2006; Diaz-Pendon *et al*, 2007; Donaire *et al*, 2009; Garcia-Ruiz *et al*, 2010; Wang *et al*, 2011), or for which processing by DCL2 is a consequence of virus manipulation, like in the case of TCV (Qu *et al*, 2008; Azevedo *et al*, 2010). For TuYV, we do not observe any perturbation of the activity of DCL2 and DCL4 in infected vasculature, nor does loss of viral P0 affect the pattern of vsiRNA production. This suggests that if DCL2 processing is caused by viral manipulation, then it is not exerted via its VSR but rather via a factor that remains to be identified. However, it is just as likely that processing by DCL2, rather than a consequence of viral modification of plant processes, could be a specific plant reaction to TuYV and/or poleroviruses in general, highlighting the promise of this virus as a tool to probe the little-understood antiviral biology of DCL2.

Nevertheless, both DCL2 and DCL4 are required to mount a full antiviral response, as loss of both results in a dramatic accumulation of TuYVs81 RNA, whereas the corresponding single mutants only allow two-fold accumulation of TuYV. Thus, the outcome of a WT TuYV infection resembles the outcome of infection by many VSR-deficient RNA viruses, and for which only loss of both DCL2 and DCL4 restores systemic movement. Accordingly, using an assay to visualize viral replication complexes, we show that both DCL2 and DCL4 are able to re-localize to TuYV VRCs in *N. benthamiana* infected leaves, showing that both retain their ability to access viral dsRNA, likely via their dsRNA-binding motifs and/or their cofactors. Although the reason for which DCL2 over DCL4 cleaves TuYV remains to be investigated, a recent work provides insight into DCL4 processing ability *in vivo* and might be applicable here: recent evidence suggests that point mutants of DCL4 are sufficient to uncouple production of 21-nt siRNA from RDR-dependent dsRNA substrates and intramolecular foldback structures, although binding to dsRNA is retained (Montavon *et al*, 2018). This suggests that DCL4 enzymatic activity is sensitive to the intrinsic properties of target RNA, and that its processivity might be more tunable than that of other DCL proteins. Thus, although both can access the dsRNA, perhaps TuYV dsRNA structural properties are incompatible with efficient processing by DCL4, leaving DCL2 free to cleave it into 22-nt. One might also ponder if dependence on DCL2 might be common in nature and in diverse pathosystems, in other words, if Arabidopsis Col-0 apparent low reliance on DCL2 represents the exception rather than the norm. Interestingly, abundant 22-nt vsiRNA were detected in cotton plants infected with cotton leafroll dwarf virus (CLRDV) (Silva *et al*, 2011), in maize leaves containing a novel polerovirus (Chen *et al*, 2016) and in wheat infected with barley yellow dwarf virus (BYDV) (Shen *et al*, 2020), suggesting that DCL2 is active against *Luteoviridae*.

Another unexpected feature observed in our study is the simultaneous reliance on AGO1 for silencing, with its specific stabilization in infected vasculature. Effective antiviral silencing requires the function of AGO–vsiRNA effector complexes and for most well-described RNA viruses, AGO2 seems to play a major antiviral role, alongside AGO1 (Harvey *et al*, 2011; Jaubert *et al*, 2011; Brosseau & Moffett, 2015; Ma *et al*, 2015; Wang *et al*, 2011; Garcia-Ruiz *et al*, 2015). The situation is quite different in the case of TuYVs81, for which the *ago1-57* mutation was sufficient to induce a two-fold increase in the amount of viral RNA and the addition of the *ago2-1* mutation did not lead to further increase. We also found that the vast majority of vsiRNA were loaded into AGO1, further underscoring its importance. Remarkably, none of the defects observed in *ago1-57* were observed in *ago1-27*, which is the most commonly used hypomorphic *ago1* allele, and the one used in most studies to assess the importance of AGO1 relative to other AGOs. It is therefore possible that the contribution of AGO1 to antiviral silencing in many plant-virus interactions has been underestimated. Use of the *ago1-57* allele, that uncouples AGO1 activity in siRNA pathways from miRNA pathways and shows little developmental phenotype will no doubt prove valuable to future investigations.

### Curtailed role of P0 in the context of TuYV infection

Along with the sole reliance on AGO1, we clearly observe increased accumulation of the protein, only in infected vasculature, a feature that did not extend to AGO2 or AGO4. This is counterintuitive, as the experimentally validated action of the VSR P0 is the degradation of unloaded AGO molecules to prevent formation of vsiRNA-RISC. We further show that this increase in vascular AGO1 protein is not a response to the viral P0, nor a consequence of the strong accumulation of 22-nt vsiRNA, as *dcl2-1* plant still exhibit the response. The differences between the results obtained with P0 in heterologous systems and the results reported here obtained during genuine virus infection are stark. This contrast highlights how expression of viral proteins as overexpressed transgenes, while often necessary to study their molecular features *in vivo*, does not always adequately recapitulate their full activity within the viral lifecycle.

When reconstituting the P0 degradation system in the vasculature, by expressing P0 under a companion cell promoter, we show that P0 leads to AGO1 degradation in the enriched vasculature and that this degradation depends on its F-box motif. Thus, the local environment of the vasculature does not hinder the function of P0, nor is vascular AGO1 insensitive to P0. Interestingly, we observe an increase of leaf chlorosis in SUC-SUL plants that contain WT vascular P0, a phenotype opposite to that of a phloem-restricted VSR that sequesters sRNA duplexes and leads to suppression of SUL siRNA cell-to-cell movement (Incarbone *et al*, 2017). We interpret this observation as a direct consequence of the decrease of AGO1 (and possibly additional AGOs) in the incipient cells, which allows an increased population of mobile siRNA duplexes to exit the companion cells. This leads to increased reach of the silencing signal to additional recipient cells, where these siRNA are loaded in turn and cause downregulation of *CHLI1/2*. Our observation is consistent with the “consumption” model proposed by Devers *et al*, (2020), in which cell-autonomous AGO proteins consume mobile sRNA as they travel cell-to-cell, leading to a modulation of the traveling silencing signal. Our results show that this holds true at the initiation site - provided it contains an active AGO1 population - as exemplified here in the vasculature.

As commented above, however, these results are in contrast with our data from TuYV infections. Given that AGO1 over-accumulates in TuYV-infected vasculature, it is unlikely that the production of the P0 protein from the viral genome is sufficient to cause measurable AGO1 decay. This gives rise to a dilemma regarding the role of P0 during genuine infection. How is P0 relevant as a VSR for poleroviruses? What causes the increase of AGO1 protein levels at the site of infection? A possible answer to the latter question is that the very abundant vsiRNA loaded into AGO1 could increase its stability and render it unsusceptible to P0-mediated degradation. In this scenario, *de novo* synthetized AGO1 molecules undergo rapid loading, are unavailable to both P0 and endogenous degradation pathways, and reiterated use of vsiRNA-RISC artificially prolongs their half-life, raising the overall population over time in the infected tissues.

As underlined in the introduction, strong initiation at the start of ORF0 is not favoured by TuYV and PLRV (Pfeffer *et al*, 2002; Juszczuk *et al*, 2000) and several isolates of polerovirus encode P0 proteins that perform poorly in silencing suppression assays, suggesting that strong suppression activity is not necessarily favoured by poleroviruses at large. The observations gathered here support the notion that the activity of TuYV P0 is mostly curtailed during infection. First: vascular AGO1 is stabilized during TuYV infection (although this does not rule out that a fraction of the AGO1 molecules can undergo P0-directed degradation). Second: non-degradable AGO1-57 offers no advantage to the host in the context of the natural mode of infection. Third: P0 and AGO1 have contrasting localizations relative to the VRCs, with AGO1 localizing in the proximity of the VRCs, while P0 signal overlaps with VRCs. The specific localization of P0 at what is most likely the replicating viral genome contrasts with its described localization when overexpressed in absence of the TuYV RNA and suggests that P0’s main activity during infection is restricted to the VRCs rather than spread throughout the cell. We therefore propose that viral P0 is mostly acting at the site of TuYV replication and causes degradation of a relatively small pool of AGO1 at contact sites between viral factories and *bona fide* replication complexes.

Tolerating a certain level of antiviral silencing and therefore minimizing perturbation to the host miRNA pathways is expected to increase host fitness during infection and could be a driving mechanism for adaptation, likely to be favoured over time. Tolerance could be achieved by temporally and/or spatially restricting silencing suppression, as we propose for TuYV P0. Furthermore, this would theoretically allow re-routing of the host RNAi pathways for the benefit of the pathogen, as discussed below. In accordance with this notion, several recent examples of viruses dampening their VSRs have emerged. The coat protein of Cucumber mosaic virus (CMV Fny strain) inhibits the translation of the 2b suppressor, which allows a degree of antiviral silencing, promoting self-attenuation and symptom recovery in the infected plants (Zhang *et al*, 2017). Similarly, βC1 protein of Cotton leaf curl Multan Betasatellite, a virulence factor for begomoviruses that exhibits silencing suppression activity (Amin *et al*, 2011), seems to be both an activator and a target of autophagy, effectively driving its own downregulation, leading to milder symptoms (Haxim *et al*, 2017; Ismayil *et al*, 2020). We expect that more examples of viruses exercising ‘self-control’ via different strategies will emerge in the coming years.

### DCL2/AGO1-dependent hijacking of vascular antiviral silencing

Finally, we observe that some of the TuYVs81-derived vsiRNA can trigger the production of siRNA from host encoded transcripts that bear the hallmarks of secondary siRNA. These are due to microhomologies between the plant transcripts and the viral RNA genome. Although there are some examples of vsiRNA targeting host transcripts in several plant-virus interactions (reviewed in Ramesh *et al*, 2021), they have mostly been associated with pathogenesis and symptoms (Smith *et al*, 2011; Shimura *et al*, 2011; Navarro *et al*, 2012; Yang *et al*, 2019). This is fundamentally at odds with the notion that many plant viruses produce strong VSRs which, by definition, restrict the use of vsiRNA, including targeting of host transcripts. As discussed above, TuYV restricts the use of its own VSR, thus offering a solution to this conundrum. Furthermore, TuYV infection of Arabidopsis is symptomless, suggesting that unlike the examples above, the function of these vsiRNA is unrelated to symptom manifestation.

It is significant that the detection of these vsiRNA-primed secondary siRNA is made possible by several key findings of this study. First: TuYVs81 RNA is cleaved by DCL2 over DCL4 to generate a cohort of 22-nt vsiRNA, amongst which the initiators of host mRNA targeting. This is of importance as 22-nt sRNA can initiate secondary siRNA cascades while 21-nt isoforms cannot (Chen *et al*, 2010; Cuperus *et al*, 2010). Second: TuYVs81 siRNA are preferentially loaded into AGO1 rather than AGO2. This fact also concurs to the production of secondary siRNA, as AGO1 can initiate tasiRNA biogenesis (Vazquez *et al*, 2004; Arribas-Hernández *et al*, 2016). By contrast, AGO2 loaded with a 22-nt siRNA can direct cleavage of the matching RNA but cannot stimulate production of secondary siRNA (Carbonell *et al*, 2012). Third: We observe curtailed action of P0 and stabilization of the AGO1 protein. This is necessary, as having excessive P0 activity would disable the targeting of these transcripts. Taken together, these observations suggest that TuYV takes advantage of the DCL2-AGO1-RDR6 axis to modify the vascular transcriptome of its host. This is made possible by the restrained VSR activity of P0. To our knowledge, this is the first report of primary vsiRNA causing amplification and spreading of silencing signal on plant host transcripts.

However, what benefit does the virus obtain from such manipulation is currently unknown. It is possible that the observed microhomologies and subsequent targeting are merely accidental and tolerated by the host. It is also unknown if the use of the amplification system is favored over a non-amplifying system in order to target additional host transcripts, or if the appearance of secondary siRNA simply reflects the reliance on DCL2 and AGO1. Among the targets, two genes, STP4 (SUGAR TRANSPORTER 4) and CINV2 (CYTOSOLIC INVERTASE 2), are involved in sugar transport and carbon partitioning. Cytosolic invertase can break down sucrose into monosaccharides, while STPs are monosaccharide/proton symporters located at the plasma membrane and responsible for the uptake of sugars. Their regulation during infection could fine tune levels of cytosolic monosaccharides in infected cells. Interestingly, STP4 expression is induced in response to aphid feeding (Moran & Thompson, 2001), a prerequisite to TuYV infection. PPR336/rPPR1 and MTERF are both organelle proteins, the first being an integral part of the mitoribosome (Waltz *et al*, 2019) and the second predicted to be involved in regulation of organelle transcription (Kleine, 2012). MAGL5 is part of a family of chloroplastic monoacyglycerol lipase, whose activity releases glycerol and fatty acids (Kim *et al*, 2016). There is thus an enrichment in genes involved in carbon and energy metabolism, which could be partly reshaped by the infection. The set also contains two additional plasma membrane-localized proteins: LRR-RK, one of many plant leucine rich repeat receptor kinase that act as extracellular sensors for various ligands to help regulate development and immune responses (Sun *et al*, 2017) and a DUF868-containing protein, which is a conserved domain found exclusively in plant proteins. On the other hand, EXO70G1 is part of the conserved exocyst complex, which is involved in the secretion of post Golgi-vesicles, enabling delivery of lipids and proteins to the plasma membrane (Chong *et al*, 2010). Finally, we also find TCP1/CCT4, a component of the chaperonin oligomeric complex involved in tubulin folding (Ahn *et al*, 2019). Deciphering how these genes contribute to the infection and/or the host response will help to better understand plant-polerovirus interactions.

Finally, we wondered if the observations obtained from a TuYV-*A. thaliana* interaction could be extended to other polerovirus-plant interactions, especially for crop species like cabbage and beet in which poleroviruses cause extensive symptoms. Using thirteen species representative of angiosperm diversity, we were able to find orthologues for all target genes identified in *A. thaliana*. This approach revealed the extent of the conservation of the target regions, with well-conserved blocks in *Brassicaceae* and in the tested dicotyledon species for most, while in the case of *LRR-RK* and *CINV2*, target region conservation was also observed in the two monocotyledon species tested (**Figure S11**). The only exception was *TCP1*, which is targeted in the 5’UTR and presented two major insertions relative to *A. thaliana*. Interestingly, targeting regions cluster in six positions along the TuYV genome (**Figure 6B**) with overlapping regions for three gene pairs. A conservation analysis performed on representative either beet-infecting (BMYV, BChV, BWYV-USA) or non-infecting (TuYV, CABYV, PLRV) poleroviruses revealed good conservation in protein coding regions of the genome, with the exception of the *MTERF-DUF868* pair, that was located at the start of the subgenomic RNA and for which sequence identity with TuYV was only found in brassica yellows virus (BrYV-ABJ) (**Figure S12**). This suggests that several polerovirus species could potentially target the same cluster of genes in their respective plant hosts, opening new avenues in the dissection of viral manipulation of host transcriptomes.

## Materials and Methods

### Plant lines

AGO1 point mutants have been described previously: *ago1-57* (Derrien *et al*, 2018), *ago1-38* (Gregory *et al*, 2008), *ago1-27* (Morel *et al*, 2002). *ago2-1* (SALK_003380), *ago5-1* (SALK_063806), *ago7-1* (SALK_037458), *ago10-3* (SALK_519738) and the resulting combination mutants have been described in Wang *et al*, 2011. *dcl2-1* (SALK_064627), *dcl4-2* (GABI_160G05), *dcl2-1/dcl4-2* (Deleris *et al*, 2006), *dcl2-5/dcl3-1* (Blevins *et al*, 2006) *dicer-like* single and combination mutants have been described previously. *rdr6-12* and *sgs3-14* (SALK_001394) (Peragine, 2004), *atg5-1* (SAIL129_B07) (Thompson *et al*, 2005), *atg7-2* (GABI655_B06) (Chung *et al*, 2010), SUC-SUL (Himber *et al*, 2003), SUC-SUL/ pSUC:P15-FHA (Incarbone *et al*, 2017) have been described previously. *ago1-57/ago2*-*1*, *ago1-57/dcl2-1* and *ago1-57/dcl4-2* plants were obtained by crossing their respective single mutants and selected by genotyping of the F2 population. SUC-SUL/*ago1-57* were obtained by crossing and selected by genotyping and for Basta resistance. The triples *dcl2-1/dcl4-2/ago1-57* was obtained by crossing the double *dcl2-1/dcl4-2* with the double *dcl2-1/ago1-57* and selected by genotyping.

### Constructs

35S:CFP-AGO1 construct and its mutant derivative were previously described in Derrien *et al*, 2018.

For overexpression constructs of P0: TuYV (Baumberger *et al*, 2007), CABYV (Pazhouhandeh *et al*, 2006), BMYV (Klein *et al*, 2014) and PLRV (Pfeffer *et al*, 2002), all ORF0s were cloned into a pBIN vector, under a 35S promoter as described in Pfeffer *et al,* (2002).

35S:tagRFP and 35S:tagRFP-AGO1 are in the pEAQΔP19-GG vector and were cloned as described in Incarbone *et al*, (2020). *AGO1* cDNA was amplified with primers containing SapI restriction sites and combined with N-terminal tagRFP fragment in a Golden Gate reaction.

For 35S:P0-tagRFP constructs, the P0 coding sequences with T156A/T159C mutations (to avoid translation of a truncated P1 protein from the P0 vector) were amplified from the XVE:P0-myc constructs (Bortolamiol *et al*, 2007; Derrien *et al*, 2018) with oligo primers containing the AttB1 and AttB2 sites and the PCR product was cloned in a pDONR221 by BP recombination. The entry clones were then recombined with the pGWB660 vector by LR gateway reaction producing 35S promoter driven P0 construct fused to tagRFP at the C terminus.

For 35S:DCL2-tagRFP and 35S:DCL4-tagRFP constructs the genomic sequence from start codon to the last codon before the stop codon was amplified with oligo primers containing the attB1 and AttB2 sites and the PCR product was subsequently cloned in a pDONR221 by BP recombination. The entry clones were then recombined with the pGWB660 vector by LR gateway reaction producing 35S promoter driven genomic constructs fused to tagRFP at their C termini.

To obtain the pCoYMV:P0-HA lines, the promoter sequence of CoYMV (1039bp) was amplified from the vector pGEM 3zf+ CoYMV-Suc51 (Srivastava *et al*, 2009) with oligo primers containing AttB4 and AttB1R recombination sites and the sequences was mobilized into the pDONR P4P1r vector by BP Gateway recombination. The P0-3xHA fusions (WT or LP1) were obtained by seamless Gibson assembly (NEB) of the P0 coding sequences with T156A/T159C mutations and a 3xHA-stop fragment with overlapping primers into the pFK202 plasmid. The resulting fusion was then amplified with AttB1 and AttB2 containing primers and mobilized into the pDONR221 plasmid by gateway BP reaction. Pieces were assembled into the pK7m24GW plasmid (http://www.psb.ugent.be/gateway/) by double recombination LR reaction to obtain the final binary plasmid. The plasmid was then introduced into Agrobacterium strain GV3101 pMP90 and used to transform SUC-SUL plants (Clough & Bent, 1998).

To obtain the pSUC:Flag-AGO1 lines, the SUC2 promoter (943bp) was amplified from the pEP1 plasmid (Imlau *et al*, 1999) with oligo primers containing AttB4 and AttB1R recombination sites and the sequences was mobilized into the pDONR P4P1r vector by BP Gateway recombination. The 3xFlag sequences was amplified from a pENTRY-3xFlag vector with primers containing attB1 and attB2 sites and a 5’ Kozak consensus sequence and the PCR product was subsequently cloned in a pDONR Zeo by BP recombination. AGO1 CDS sequence was amplified with primers containing attB2R and attB3 sites and mobilized into the p2R-P3 plasmid by gateway BP reaction. All fragments were assembled by three-way LR gateway reaction into the pB7m34GW vector. The plasmid was then introduced into Agrobacterium strain GV3101 pMP90 and used to transform Col-0 plants.

For cloning PCR products, Phusion high fidelity DNA polymerase 2X master mix (Thermo Scientific) was used, except for the genomic DCL2 and DCL4 for which Platinum SuperFi DNA Polymerase (Invitrogen) was used, following the manufacturer’s protocol. All clones were verified by in-house Sanger sequencing before proceeding to the next step. All primers used for cloning are available in **supplemental Table 4**.

### Plant growth and infection conditions

For *in vitro* culture, seeds were surface sterilized using ethanol, plated on growth medium (MS salts [Duchefa], 1% sucrose, and 0.8% agar, pH 5.7), stored 2 days at 4°C in the dark, and then transferred to a plant growth chamber under a 16-h-light/8-h-dark photoperiod (22°C/20°C). For standard plant growth, seeds were directly sown on soil (Hawita Fruhstorfer) in trays and kept under a 12-h-light/12-h-dark regime for 14 days, then transferred in 16-h-light/8-h-dark growth chambers, under fluorescent light (Osram Biolux 49W/965).

For infection conditions, seeds were directly sown on soil (Hawita Fruhstorfer) in trays and kept under a 12-h-light/12-h-dark regime for about 4 weeks. *Agrobacterium tumefaciens* containing binary plasmids of TuMV-GFP, TuMV-AS9-GFP, TRV-PDS (RNA1+RNA2-PDS) recombinant viruses were grown overnight at 28°C in antibiotic-supplemented LB media, pelleted 15 minutes at 4000g and resuspended in infiltration medium (10mM MgCl_2_/10mM MES/200µM Acetosyringone). The cells were then adjusted to an OD_600nm_ of 0.5 and agroinfiltrated into three leaves per plants with a needleless syringe. Plants post-inoculation were then grown in either 12-h-light/12-h-dark period with fluorescent light or under long day condition in greenhouse. For TuYVs81, TuYVs81P0^−^ or the empty vector (mock), the procedure was the same with the following modifications: after pelleting of the initial cultures, cells were resuspended in 10.5 g/L K2HPO_4_/4.5 g/L KH_2_PO_4_/1 g/L (NH_4_)_2_SO_4_/0.5 g/L sodium citrate/0.1 g/L MgSO_4_/0.4% (v/v) glycerol/0.1 g/L MES/200 μM Acetosyringone and incubated 5-6 hours in the dark. Cells were pelleted again and resuspended in infiltration medium and handled as described above. Infections were left to progress until the time indicated in the figure legends, and systemic leaves were collected from the indicated number of individuals. Plant picture were taken at the indicated time using a DSLR camera mounted on a stand at a fixed distance, or for systemic TuMV-AS9-GFP propagation, using a Zeiss axio zoom equipped with GFP filter.

For analysis of infiltrated leaves, the same protocol was used, but five leaves per plant were infiltrated with a needleless syringe. Only the inoculated leaves were sampled at the indicated time points. To account for biological variation, several inoculated leaves were harvested per time point. For one given experiment either all the inoculated leaves were taken off a set number of plants, or one inoculated leaf from each individual was harvested per time point, with identical result.

For aphid-mediated inoculation, seeds were directly sown on soil (Hawita Fruhstorfer) in trays and kept under a 12-h-light/12-h-dark regime for 18 days. Each plant was then challenged with two *Myzus persicae* fed on either 20% sucrose solution, 20% sucrose solution containing 67mg/ml TuYV virions or alternatively were left untreated. After 4 days, insecticide was applied to kill the aphids and plants were kept under a 12-h-light/12-h-dark period with fluorescent light for 25 days before sampling. Each plant was individually weighed (after removal of root system and cotyledons) and ground with liquid N_2_ with a mortar and pestle. For each, 100mg (± 5mg) were kept at -80°C in Eppendorf tubes for DAS-ELISA.

### Transient Expression Assays and local leaf infection in *N. benthamiana*

Binary constructs were transformed in Agrobacterium GV3101 pMP90 and infiltrated in *N. benthamiana* for transient expression assays. Agrobacterium cells were grown overnight at 28°C in 5 to 10 mL LB medium supplemented with antibiotics, resuspended in infiltration medium at an OD_600nm_ of 0.1-0.3 per construct, and incubated for 2 to 4 h at room temperature before being infiltrated into leaves of 4-week-old plants with a needleless syringe. Plants were maintained in growth chambers under a 16-h-light/8-h-dark photoperiod with a constant temperature of 22°C. For VRC colocalization assays, 35S:B2-GFP *N. benthamiana* plants (Monsion *et al*, 2018) were first screened under U.V light in order to select plants with visible GFP. These plants were infiltrated with a mixture of the TuYVs81 infectious clone or the empty vector and with the construct of interest. Plants were kept 3 days in the growth chamber before imaging.

### DAS-ELISA

TuYV was detected in non-inoculated leaves of *A. thaliana* by double-antibody sandwich enzyme-linked immunosorbent assay (Clark & Adams, 1977). Flat-bottomed NUNC 96-wells plates were coated with 1:400 BWYV IgG (Loewe) (Herrbach *et al*, 1991) in 1.59 g/L Na_2_CO_3_/2.93 g/L NaHCO_3_ pH=9.6, 100 µl per well and incubated at 37°C in the dark for 4 hours. N_2_ chilled samples were ground in safe lock Eppendorf tubes using a Silamat S7 (Ivoclar vivadent) and extracted in 400 µl 1X PBS/0.05% (v/v) Tween 20/2% (w/v) soluble PVP 360, resuspended by shaking for 15 minutes and centrifuged for 5 minutes at 3000rpm at 4°C. Coated plates were washed three times with 1X PBS/0.05% (v/v) Tween 20 and 100 µl of supernatant was loaded into each well for an overnight binding at 4°C. The plates were then washed three times in 1X PBS/0.05% (v/v) Tween 20, and 100 µl per well of 1:400 BWYV AP conjugate (Loewe) in 1X PBS/0.05% (v/v) Tween 20/2% (w/v) PVP 360 was added and plates were incubated at 37°C in the dark for 3 hours. Plates were washed three time as described above and 100 µl per well of freshly prepared substate buffer was added (1M diethanolamine/5mM MgCl_2_/1mg/ml substrate tablet [4-Nitrophenyl phosphate Na_2_-salt]). O.D_405nm_ was determined after 90 minutes of incubation. Technical triplicates were obtained for one given sample, and each plate contained a positive control (TuYVs81 with yellow veins) and a negative control (Mock-infected individual). Blank was obtained by performing the reaction on buffer only.

### Infection kinetic scoring by dot blot

A leaf disk (5mm Ø) from young systemic leaves of each individual plant inoculated with TuYVs81 was removed at the indicated time after inoculation. Protein extracts were obtained by adding 50 µl of 2X Laemmli buffer (200 mM Tris HCl pH 6.8/ 8% [w/v] SDS/ 40% [v/v] Glycerol/ 0.05% [w/v] Bromophenol Blue/ 3% [v/v] B-mercaptoethanol) to each leaf disk in 96-well plates and grinding at maximum speed for 3 minutes in a Retsch mixer mill at room temperature. The plates were then centrifuged for 5 minutes at 4000g and the supernatant was transferred to a fresh 96-well plate to denature proteins for 5 minutes at 95°C. 5 µl of the denatured protein sample was manually spotted onto an activated PVDF Immobilon-P membrane (Millipore) mounted on top of a paper stack (bottom up: dry stack of paper towel, one dry whatman filter paper cut to membrane dimension, one wet whatman filter paper cut to membrane dimension in 25mM Tris base/192mM glycine/20% (v/v) ethanol), for a maximum of 96 samples per membrane, mirroring the plate. The membrane was left to dry at 60°C for about 10 minutes, reactivated in absolute ethanol, washed in deionized water for 2 minutes, incubated in TBST +5% milk (20mM Tris base/150mM NaCl/0.1% Tween 20, pH 7.4) for about 20 minutes, washed in TBST without milk and hybridized overnight in 1:15000 (v/v) of anti-readthrough (RT) (Reutenauer *et al*, 1993) rabbit antibody in TBST +2.5% (w/v) BSA. Membranes were then processed as indicated for western blots, and signal was acquired on ECL films. Membranes were then colored in Coomassie blue to mark the position of the samples, dried, and film and membrane were overlayed to pinpoint infected individuals. This process was repeated several times for each inoculated plant, and individual plants were counted as positive only if at least two independent time points returned positive signals.

### Protein analysis and immunoblotting

N_2_ chilled samples were ground in safe lock Eppendorf tubes containing 2mm Ø glass beads, using a Silamat S7 (Ivoclar vivadent) and total proteins were extracted in 2X Laemmli buffer by mixing again in the Silamat S7 for 20 seconds. Samples were denatured for 5 minutes at 95°C and quantified using the amido black method. 10 µl of supernatant mixed with 190 µl of deionized water and added to 1 ml of normalized 10% (v/v) Acetic acid/90% (v/v) methanol/0,05% (w/v) Amido Black (Naphtol Blue Black, Sigma N3393) buffer, mixed and centrifuged for 10 minutes at maximum speed. Pellets are then washed in 1 ml of 10% (v/v) Acetic acid/90% (v/v) ethanol, centrifuged 5 minutes at maximum speed and resuspended in 0.2N NaOH. O.D_630nm_ was determined, with NaOH solution as blank, and protein concentration is calculated using the O.D=a[C]+b determined curve. 2.5 to 40 µg of total protein extracts were separated on SDS-PAGE gels and blotted onto PVDF Immobilon-P membrane (Millipore). AGO1 protein was detected using the anti-AGO1 antibody (rabbit polyclonal, AS09 527; Agrisera) diluted 1:10,000 (v/v). AGO2 protein was detected using the anti-AGO2 antibody (rabbit polyclonal, AS13 2682; Agrisera) diluted 1:5000 (v/v). AGO4 protein was detected using the anti-AGO4 antibody (rabbit polyclonal, AS09 617; Agrisera) diluted 1:5000 (v/v). Myc-tagged proteins were detected using anti-myc antibody (mouse monoclonal; Roche) di-luted 1:5000 (v/v), or anti-myc antibody HRP-coupled (mouse monoclonal, Miltenyi Biotec) diluted 1:2000 (v/v). CULLIN1 protein was detected using anti-CUL1 antibody (Shen et al., 2002) diluted 1:5000. GFP-tagged proteins were detected using the anti-GFP antibody (JL-8, Clontech Takara) diluted 1:2000 (v/v) or anti-GFP HRP-coupled (mouse monoclonal, Miltenyi Biotec) diluted 1:5000 (v/v). Flag-tagged proteins were detected using the anti-Flag HRP-coupled antibody (mouse monoclonal, A8592; Sigma-Aldrich) diluted 1:5000 (v/v). H-tagged proteins were detected using the anti-HA HRP-coupled antibody (mouse monoclonal, Miltenyi Biotec) diluted 1:5000 (v/v). tagRFP fusion proteins were detected using the anti-tRFP (rabbit polyclonal, Evrogen AB233) diluted 1:5000 (v/v). TuYV Readthrough protein was detected as described in the dot blot section. DCL1 protein was detected using anti-DCL1 antibody (rabbit polyclonal, AS19 4307; Agrisera) diluted 1:2000 (v/v). Mouse monoclonal antibodies were detected with a goat anti-mouse IgG HRP-linked antibody (62-6520; Invitrogen) diluted 1:10,000 (v/v). Rabbit polyclonal antibody were detected with a goat anti-rabbit IgG HRP-linked antibody (65-6120; Invitrogen) diluted 1:10,000 (v/v). Hybridized membranes were reacted with Clarity or Clarity Max ECL (Biorad) and imaged using a Fusion FX (Vilbert). For signal quantification, the plot lane function of ImageJ was used to obtain the raw intensity for a signal of interest as well as from the whole Coomassie stained lane. Each signal was subsequently normalized to the total protein signal, and the values obtained from all the replicates were plotted using with(data,boxplot(value∼treat)) command in R were data contains the imputed values, separated according to treatment (∼treat).

### RNA related methods

Total RNA extraction was performed in Tri-reagent (MRC) and RNA blots were performed as described in Derrien *et al*, 2018 with starting material between 10 to 20 µg total RNA. DNA oligo sequences used for [γ-^32^P]ATP labelling and DNA oligos used to generate the PCR products for [α-^32^P]CTP-labeled Klenow products are available in **supplemental Table 4**. For quantitative RT-PCR, 1-2 μg of total RNA treated with DNase I (Thermo Fisher Scientific) was reverse transcribed using either the High-capacity cDNA reverse transcription kit (Applied Biosystems) or the SuperScript IV Reverse Transcriptase (Invitrogen). All RT reactions performed from TuYVs81-infected samples and their control counterparts contained 2.5 μM random hexamers and 0.05 μM Tu-4942-rev primer. Quantitative PCR reactions were performed in a total volume of 10 μL of SYBR Green master mix I (Roche) in 384-wells plates on a Lightcycler LC480 apparatus (Roche) according to the manufacturer’s instructions. For each cDNA, technical triplicates were obtained from the same plate, and expression data was normalized using the SAND (AT2G28390) and EF1a (AT5G60390) genes as internal controls, using the ΔΔCt method.

### Immunoprecipitation of AGO1 and AGO2

Immunoprecipitation of endogenous AGO1 were performed as described in Derrien *et al*, 2018, using the anti-AGO1 and anti-AGO2 antibodies, described in the immunoblotting section. In both cases, 5 µg of water-resuspended antibody was used to bind to 30 µl of PureProteome Protein A magnetic beads (Millipore).

### Confocal microscopy

All confocal imaging was done using a Leica SP8 CLSM. For agro-infiltrated tobacco leaves, abaxial epidermal cells were imaged from leaf-disks (at least two disks per leaf, with at least 3 different leaves per combination, on separate plants). Leaf disks were mounted between a microscope slide and a coverslip, with a tape in between to account for the thickness and the coverslip was further taped to the slide before adding deionized water. Leaf disks were vacuum infiltrated before microscopy. Usual excitation/detection range parameters for GFP and tagRFP were 488 nm/505–520 nm and 561 nm/570–630 nm. For CFP and GFP simultaneous imaging, excitation/detection range parameters were 405 nm/440–475 nm, 488 nm/505–520 nm, respectively. Emissions were collected using the system’s hybrid (Hyd) detectors and sequential scanning was employed for all acquisitions. Images were processed using FIJI (Schindelin *et al*, 2012).

### Meselect for vascular bundle enrichment

To obtain one sample of vascular tissues for protein or RNA analysis, the indicated amount of leaves (in the figure legends) from a given genotype or treatment were harvested fresh from different plants. Leaves were carefully laid on their abaxial side to the sticky side of a labelling tape. Another tape was added on the adaxial side of the leaf, and this tape was slowly removed, taking away the adaxial epidermal cells. During preparation of the remaining leaves, the processed leaves on tape containing the abaxial epidermis were left face down in ice cold 0.4 M mannitol/10 mM CaCl_2_/20 mM KCl/0.1 % (w/v) BSA/20 mM MES pH 5.7. Once all were processed, they were incubated face down in the same buffer with the addition of 1% (w/v) cellulase Onozuka (Yakult) and 0.25% (w/v) macerozyme Onozuka R10 (Yakult) for 15 minutes under gentle rotation at room temperature. Mesophyll cells that were released in the protoplast buffer were collected by pipetting gently with a cut tip and centrifuged at 200g at 4°C with brakes on. The pelleted cells were extracted either in Laemmli 2X buffer for protein extraction, or in 1ml Tri-Reagent for RNA extraction. The tapes were left face down in ice-cold wash buffer (154 mM NaCl/125 mM CaCl_2_/5 mM KCl/5 mM Glucose/2 mM MES pH 5.7), were gently shaken, and the buffer was refreshed once for further washing. The vascular tissue network was then lifted off the leaves with tweezers, starting from the central vein, and collected into a tube with ice cold wash buffer until all were processed. All collected vascular bundles were quickly drained on paper towel, and flash frozen in a safelock Eppendorf tube containing 2mm Ø glass beads. Proteins were extracted in Laemmli 2X as described in the immunoblotting section, and RNA was extracted in tri-reagent after grinding of the frozen vasculatures in the silamat S7.

### sRNA library preparation

Total RNA samples and AGO1 IP samples were both obtained from Col-0 and *ago1-57* rosette leaves from TuYVs81 infected of mock -inoculated plants 16 days after inoculation. For both genotypes, only plants with vein yellowing were included in the analysis. Plants were separated into pools of 7-8 plants per biological replicate, and the resulting mixed and ground tissue was used as the input material for the AGO1 IP and the total RNA. About 100 mg of material was used to extract total RNA using tri-reagent as described, and IPs were performed from 1 g starting material. AGO1-loaded sRNAs were then extracted by adding Tri-Reagent directly on the magnetic beads. Library preparation and sRNA sequencing (single-read, 50bp, V4 chemistry) on Illumina HiSeq were performed by Fasteris (http://www.fasteris.com), with each sample split and sequenced in two independent lanes. FASTQ file generation, demultiplexing, and adapter removal were done by Fasteris.

### Mapping, quantification, and differential analysis of sRNA

Reads (18 to 26 nucleotides long) were aligned and quantified using using Shortstack v3.8.5 with the ‘fractional’ option (Johnson *et al*, 2016), allowing for no mismatches, with the Arabidopsis genome (TAIR10) and TuYVs81 genome as reference. Mapping statistics are provided in **Figure S2A**. Total 20 to 24 nucleotides reads mapped per library were normalized in read per million mapped (RPM) and the obtained values per category were plotted as stacked histograms using ggplot2 in R. Alternatively, RPM values for the 21, 22 and 24-nt subsets were used for filled histograms per category. To obtain the vsiRNA distribution graphs over TuYVs81 genome, bam files were used as input in MISIS (Mapped short interfering RNA Spots Identification Software, Seguin *et al*, 2014) and plotted with the TuYVs81 genome on the x-axis. For coverage plots of target genes, 18 to 26 nucleotide reads were mapped to TAIR10 genome only, no mismatches allowed, using bowtie1.2.3 (Langmead *et al*, 2009) and normalized in Count per million mapped reads (CPM) with a bin size =1 using deepTools version 3.3.0 (Ramírez *et al*, 2016). Reads from minus strand were converted into negative values, and the resulting bigWig files were visualized using JBrowse (Buels *et al*, 2016). Raw counts for TAIR10 loci (>10 counts across conditions) were used for differential analysis between mock and TuYVs81 libraries, Col-0 mock and TuYVs81 libraries, *ago1-57* mock and TuYVs81 libraries, ago1-57 and Col-0 libraries, either from total RNA of AGO1-purified sRNA using DEseq2 v1.12.4 (Love et al., 2014). Loci with an adjusted p value < 0.05 were considered as having differential sRNA accumulation as represented in the MAplot in **Figure S12B**. Heatmaps were generated from the top variable loci across either total RNA or AGO1 IP datasets, using the pheatmap R package. See **supplemental Table 1** for all DA loci.

### Primary vsiRNA target prediction

For vsiRNA-target pairing predictions, TuYVs81 matching vsiRNA counts (18 to 26-nt) were matched against the mRNA sequence of the genes identified by the differential analysis using psRNATarget V2 (Dai *et al*, 2018). Only regions in the transcript that contained a continuous population of matching vsiRNA in 5’ of secondary siRNA and with an expectation score ≤ 3.5 were considered. The same analysis was conducted from Col-0 total RNA library R1 and Col-0 AGO1 IP library R1 with identical regions delineated. Output files are **Supplemental Table 2 and 3**.

### Multiple sequence alignments and target region conservation analysis

Thirteen angiosperm species representing known polerovirus host species and relatives in the same clades were used for multiple sequence alignments. For Brassicaceae, *Arabidopsis thaliana*, *Arabidopsis lyrata*, *Eutrema salsugineum*, *Capsella rubella*, *Brassica rapa*, *Brassica oleracea, Brassica napus* were used, with *Theobroma cacao* and *Medicago truncatula* for the larger Rosid clade, *Solanum tuberosum* as representative for Asterids and host of PLRV, *Beta vulgaris* as representative for Amaranthaceae and host to several poleroviruses, as well as *Oryza sativa* and *Sorgum bicolor* as representative of monocotyledones. For each target identified in *A. thaliana*, orthologues sequences from angiosperms were mined using the interMine interface of Phytozome12 (https://phytozome.jgi.doe.gov/pz/portal.html). Orthologues in *Brassica napus* and *Beta vulgaris* were retrieved from EnsemblPlants (https://plants.ensembl.org/index.html). Nucleotide sequences were aligned using multiple sequence comparison by log-expectation web service (muscleWS) bundled with Jalview 2.11.1.3, with parameters set to default, and targeted regions were defined based on the ones predicted in *Arabidopsis thaliana*. In the event that more than one orthologue was identified in one species, all orthologues from that species were used for a first multiple alignment, and only ones that did not introduce indels in the target region was kept per alignment block, and the alignment was redone with the same parameters using only the preferred orthologue. All alignments were refined by removing species that introduced indels in the target region and realigning after removal. Codon triplets based on *A.thaliana* are indicated between dashed lines, and sequence logo for the considered regions were obtained using Weblogo3.

For polerovirus diversity, whole genomes of poleroviruses (https://www.ncbi.nlm.nih.gov/genomes/GenomesGroup.cgi?taxid=119164) were recovered from NCBI and aligned using muscleWS bundled with Jalview 2.11.1.3, with parameters set to default. Because of the extensive sequence variability observed for fast evolving polerovirus genomes, only turnip yellows virus (TuYV, NC_003743.1), cucurbit aphid-borne yellows virus (CaBYV, NC_003688.1), potato leafroll virus (PLRV, NC_001747.1) were used as representative of non-beet infecting poleroviruses, while beet mild yellowing virus (BMYV, NC_003491.1), beet chlorosis virus (BchV, NC_002766.1) and beet western yellows virus stain USA (BWYV-USA, AF473561.1) were used as beet-infecting representatives. An additional alignment containing brassica yellows virus (BrYV-ABJ NC_016038.2) was obtained for the MTERF/AT5G11000 targeting region located at the start of the subgenomic that has several extensions found only in TuYV and BrYV.

## Acknowledgements

The authors would like to thank Herve Vaucheret for the gift of *ago7-1*, *ago10-3* and multiple combinations containing *ago2-1* and *ago1-27* seeds. Todd Blevins and Christophe Himber at the IBMP for the gift of *ago5-1* and *dcl2-5/dcl3-1* seeds. Christophe Ritzenthaler at the IBMP for providing the Bentha B2-GFP plants. Brian G. Ayre at the University of north Texas for the pCoymv containing plasmid. Yasin Dagdas at the GMI for providing the pBin20 ER-CFP marker. We are grateful to all members of the Genschik lab, Benoit Derrien, Thibaut Hacquard, Nicolas Baumberger and Todd Blevins for fruitful discussions and Jerome Mutterer for assistance with microscopy. Pascal Genschik acknowledges support from the European Research Council under the European Union’s Seventh Framework Programme (FP7/2007-2013)/ERC advanced Grant 338904, and from the Agence Nationale de la Recherche LABEX, ANR-10-LABX-0036_NETRNA.

## Author contributions

Conceptualisation and design: Marion Clavel, Pascal Genschik. Data gathering, formal analysis, validation: Marion Clavel, Esther Lechner, Marco Incarbone, Timothée Vincent, Valerie Cognat, Ekaterina Smirnova, Maxime Lecorbeiller, Véronique Brault. Writing – original draft: Marion Clavel. Writing – review and editing: Marion Clavel, Esther Lechner, Véronique Ziegler-Graff and Pascal Genschik.

## Conflict of interest

The authors declare that they have no conflict of interest.

**Supplemental table 1:** Output table for DEseq2 pairwise comparisons, showing loci differentially accumulating sRNA reads. Pairwise comparisons for AGO1 IPs are the following: TuYVs81 *vs.* Mock all genotypes, *ago1-57 vs.* Col-0 all treatments, TuYVs81 *vs.* Mock Col-0, TuYVs81 *vs.* Mock *ago1*-*57*, TuYVs81 *ago1-57 vs.* TuYVs81 Col-0. Pairwise comparisons for total RNA are the following: TuYVs81 *vs.* mock all genotypes, TuYVs81 *vs.* mock Col-0, TuYVs81 *vs.* mock *ago1-57*. Rows colored in orange have a LFC>0, rows colored in blue have a LFC<0. Only annotations units with adjusted p-value<0.05 are shown.

**Supplemental table 2:** psRNATarget output table for the Col-0 TuYVs81 R1 library (JBT5) 18-26nt reads mapping only to the TuYVs81 genome (0mm) matched against the nine candidate target messengers. For each the region presumed to at the origin of the primary vsiRNA is colored in blue (expectation ≤ 3.5).

**Supplemental table 3:** psRNATarget output table for the Col-0 AGO1 IP TuYVs81 R1 library (JBT13) 18-26nt reads mapping only to the TuYVs81 genome (0mm) matched against the nine candidate target messengers. For each the region presumed to at the origin of the primary vsiRNA is colored in blue (expectation ≤ 3.5).

**Supplemental table 4:** Oligonucleotides used in this study.

---

**Supplemental Figure 1:**
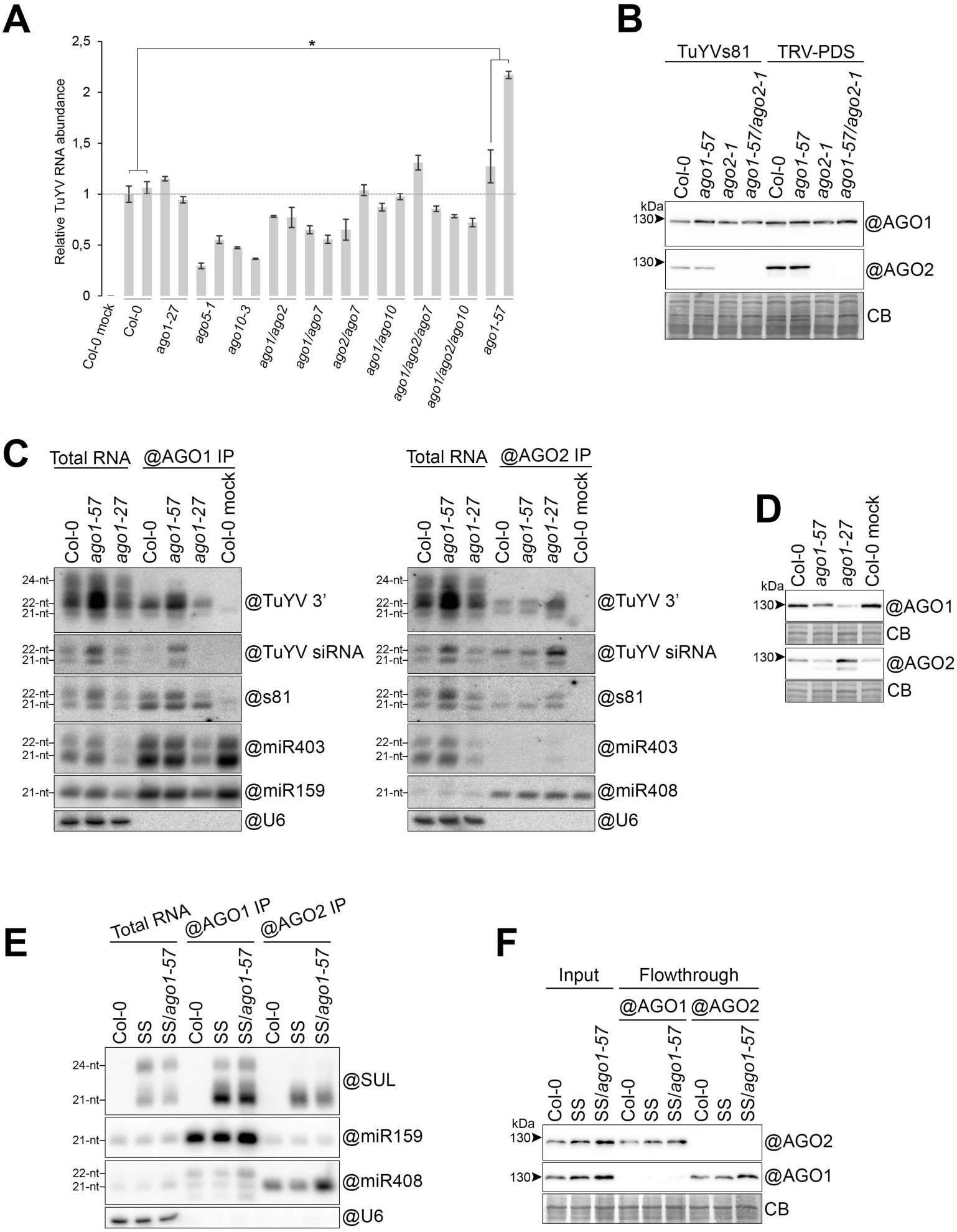
*ago1-57* is the only tested *argonaute* allele that affects TuYV RNA abundance (Supports Figure 1). **(A.)** TuYVs81 viral RNA abundance in systemic leaves of the indicated mutants at 21 dpi measured by RT-qPCR. Mock stands for mock-inoculated plants. Levels are displayed relative to infected Col-0. For each genotype, two independent samples were acquired from separate pools of infected plants to account for biological variability. Only leaves exhibiting yellow veins were harvested. *p value = 0.015371<0.05, with Student’s t-test, one-tailed, unequal variance. **(B.)** AGO1 and AGO2 protein abundance in systemic leaves of TuYVs81 and TRV-PDS inoculated plants at 20 dpi (n=5 plants), measured by immunoblot. Samples are input fractions of the AGO1 and AGO2 IPs presented in Figure 1J “@” indicates hybridization with antibody, and loading control is obtained by post-staining the membrane with Coomassie blue (CB). **(C.)** Analysis of AGO1 and AGO2-bound sRNA in WT, *ago1-57* and *ago1-27* TuYVs81 infected plants at 20 dpi (n=4). “@” indicates hybridization with DNA probe or use of a specific antibody for immunoprecipitation. Hybridization with miR159 (AGO1-bound) and miR408 (AGO2-bound) show efficient immunoprecipitation of the RISC in all samples. **(D.)** AGO1 and AGO2 protein abundance in mock and TuYVs81 systemic leaves measured by immunoblot. Samples are input fractions of the AGO1 and AGO2 IPs presented in **Figure S1C**. “@” indicates hybridization with antibody, and loading control is obtained by post-staining the membrane with Coomassie blue (CB). **(E.)** Analysis of AGO1 and AGO2-bound sRNA from rosette leaves of Col-0, SUC:SUL (SS), and SS/*ago1-57* plants. AGO1 binding of SS-derived siRNA is not affected by the *ago1-57* mutation nor does it lead to an increase of SS 21-nt siRNA in AGO2. “@” indicates hybridization with DNA probe or use of a specific antibody for immunoprecipitation. Hybridization with miR159 (AGO1-bound) and miR408 (AGO2-bound) show efficient immunoprecipitation of the RISC in all samples. **(F.)** AGO1 and AGO2 protein abundance in the rosette leaves used for the IP in **Figure S1E** before (input) and after (flowthrough) formation of the immune complexes. “@” indicates hybridization with antibody, and loading control is obtained by post-staining the membrane with Coomassie blue (CB).

**Supplemental Figure 2:**
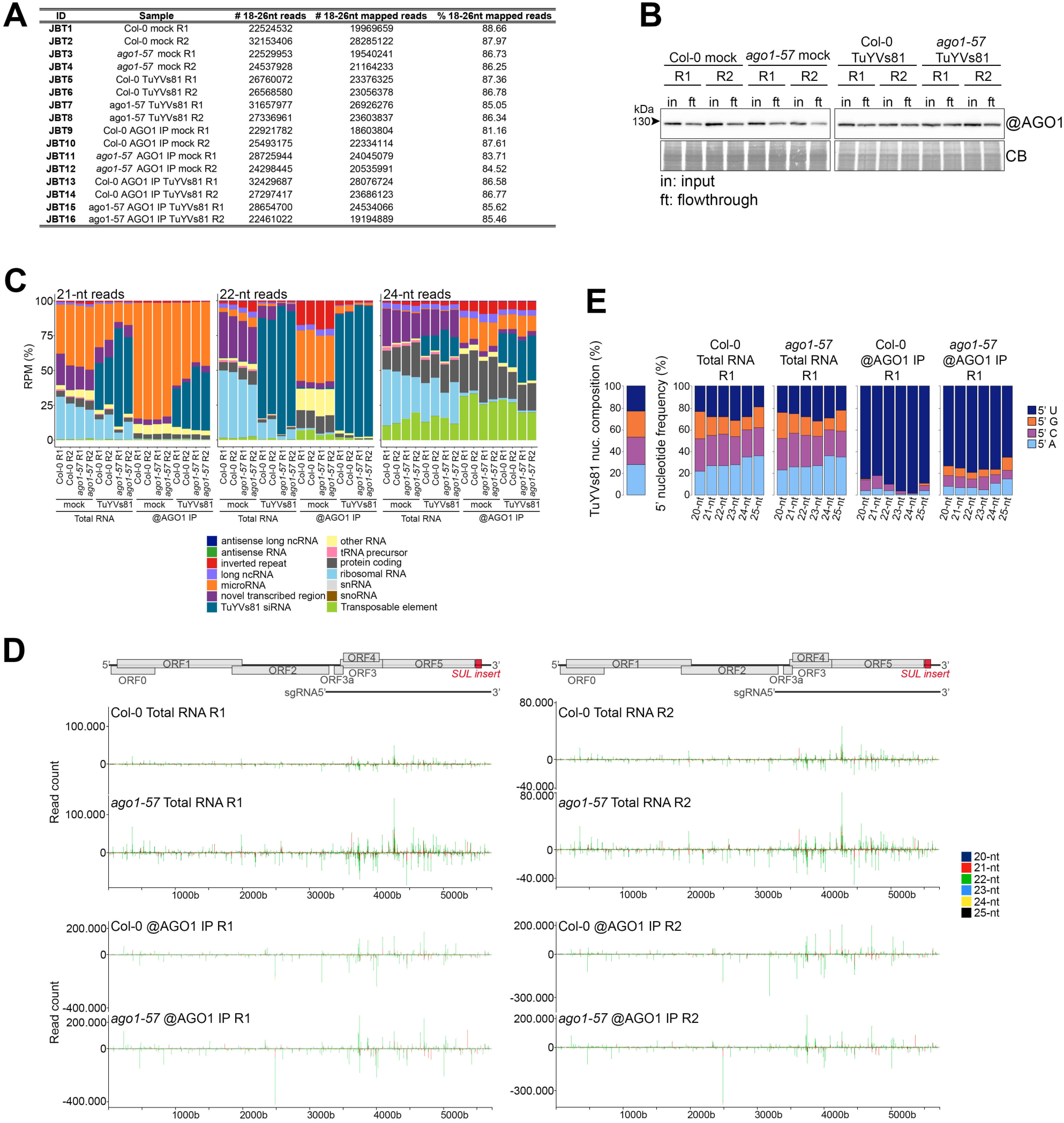
small RNA sequencing of TuYVs81-infected systemic leaves. **(A.)** Library-to-sample matching for the sixteen sRNA libraries obtained. For each library the number of mapped reads and percentage of mapped reads in the 18-26-nt interval is shown. Mapped reads were then used for further analysis and normalization. **(B.)** AGO1 protein abundance in mock-inoculated (mock) and TuYVs81 infected systemic leaves, measured by immunoblot, before (input) and after (flowthrough) formation of the immune complexes. Samples are from the AGO1 IPs sRNA deep sequencing experiment. “@” indicates hybridization with AGO1 antibody, and loading control is obtained by post-staining the membrane with Coomassie blue (CB). **(C.)** Percentage of normalized (RPM) sRNA reads aligned to the reference Arabidopsis genome and TuYVs81 genome per functional categories, broken down by size (21-nt, 22-nt and 24-nt). **(D.)** Distribution of TuYVs81-derived sRNA reads (20-nt to 25-nt) along the TuYVs81 genome in replicate 1 (R1, left panels) and replicate 2 (R2, right panel). Top panels compare Col-0 and *ago1-57* samples (fixed Y-axis) read distribution and abundance in total RNA samples. Bottom panels compare Col-0 and *ago1-57* samples (fixed Y-axis) read distribution and abundance in AGO1 immunoprecipitated samples. Bars indicate the position of the 5’ (+ strand) and 3’ (-strand) extremity of each mapped sRNA. Y-axis represents read counts, and each read size is represented in the indicated color. **(E.)** TuYVs81 nucleotide composition (%) and 5’ nucleotide frequency of the vsiRNA reads mapped to the TuYVs81 genome in the indicated libraries.

**Supplemental Figure 3:**
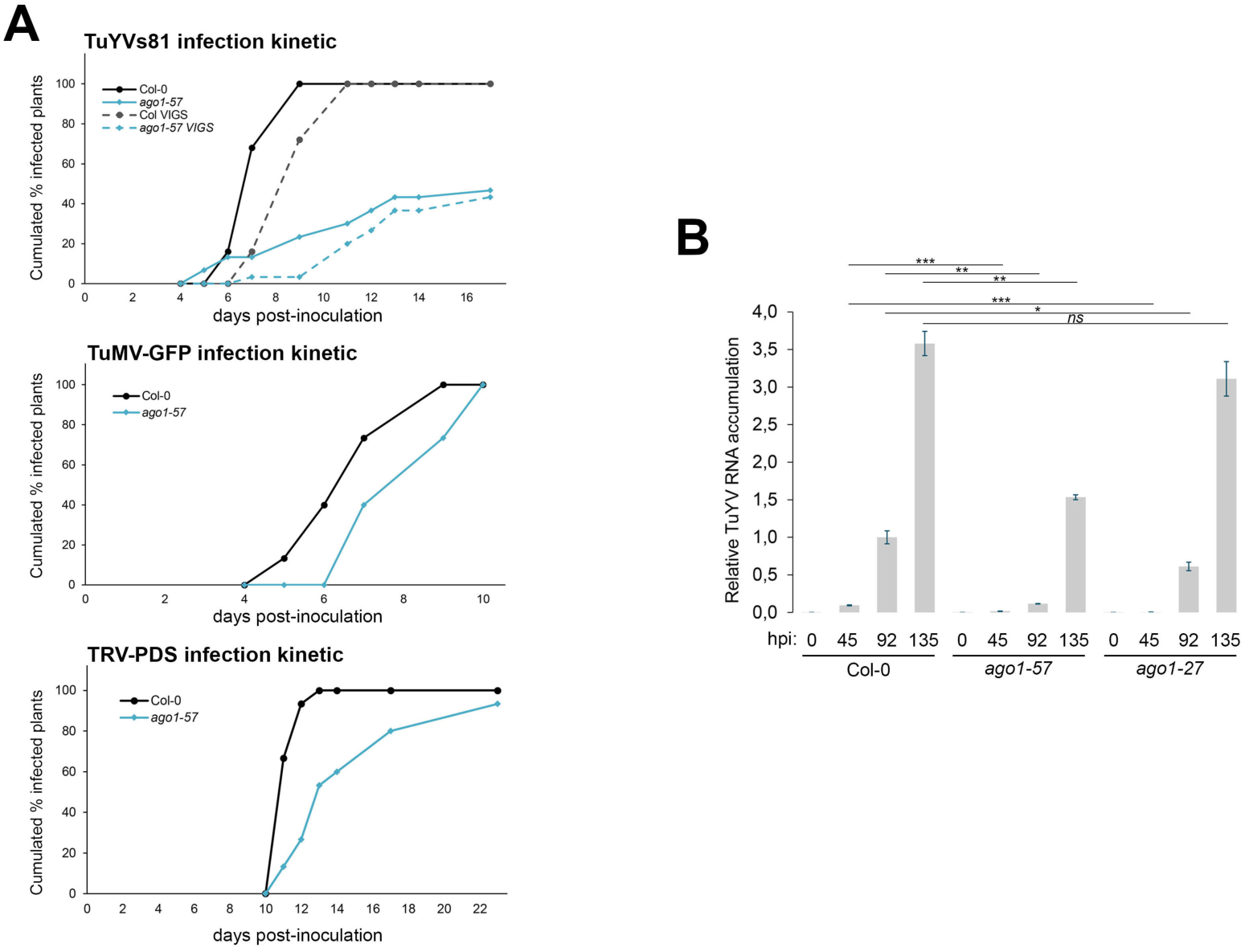
Unique interaction of TuYVs81 and *ago1-57* results in slower infection and lesser movement (supports Figure 2) **(A.)** Comparative infection kinetic between Col-0 and *ago1-57* for TuYVs81, TuMV-GFP and TRV-PDS. While TuYVs81 reaches 100% infection rate in the Col-0 plant population (n=25), only 43,3% of the *ago1-57* plants (n=30) result in a successful systemic infection (top panel). This is markedly different from either TuMV-GFP (middle panel, n=15 for both genotypes) or TRV-PDS infection (bottom panel, n=15 for both genotypes), in which *ago1-57* displays delayed infection but still reaches 100% systemic infection success. To avoid any confounding effect introduced by VIGS deficiency in the mutants, infected individuals were scored by detection of the TuYVs81 readthrough protein (RT, ORF5) in leaf patches from young systemic leaves at 5, 6, 7, 9, 11, 12, 14 and 17 dpi using western blot. Apparition of VIGS was counted separately and plotted as a dotted line. TuMV-GFP infection was scored by the detection of GFP in systemic leaves at the indicated day either using western blot on individual leaf patches or by illuminating the plants with a U.V lamp. TRV-PDS infection was scored by the detection of systemic leaf whitening at the indicated day. **(B.)** Accumulation of TuYVs81 RNA in inoculated leaves at 0, 45, 92, and 135 hours post infiltration (hpi) measured by RT-qPCR in the indicated genetic backgrounds represented as bar graph (left panel) or scatter plot along time (right panel), relative to Col-0 92hpi. Represented values are means of technical triplicates, error bars represent the SEM. For each time point, 1 infiltrated leaf from each individual (n=10 per genotype) was harvested, and plants were kept in the analyzed pool for the following time point. *p<0.05, **p<0.01, ***p<0.001 with Student’s t-test, one-tailed, paired.

**Supplemental Figure 4:**
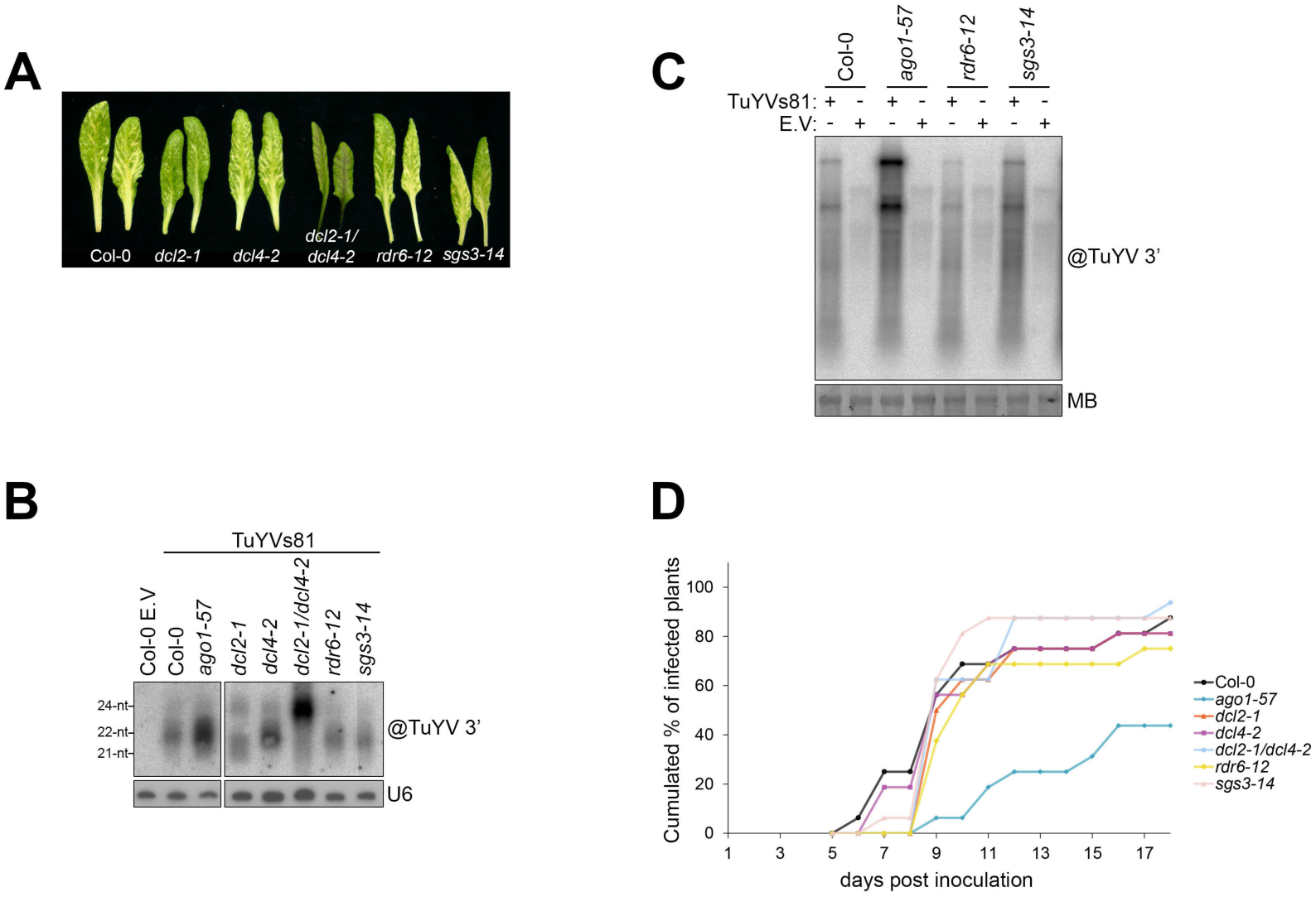
Impact of silencing mutants during TuYVs81 infections (supports Figure 3). **(A.)** Representative systemic infected leaves from the indicated genotypes. Pictures were taken at 17dpi from plants that scored positive for the virus at 9dpi. **(B.)** Analysis of vsiRNA accumulation in systemic leaves of the indicated genotypes at 22 dpi measured by RNA gel blot. Each lane represents a pool of four to ten individuals that scored positive for the presence of systemic TuYVs81 (via detection of the RT protein in leaf patches). “@” indicates hybridization with DNA probe against the 3’ part of the TuYV genome, U6 signal is the loading control. **(C.)** Accumulation of TuYVs81 RNA from the same samples as in **B**. Loading control is obtained by staining the membrane with methylene blue (MB). “@” indicates hybridization with DNA probe against the 3’ part of the TuYV genome. **(D.)** Kinetic of systemic TuYVs81 infection in the indicated genotypes represented as the cumulated percentage of infected plants in the inoculated population (n=16 individuals per genotype). Infected individuals were scored by the detection of the TuYVs8 RT protein in leaf patch from young systemic leaves at 9, 12 and 18 dpi by dot blot. Plants that exhibit systemic VIGS outside these sampling times are also counted as infected at the time point at which the VIGS was first observed. **(E.)** Top panel: Measurement of TuYVs81 RNA in inoculated leaves at 41, 60, 72 and 96 hour post infiltration (hpi) in the indicated genetic backgrounds measured by RNA gel blot. “@” indicates hybridization with TuYV 3’ probe and loading control is obtained by staining the membrane with methylene blue (MB). Bottom panel: Accumulation of TuYVs81 readthrough protein (RT) measured by immunoblot in equivalent samples. “@” indicates hybridization with RT antibody, loading control is obtained by post-staining the membrane with Coomassie blue (CB). For each time point, 5 infiltrated leaves from 4 to 5 individuals were harvested for each genotype.

**Supplemental Figure 5:**
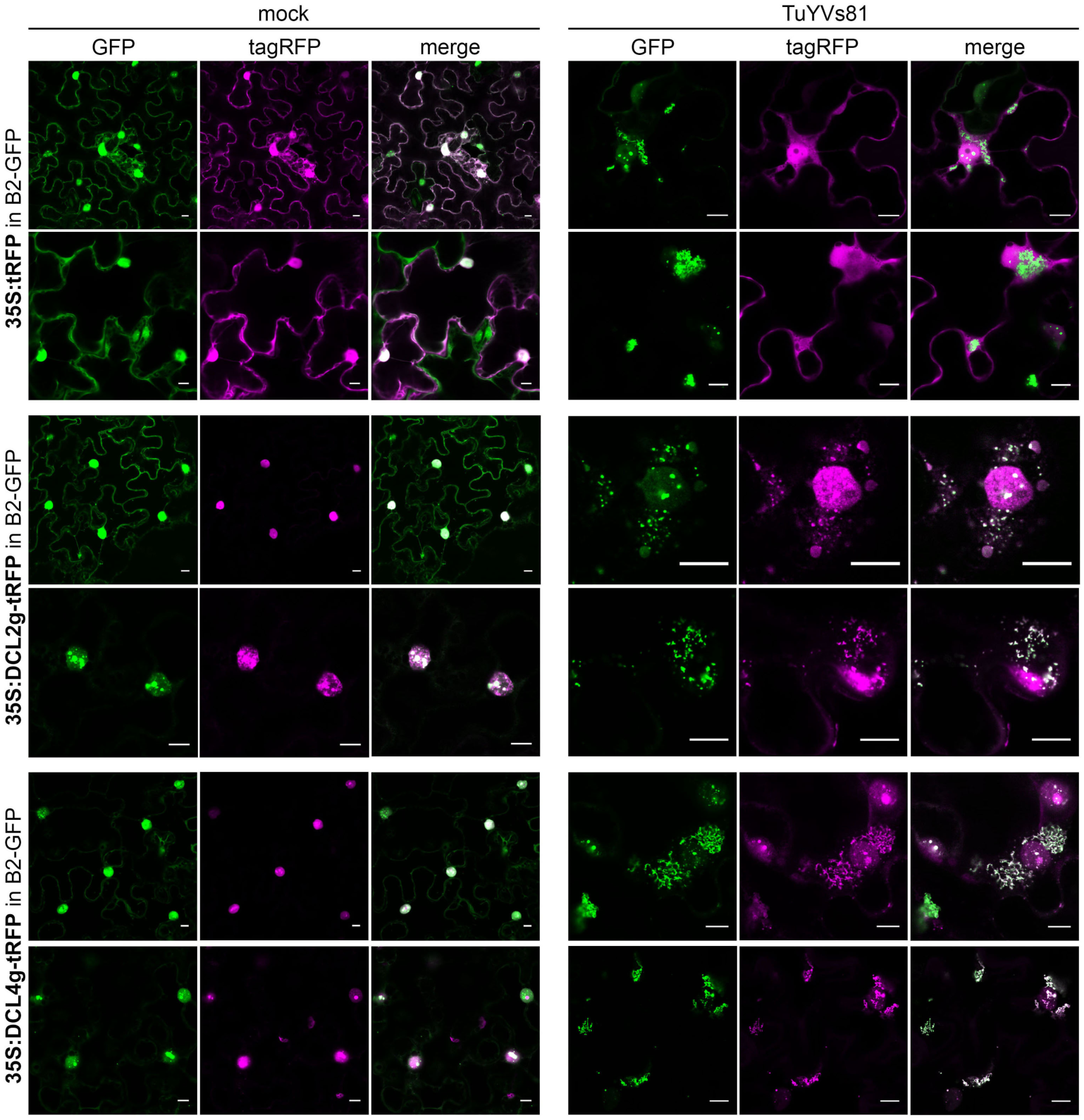
Localization of DCL2 and DCL4 upon TuYVs81 infection in *N. benthamiana* (supports Figure 3). Additional single plane confocal images of transiently expressed 35S:tRFP, 35S:DCL2genomic-tRFP and 35S:DCL4genomic-tRFP with (TuYVs81) or without (mock) the virus in leaves of transgenic *N. benthamiana* stably expressing the double-stranded RNA-binding B2-GFP protein. Observations are from leaf discs of 3 to 5 days post-infiltration. Inset scale bar is 10 µm.

**Supplemental Figure 6:**
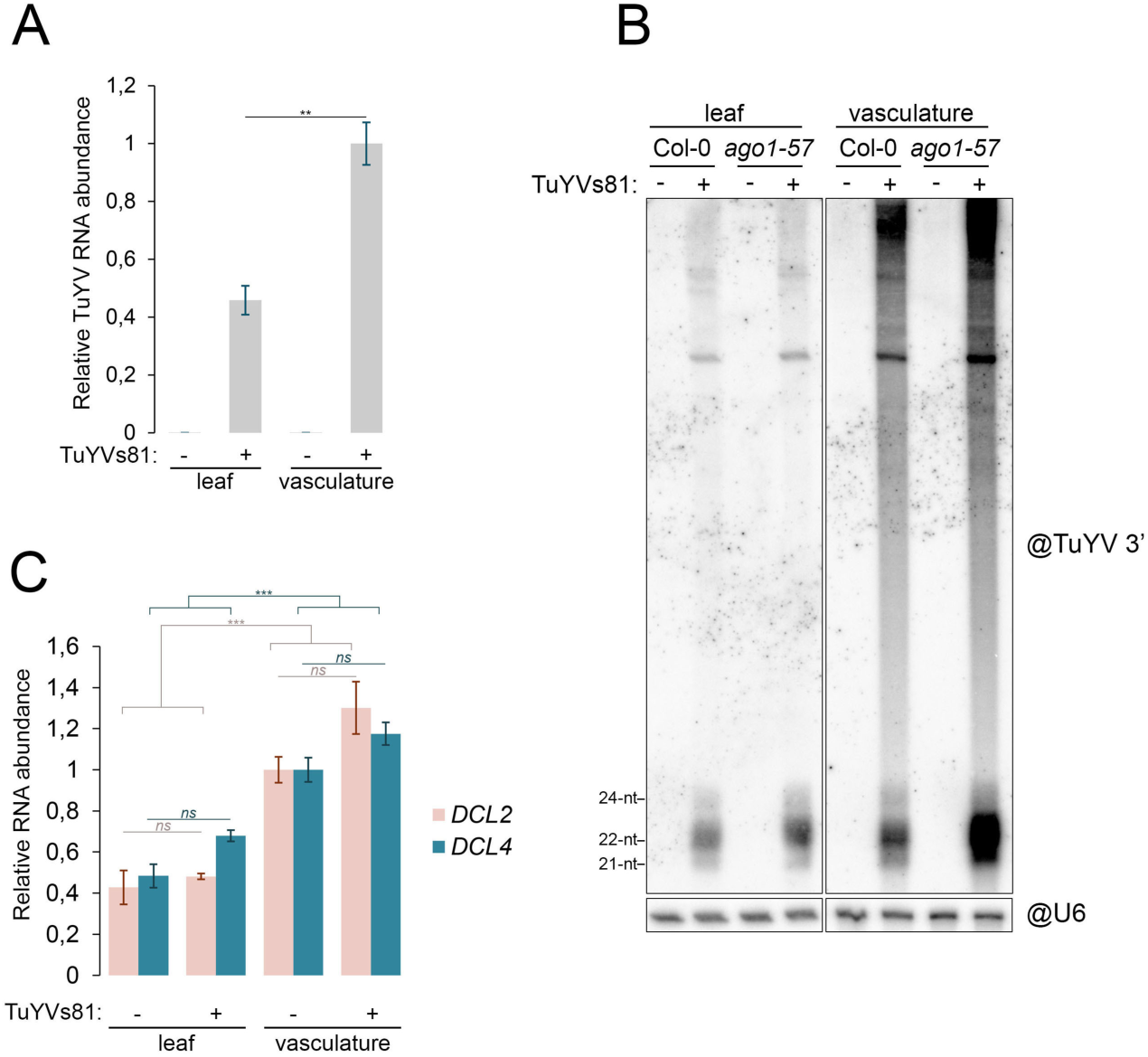
DCL2 expression is not elevated in TuYVs81-infected vasculatures (supports Figure 4) **(A.)** Measurement of TuYVs81 RNA in systemic whole leaves of Col-0 plants (n=12 individuals) or enriched vascular bundles of the equivalent plants (n=18 leaves) at 21 dpi by RT-qPCR represented as bar graph relative to infected vasculatures. Represented values are means of technical triplicates, error bars represent the SEM. **p<0.01, with Student’s t-test, one-tailed, paired. **(B.)** Uncropped RNA blot from Figure 4C. Abundant TuYVs81 replication intermediates/sRNA precursors are detected in enriched vascular tissues. “@” indicates hybridization with DNA probe and U6 signal is the loading control. **(C.)** Measurement of *DCL2* and *DCL4* mRNA abundance in the same samples as in A by RT-qPCR, represented as bar graph relative to - Col-0 vasculatures. Represented values are means of technical triplicates, error bars represent the SEM. ***p<0.001, *ns* p>0.05 with Student’s t-test, two-tailed, paired.

**Supplemental Figure 7:**
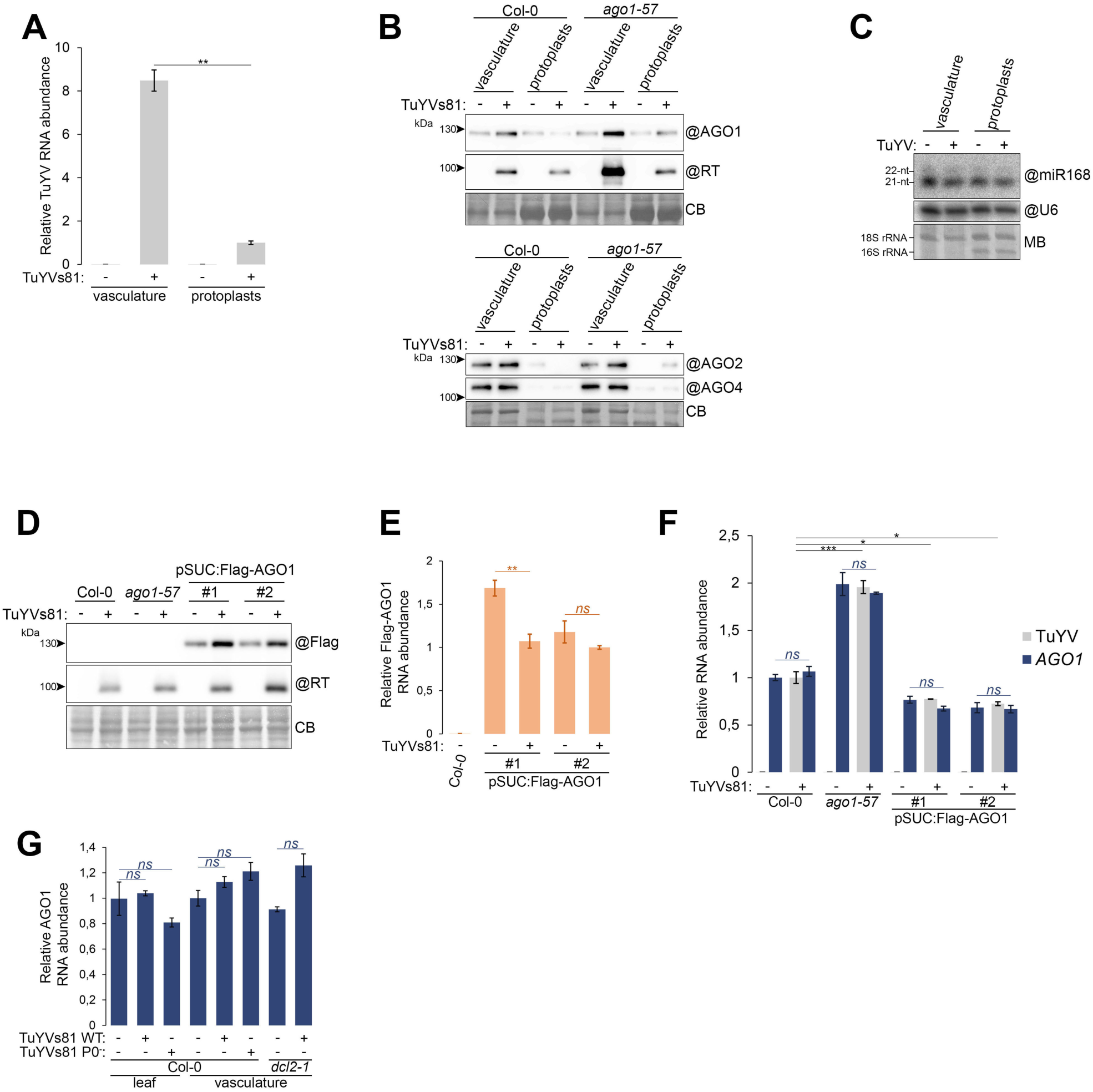
Stabilization of vascular AGO1 in presence of TuYVs81 (supports Figure 5). **(A.)** MeSelect was applied to enrich vascular bundles (vasculature), which were compared to the leaf mesophyll cells (protoplasts). Both were prepared from 10 leaves from a pool of 6 individuals of the indicated genotype at 15dpi. TuYVs81 RNA abundance was measured by RT-qPCR and is represented as a bar graph relative to the infected protoplasts. Values are means of technical triplicates, error bars represent the SEM. **p<0.01 with Student’s t-test, one-tailed, paired. **(B.)** The viral RT protein is enriched in vasculatures compared to protoplasts and AGO1 is the only tested ARGONAUTE protein to exhibit consistently increased abundance in response to the infection, in the three genotypes assayed. Protein samples are from the same experiment as in **A**. “@” indicates hybridization with the indicated antibodies, and loading control is obtained by post-staining the membrane with Coomassie blue. **(C.)** Abundance of vascular miR168 is not significantly affected by TuYVs81 infection as measured by RNA gel blot. RNA samples are the same as in **A**. “@” indicates hybridization with DNA probe and U6 signal is the loading control. The membrane was also stained with methylene blue (MB) which allows for visualization of cytoplasmic rRNA (here 18S) as well as chloroplastic 16S, that is absent from the vasculature. **(D.)** Phloem-restricted Flag-AGO1 is overaccumulated in TuYV infected leaves. Systemic whole leaves (n=6 individuals) at 21 dpi in Col-0, *ago1-57* and two transgenic lines expressing Flag-AGO1 from the SUC2 promoter in the absence (-) or presence (+) of TuYVs81. “@” indicates hybridization with the indicated antibodies, and loading control is obtained by post-staining the membrane with coomassie blue (CB). **(E.)** Accumulation of the *Flag-AGO1* transcript measured by RT-qPCR in the absence (-) or presence (+) of TuYVs81 in the indicated genetic backgrounds represented as bar graph. Represented values are means of technical triplicates, error bars represent the SEM. ns>0.05, **p<0.01 with Student’s t-test, two-tailed, paired. **(F.)** Accumulation of *AGO1* and TuYV RNA measured by RT-qPCR in the same samples as **D** and **E**. Represented values are means of technical triplicates, error bars represent the SEM. ns>0.05, **p<0.01, ***p<0.001 with Student’s t-test, two-tailed, paired for *AGO1* and two-tailed equal variance for TuYV. **(G.)** Levels of *AGO1* mRNA is not significantly affected by the presence of TuYVs81 WT or P0^-^ in the plant vasculature. mRNA abundance in the same samples as **Figure 4A & F** measured by RT-qPCR, represented as bar graph relative to - Col-0 vasculatures. Represented values are means of technical triplicates, error bars represent the SEM. *ns* p>0.05 with Student’s t-test, two-tailed, paired.

**Supplemental Figure 8:**
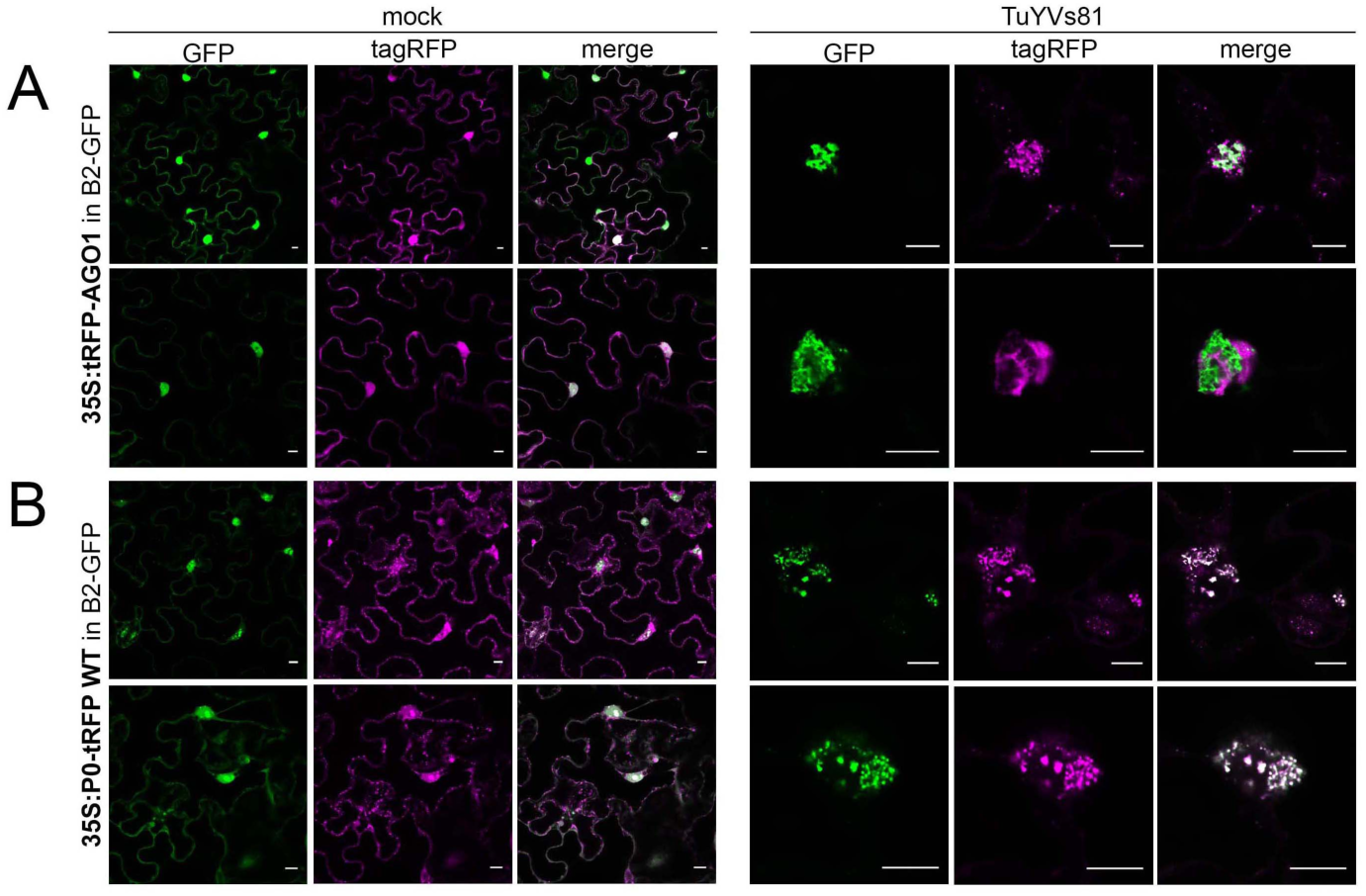
Localization of AGO1 and P0 upon TuYVs81 infection in *N. benthamiana* (supports Figure 5). Additional confocal images of transiently expressed **(A.)** 35S:tRFP-AGO1, **(B.)** 35S:P0-tRFP with (TuYVs81) or without (mock) the virus in leaves of transgenic *N. benthamiana* stably expressing the double-stranded RNA-binding B2-GFP protein. Observations are from leaf discs of 3 dpi. Inset scale bar is 10 µm.

**Supplemental Figure 9:**
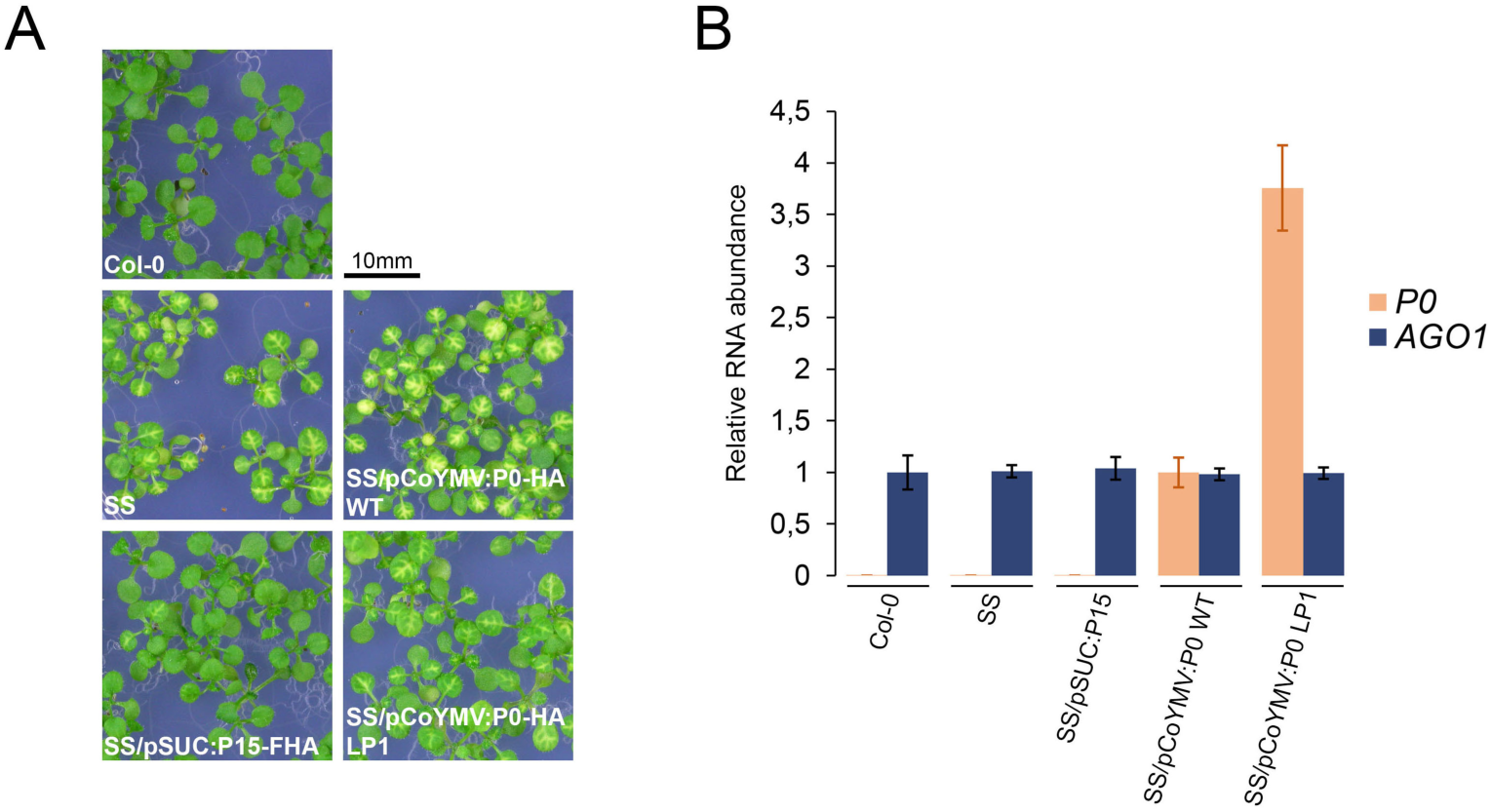
Phloem companion cell expression of P0 does not suppress cell-to-cell movement of *SUL* siRNA (supports Figure 5). **(A.)** *SUL*-silencing phenotype in SUC:SUL (SS), SS/pSUC:P15-FHA, SS/pCoYMV:P0 WT and LP1 mutant in 11 day-old seedlings grown on MS medium. **(B.)** Expression levels of transgenic P0 and endogenous AGO1 in the same seedlings (n=20 per sample) as in **A** by RT-qPCR represented as bar graphs. Represented values are means of technical triplicates, error bars represent the SEM.

**Supplemental figure 10:**
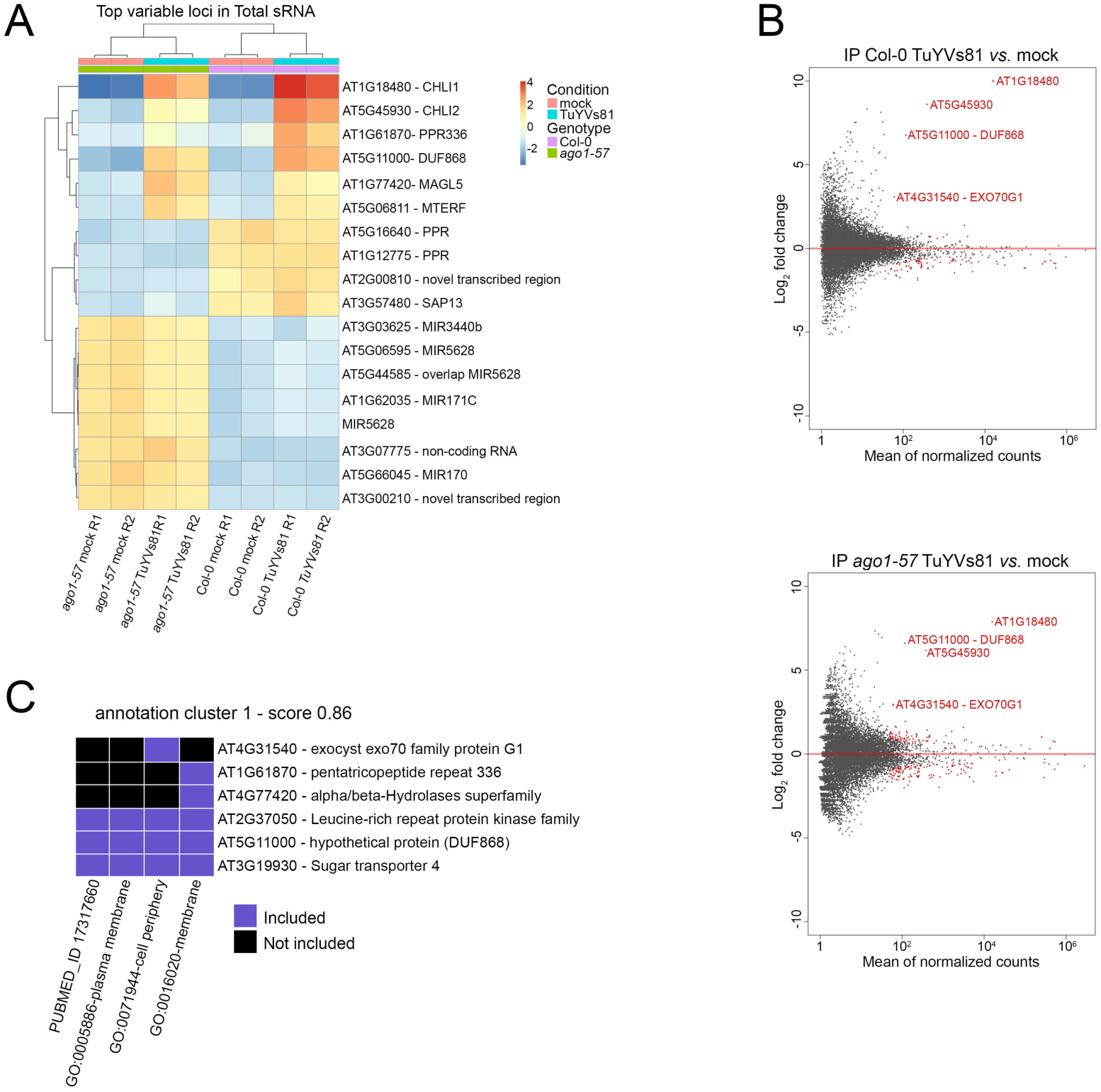
Production of vsiRNA-triggered secondary siRNA (supports Figure 6) **(A.)** Heatmap of annotation units showing the most variation in small RNA abundance across eight total RNA libraries. The first cluster contains genes that present enrichment of sRNA during TuYVs81 infection only, while the second and third clusters contain genes for which sRNA abundance if affected by the *ago1-57* mutation. **(B.)** Differential analysis of mock inoculated (mock) normalized sRNA reads compared to TuYVs81-infected normalized sRNA reads. Top panel: Normalized AGO1 IP sRNA libraries in Col-0. Bottom panel: Normalized AGO1 IP libraries in *ago1-57*. Abundance (mean of normalized counts) is displayed on the horizontal axis and log_2_ fold change on the vertical axis. Loci with an adjusted p-value lower than 0.05 are highlighted in red. **(C.)** Functional annotation clustering using DAVID 6.8 (2D view) for vsiRNA-targeted genes. Annotation cluster 1 (enrichment score 0.86) reveals enrichment for GO-terms related to membrane and cell periphery.

**Supplemental figure 11:**
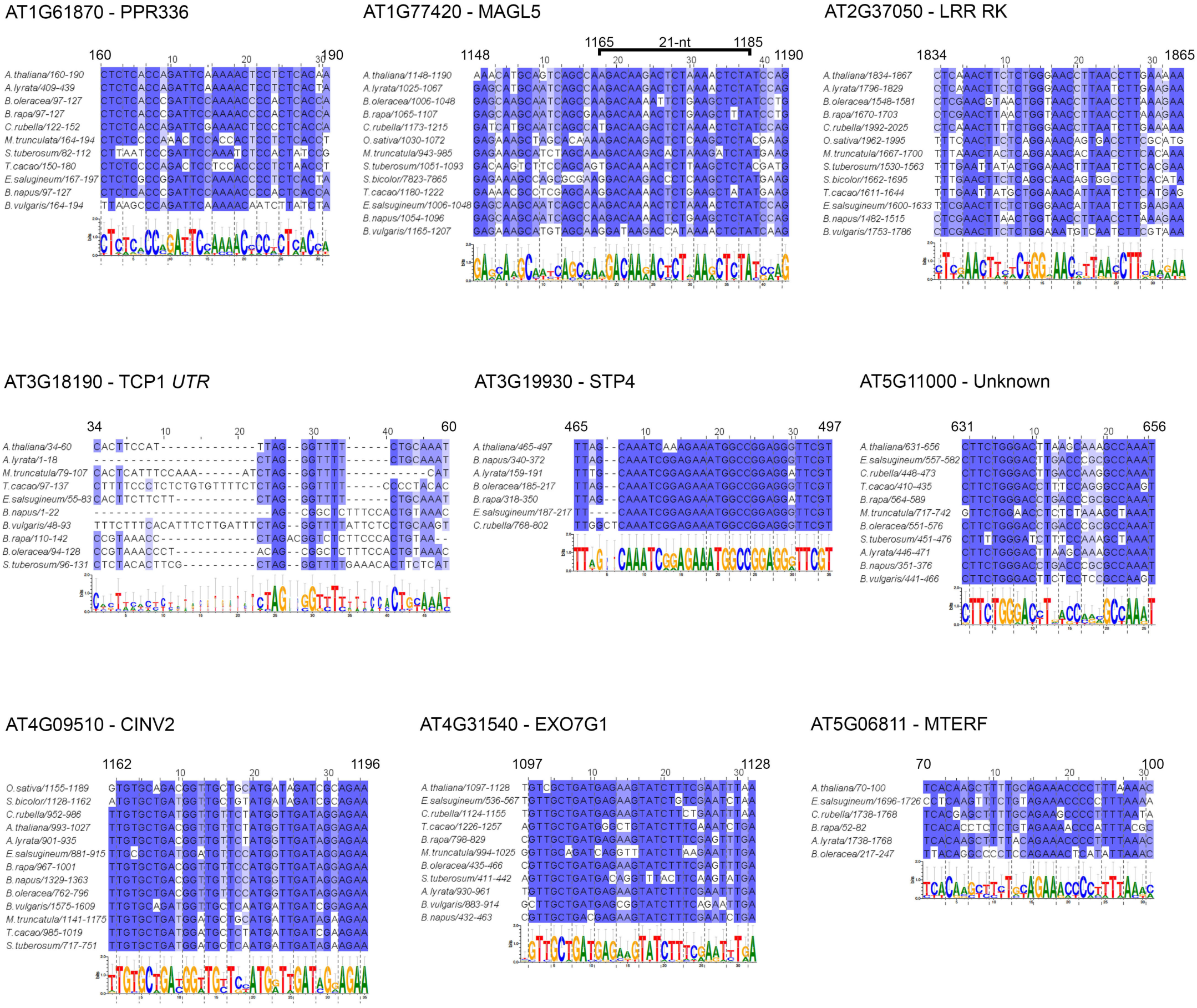
Alignment of TuYV vsiRNA targeted regions for the nine affected mRNA siRNA (supports Figure 6) For each candidate in Arabidopsis, orthologues sequences from angiosperms were found using the interMine interface of Phytozome12 (https://phytozome.jgi.doe.gov/pz/portal.html) for *Arabidopsis lyrata*, *Eutrema salsugineum*, *Capsella rubella*, *Brassica rapa*, *Brassica oleracea*, *Theobroma cacao*, *Medicago truncatula*, *Solanum tuberosum*, *Oryza sativa* and *Sorgum bicolor*. Orthologues in *Brassica napus* and *Beta vulgaris* were retrieved from EnsemblPlants (https://plants.ensembl.org/index.html). Nucleotide sequences were aligned using aligned using multiple sequence comparison by log-expectation web service (muscleWS) bundled with Jalview 2.11.1.3, with parameters set to default, and targeted regions were defined based on the ones predicted in *Arabidopsis thaliana*. Apart from TCP1, which is predicted to be targeted in the 5’UTR, all alignments were refined by removing species that introduced indels in the region of interest. Codon triplets based on *A. thaliana* are indicated between dashed lines, and sequence logo for the considered regions were obtained using Weblogo3. Conserved regions potentially targetable by TuYV are found for eight out of nine genes in at least Brassicaceae.

**Supplemental figure 12:**
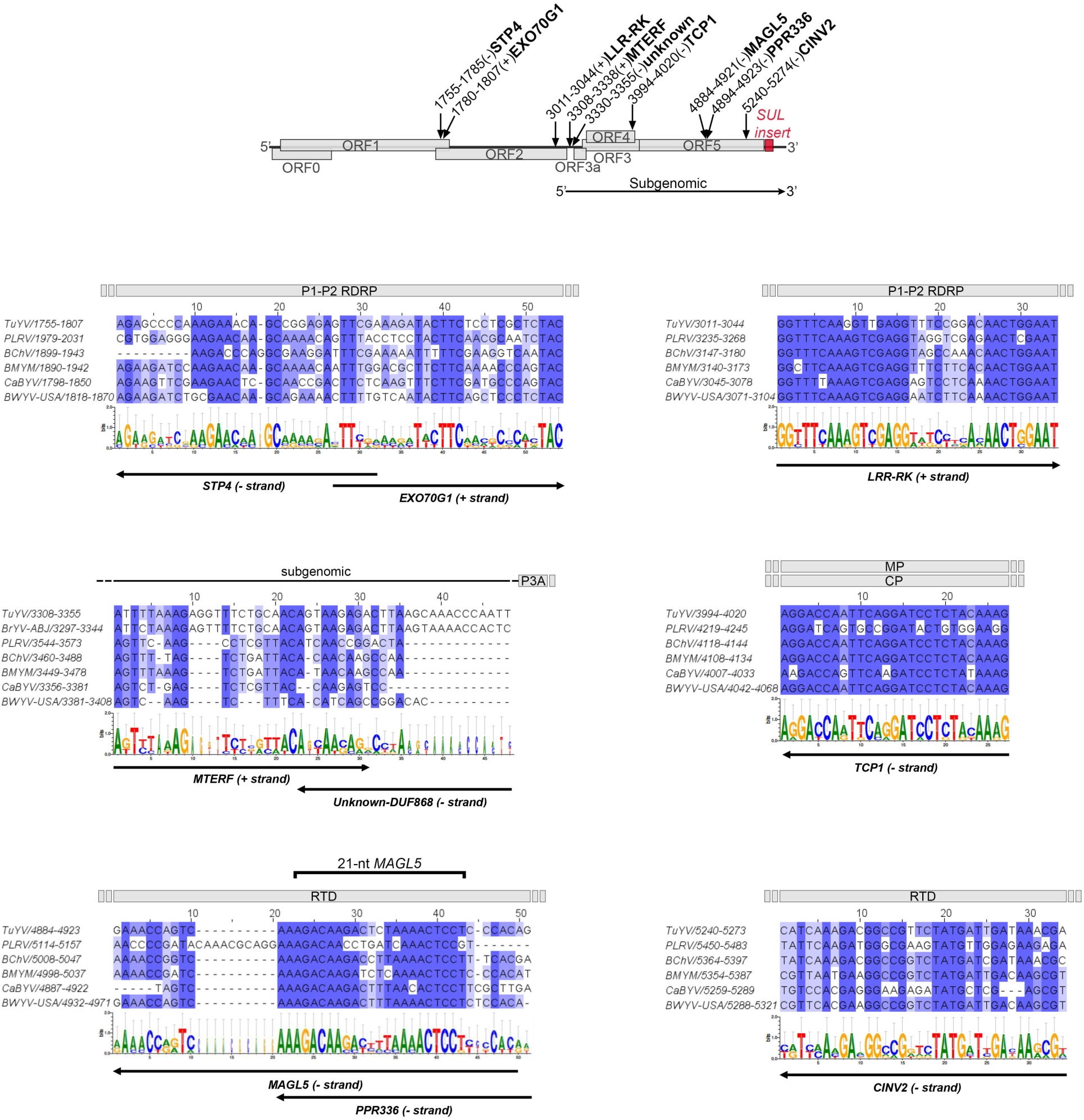
Alignment of Polerovirus targeting clusters siRNA (supports Figure 6) Six clusters for targeting sequences were defined based on the predicted regions in TuYVs81. Whole genomes of poleroviruses (https://www.ncbi.nlm.nih.gov/genomes/GenomesGroup.cgi?taxid=119164) were recovered from NCBI and aligned using muscleWS bundled with Jalview 2.11.1.3, with parameters set to default. Because of the extensive sequence variability observed for fast evolving polerovirus genomes, only turnip yellows virus (TuYV, NC_003743.1), cucurbit aphid-borne yellows virus (CaBYV, NC_003688.1), potato leafroll virus (PLRV, NC_001747.1) were used as representative of non-beet infecting poleroviruses, while beet mild yellowing virus (BMYV, NC_003491.1), beet chlorosis virus (BchV, NC_002766.1) and beet western yellows virus stain USA (BWYV-USA, AF473561.1) were used as beet-infecting representatives. An additional alignment containing brassica yellows virus (BrYV-ABJ NC_016038.2) was obtained for the MTERF/AT5G11000 targeting region located at the start of the subgenomic that has several extensions found only in TuYV and BrYV. Apart from the latter, all defined regions are conserved and could potentially serve as template for vsiRNA production targeting the nine candidate genes in the *Brassicaceae* family and beyond.

## References

1. Ahn HK, Yoon JT, Choi I, Kim S, Lee HS & Pai HS (2019) Functional characterization of chaperonin containing T-complex polypeptide-1 and its conserved and novel substrates in Arabidopsis. J. Exp. Bot. 70: 2741–2757

2. Almasi R, Miller WA & Ziegler-Graff V (2015) Mild and severe cereal yellow dwarf viruses differ in silencing suppressor efficiency of the P0 protein. Virus Res. 208: 199–206

3. Amin I, Hussain K, Akbergenov R, Yadav JS, Qazi J, Mansoor S, Hohn T, Fauquet CM & Briddon RW (2011) Suppressors of RNA Silencing Encoded by the Components of the Cotton Leaf Curl Begomovirus-BetaSatellite Complex. Mol. Plant-Microbe Interact. 24: 973–983 Available at: https://apsjournals.apsnet.org/doi/10.1094/MPMI-01-11-0001

4. Arribas-Hernández L, Marchais A, Poulsen C, Haase B, Hauptmann J, Benes V, Meister G & Brodersen P (2016) The Slicer Activity of ARGONAUTE1 is Required Specifically for the Phasing, Not Production, of Trans-Acting Short Interfering RNAs in Arabidopsis. Plant Cell 28: tpc.00121.2016 Available at: http://www.plantcell.org/lookup/doi/10.1105/tpc.16.00121

5. Azevedo J, Garcia D, Pontier D, Ohnesorge S, Yu A, Garcia S, Braun L, Bergdoll M, Hakimi MA, Lagrange T & Voinnet O (2010) Argonaute quenching and global changes in Dicer homeostasis caused by a pathogen-encoded GW repeat protein. Genes Dev. 24: 904–915

6. Baumberger N, Tsai C-H, Lie M, Havecker E & Baulcombe DC (2007) The Polerovirus Silencing Suppressor P0 Targets ARGONAUTE Proteins for Degradation. Curr. Biol. 17: 1609–1614 Available at: https://linkinghub.elsevier.com/retrieve/pii/S0960982207018568

7. Bayne EH, Rakitina D V., Morozov SY & Baulcombe DC (2005) Cell-to-cell movement of Potato Potexvirus X is dependent on suppression of RNA silencing. Plant J. 44: 471–482

8. Blevins T, Rajeswaran R, Shivaprasad P V., Beknazariants D, Si-Ammour A, Park HS, Vazquez F, Robertson D, Meins F, Hohn T & Pooggin MM (2006) Four plant Dicers mediate viral small RNA biogenesis and DNA virus induced silencing. Nucleic Acids Res. 34: 6233–6246

9. Bohmert K, Camus I, Bellini C, Bouchez D, Caboche M, Benning C & Banning C (1998) AGO1 defines a novel locus of Arabidopsis controlling leaf development. EMBO J. 17: 170–80

10. Bortolamiol-Bécet D, Monsion B, Chapuis S, Hleibieh K, Scheidecker D, Alioua A, Bogaert F, Revers F, Brault V & Ziegler-Graff V (2018) Phloem-Triggered Virus-Induced Gene Silencing Using a Recombinant Polerovirus. Front. Microbiol. 9: 1–15 Available at: https://www.frontiersin.org/article/10.3389/fmicb.2018.02449/full

11. Bortolamiol D, Pazhouhandeh M, Marrocco K, Genschik P & Ziegler-Graff V (2007) The Polerovirus F Box Protein P0 Targets ARGONAUTE1 to Suppress RNA Silencing. Curr. Biol. 17: 1615–1621 Available at: https://linkinghub.elsevier.com/retrieve/pii/S0960982207017782

12. Bouché N, Lauressergues D, Gasciolli V & Vaucheret H (2006) An antagonistic function for Arabidopsis DCL2 in development and a new function for DCL4 in generating viral siRNAs. EMBO J. 25: 3347–3356

13. Brosseau C & Moffett P (2015) Functional and Genetic Analysis Identify a Role for Arabidopsis ARGONAUTE5 in Antiviral RNA Silencing. Plant Cell 27: 1742–1754 Available at: http://www.plantcell.org/lookup/doi/10.1105/tpc.15.00264

14. Buels R, Yao E, Diesh CM, Hayes RD, Munoz-Torres M, Helt G, Goodstein DM, Elsik CG, Lewis SE, Stein L & Holmes IH (2016) JBrowse: a dynamic web platform for genome visualization and analysis. Genome Biol. 17: 66 Available at: http://dx.doi.org/10.1186/s13059-016-0924-1

15. Carbonell A & Carrington JC (2015) Antiviral roles of plant ARGONAUTES. Curr. Opin. Plant Biol. 27: 111–117 Available at: http://dx.doi.org/10.1016/j.pbi.2015.06.013

16. Carbonell A, Fahlgren N, Garcia-Ruiz H, Gilbert KB, Montgomery TA, Nguyen T, Cuperus JT & Carrington JC (2012) Functional Analysis of Three Arabidopsis ARGONAUTES Using Slicer-Defective Mutants. Plant Cell 24: 3613–3629 Available at: http://www.plantcell.org/cgi/doi/10.1105/tpc.112.099945

17. Cascardo RS, Arantes ILG, Silva TF, Sachetto-Martins G, Vaslin MFS & Corrêa RL (2015) Function and diversity of P0 proteins among cotton leafroll dwarf virus isolates. Virol. J. 12: 123 Available at: http://dx.doi.org/10.1186/s12985-015-0356-7

18. Chen HM, Chen LT, Patel K, Li YH, Baulcombe DC & Wu SH (2010) 22-Nucleotide RNAs trigger secondary siRNA biogenesis in plants. Proc. Natl. Acad. Sci. U. S. A. 107: 15269–15274

19. Chen S, Jiang G, Wu J, Liu Y, Qian Y & Zhou X (2016) Characterization of a Novel Polerovirus Infecting Maize in China. Viruses 8: 120 Available at: http://www.mdpi.com/1999-4915/8/5/120

20. Cheng C-Y, Krishnakumar V, Chan AP, Thibaud-Nissen F, Schobel S & Town CD (2017) Araport11: a complete reannotation of the Arabidopsis thaliana reference genome. Plant J. 89: 789–804 Available at: http://doi.wiley.com/10.1111/tpj.13415

21. Chiba S, Hleibieh K, Delbianco A, Klein E, Ratti C, Ziegler-Graff V, Bouzoubaa S & Gilmer D (2013) The Benyvirus RNA silencing suppressor is essential for long-distance movement, requires both zinc-finger and nols basic residues but not a nucleolar localization for its silencing-suppression activity. Mol. Plant-Microbe Interact. 26: 168–181

22. Chong YT, Gidda SK, Sanford C, Parkinson J, Mullen RT & Goring DR (2010) Characterization of the Arabidopsis thaliana exocyst complex gene families by phylogenetic, expression profiling, and subcellular localization studies. New Phytol. 185: 401–419

23. Chung T, Phillips AR & Vierstra RD (2010) ATG8 lipidation and ATG8-mediated autophagy in Arabidopsis require ATG12 expressed from the differentially controlled ATG12A AND ATG12B loci. Plant J. 62: 483–493 Available at: http://doi.wiley.com/10.1111/j.1365-313X.2010.04166.x

24. Clark MF & Adams AN (1977) Characteristics of the Microplate Method of Enzyme-Linked Immunosorbent Assay for the Detection of Plant Viruses. J. Gen. Virol. 34: 475–483 Available at: https://www.microbiologyresearch.org/content/journal/jgv/10.1099/0022-1317-34-3-475

25. Clough SJ & Bent AF (1998) Floral dip: A simplified method for Agrobacterium-mediated transformation of *Arabidopsis thaliana*. Plant J. 16: 735–743

26. Cuperus JT, Carbonell A, Fahlgren N, Garcia-Ruiz H, Burke RT, Takeda A, Sullivan CM, Gilbert SD, Montgomery TA & Carrington JC (2010) Unique functionality of 22-nt miRNAs in triggering RDR6-dependent siRNA biogenesis from target transcripts in Arabidopsis. Nat. Struct. Mol. Biol. 17: 997–1003

27. Dai X, Zhuang Z & Zhao PX (2018) psRNATarget: a plant small RNA target analysis server (2017 release). Nucleic Acids Res. 46: W49–W54 Available at: https://academic.oup.com/nar/article/46/W1/W49/4990032

28. Deleris A, Gallago-Bartolome J, Bao J, Kasschau KD, Carrington JC & Voinnet O (2006) Hierarchical action and inhibition of plant dicer-like proteins in antiviral defense. Science (80-.). 313: 68–71

29. Delfosse VC, Agrofoglio YC, Casse MF, Kresic IB, Hopp HE, Ziegler-Graff V & Distéfano AJ (2014) The P0 protein encoded by cotton leafroll dwarf virus (CLRDV) inhibits local but not systemic RNA silencing. Virus Res. 180: 70–75 Available at: http://dx.doi.org/10.1016/j.virusres.2013.12.018

30. Deng Y, Wang J, Tung J, Liu D, Zhou Y, He S, Du Y, Baker B & Li F (2018) A role for small RNA in regulating innate immunity during plant growth. PLOS Pathog. 14: e1006756 Available at: http://journals.plos.org/plospathogens/article/file?id=10.1371/journal.ppat.1006756&type=printable

31. Derrien B, Baumberger N, Schepetilnikov M, Viotti C, De Cillia J, Ziegler-Graff V, Isono E, Schumacher K & Genschik P (2012) Degradation of the antiviral component ARGONAUTE1 by the autophagy pathway. Proc. Natl. Acad. Sci. 109: 15942–15946 Available at: http://www.pnas.org/cgi/doi/10.1073/pnas.1209487109

32. Derrien B, Clavel M, Baumberger N, Iki T, Sarazin A, Hacquard T, Ponce MR, Ziegler-Graff V, Vaucheret H, Micol JL, Voinnet O & Genschik P (2018) A Suppressor Screen for AGO1 Degradation by the Viral F-Box P0 Protein Uncovers a Role for AGO DUF1785 in sRNA Duplex Unwinding. Plant Cell 30: 1353–1374 Available at: http://www.plantcell.org/lookup/doi/10.1105/tpc.18.00111

33. Devers EA, Brosnan CA, Sarazin A, Albertini D, Amsler AC, Brioudes F, Jullien PE, Lim P, Schott G & Voinnet O (2020) Movement and differential consumption of short interfering RNA duplexes underlie mobile RNA interference. Nat. plants 6: Available at: http://www.ncbi.nlm.nih.gov/pubmed/32632272

34. Diaz-Pendon JA, Li F, Li W-X & Ding S-W (2007) Suppression of Antiviral Silencing by Cucumber Mosaic Virus 2b Protein in Arabidopsis Is Associated with Drastically Reduced Accumulation of Three Classes of Viral Small Interfering RNAs. Plant Cell 19: 2053–2063 Available at: http://www.plantcell.org/lookup/doi/10.1105/tpc.106.047449

35. Donaire L, Barajas D, Martinez-Garcia B, Martinez-Priego L, Pagan I & Llave C (2008) Structural and Genetic Requirements for the Biogenesis of Tobacco Rattle Virus-Derived Small Interfering RNAs. J. Virol. 82: 5167–5177 Available at: http://jvi.asm.org/cgi/doi/10.1128/JVI.00272-08

36. Donaire L, Wang Y, Gonzalez-Ibeas D, Mayer KF, Aranda MA & Llave C (2009) Deep-sequencing of plant viral small RNAs reveals effective and widespread targeting of viral genomes. Virology 392: 203–214 Available at: http://dx.doi.org/10.1016/j.virol.2009.07.005

37. Fusaro AF, Correa RL, Nakasugi K, Jackson C, Kawchuk L, Vaslin MFS & Waterhouse PM (2012) The Enamovirus P0 protein is a silencing suppressor which inhibits local and systemic RNA silencing through AGO1 degradation. Virology 426: 178–187 Available at: http://dx.doi.org/10.1016/j.virol.2012.01.026

38. Garcia-Ruiz H, Carbonell A, Hoyer JS, Fahlgren N, Gilbert KB, Takeda A, Giampetruzzi A, Garcia Ruiz MT, McGinn MG, Lowery N, Martinez Baladejo MT & Carrington JC (2015) Roles and Programming of Arabidopsis ARGONAUTE Proteins during Turnip Mosaic Virus Infection. PLoS Pathog. 11: 1–27 Available at: http://dx.doi.org/10.1371/journal.ppat.1004755

39. Garcia-Ruiz H, Takeda A, Chapman EJ, Sullivan CM, Fahlgren N, Brempelis KJ & Carrington JC (2010) Arabidopsis RNA-dependent RNA polymerases and dicer-like proteins in antiviral defense and small interfering RNA biogenesis during Turnip mosaic virus infection. Plant Cell 22: 481–496

40. Gregory BD, O’Malley RC, Lister R, Urich MA, Tonti-Filippini J, Chen H, Millar AH & Ecker JR (2008) A Link between RNA Metabolism and Silencing Affecting Arabidopsis Development. Dev. Cell 14: 854–866 Available at: https://linkinghub.elsevier.com/retrieve/pii/S1534580708001688

41. Guo HS & Ding SW (2002) A viral protein inhibits the long range signaling activity of the gene silencing signal. EMBO J. 21: 398–407 Available at: http://emboj.embopress.org/cgi/doi/10.1093/emboj/21.3.398

42. Han YH, Xiang HY, Wang Q, Li YY, Wu WQ, Han CG, Li DW & Yu JL (2010) Ring structure amino acids affect the suppressor activity of melon aphid-borne yellows virus P0 protein. Virology 406: 21–27 Available at: http://dx.doi.org/10.1016/j.virol.2010.06.045

43. Harvey JJW, Lewsey MG, Patel K, Westwood J, Heimstädt S, Carr JP & Baulcombe DC (2011) An antiviral defense role of AGO2 in plants. PLoS One 6: 1–6

44. Havelda Z, Hornyik C, Crescenzi A & Burgyan J (2003) In Situ Characterization of Cymbidium Ringspot Tombusvirus Infection-Induced Posttranscriptional Gene Silencing in Nicotiana benthamiana. J. Virol. 77: 6082–6086

45. Haxim Y, Ismayil A, Jia Q, Wang Y, Zheng X, Chen T, Qian L, Liu N, Wang Y, Han S, Cheng J, Qi Y, Hong Y & Liu Y (2017) Autophagy functions as an antiviral mechanism against geminiviruses in plants. Elife 6: 1–17 Available at: https://elifesciences.org/articles/23897

46. Herrbach E, Lemaire O, Ziegler-Graff V, Lot H, Rabenstein F & Bouchery Y (1991) Detection of BMYV and BWYV isolates using monoclonal antibodies and radioactive RNA probes, and relationships among luteoviruses. Ann. Appl. Biol. 118: 127–138 Available at: https://onlinelibrary.wiley.com/doi/abs/10.1111/j.1744-7348.1991.tb06091.x

47. Himber C, Dunoyer P, Moissiard G, Ritzenthaler C & Voinnet O (2003) Transitivity-dependent and -independent cell-to-cell movement of RNA silencing. EMBO J. 22: 4523–4533

48. Huang DW, Sherman BT & Lempicki RA (2009) Systematic and integrative analysis of large gene lists using DAVID bioinformatics resources. Nat. Protoc. 4: 44–57 Available at: http://www.nature.com/articles/nprot.2008.211

49. Imlau A, Truernit E & Sauer N (1999) Cell-to-Cell and Long-Distance Trafficking of the Green Fluorescent Protein in the Phloem and Symplastic Unloading of the Protein into Sink Tissues. Plant Cell 11: 309–322 Available at: http://www.plantcell.org/lookup/doi/10.1105/tpc.11.3.309

50. Incarbone M, Clavel M, Monsion B, Kuhn L, Scheer H, Poignavent V, Dunoyer P, Genschik P & Ritzenthaler C (2020) Immunocapture of dsRNA-bound proteins provides insight into tobacco rattle virus replication complexes and reveals Arabidopsis DRB2 to be a wide-spectrum antiviral effector. bioRxiv

51. Incarbone M & Dunoyer P (2013) RNA silencing and its suppression: Novel insights from in planta analyses. Trends Plant Sci. 18: 382–392 Available at: http://dx.doi.org/10.1016/j.tplants.2013.04.001

52. Incarbone M, Zimmermann A, Hammann P, Erhardt M, Michel F & Dunoyer P (2017) Neutralization of mobile antiviral small RNA through peroxisomal import. Nat. Plants 3: Available at: http://dx.doi.org/10.1038/nplants.2017.94

53. Ismayil A, Yang M, Haxim Y, Wang Y, Li J, Han L, Wang Y, Zheng X, Wei X, Nagalakshmi U, Hong Y, Hanley-Bowdoin L & Liu Y (2020) Cotton leaf curl Multan virus βC1 Protein Induces Autophagy by Disrupting the Interaction of Autophagy-Related Protein 3 with Glyceraldehyde-3-Phosphate Dehydrogenases. Plant Cell 32: 1124–1135 Available at: https://academic.oup.com/plcell/article/32/4/1124-1135/6115663

54. Jaubert M, Bhattacharjee S, Mello AFS, Perry KL & Moffett P (2011) ARGONAUTE2 Mediates RNA-Silencing Antiviral Defenses against Potato virus X in Arabidopsis. Plant Physiol. 156: 1556–1564 Available at: http://www.plantphysiol.org/lookup/doi/10.1104/pp.111.178012

55. Johnson NR, Yeoh JM, Coruh C & Axtell MJ (2016) Improved Placement of Multi-mapping Small RNAs. G3 6: 2103–2111 Available at: https://academic.oup.com/g3journal/article/6/7/2103-2111/6027713

56. Juszczuk M, Paczkowska E, Sadowy E, Zagórski W & Hulanicka DM (2000) Effect of genomic and subgenomic leader sequences of potato leafroll virus on gene expression. FEBS Lett. 484: 33–36

57. Kim RJ, Kim HJ, Shim D & Suh MC (2016) Molecular and biochemical characterizations of the monoacylglycerol lipase gene family of Arabidopsis thaliana. Plant J. 85: 758–771 Available at: http://doi.wiley.com/10.1111/tpj.13146

58. Klein E, Brault V, Klein D, Weyens G, Lefèbvre M, Ziegler-Graff V & Gilmer D (2014) Divergence of host range and biological properties between natural isolate and full-length infectious cDNA clone of the Beet mild yellowing virus 2ITB. Mol. Plant Pathol. 15: 22–30 Available at: https://onlinelibrary.wiley.com/doi/abs/10.1111/mpp.12061

59. Kleine T (2012) Arabidopsis thaliana mTERF proteins: evolution and functional classification. Front. Plant Sci. 3: 1–15 Available at: http://journal.frontiersin.org/article/10.3389/fpls.2012.00233/abstract

60. Kozlowska-Makulska A, Guilley H, Szyndel MS, Beuve M, Lemaire O, Herrbach E & Bouzoubaa S (2010) P0 proteins of European beet-infecting poleroviruses display variable RNA silencing suppression activity. J. Gen. Virol. 91: 1082–1091

61. Langmead B, Trapnell C, Pop M & Salzberg SL (2009) Ultrafast and memory-efficient alignment of short DNA sequences to the human genome. Genome Biol. 10: R25 Available at: http://genomebiology.biomedcentral.com/articles/10.1186/gb-2009-10-3-r25

62. Li F, Pignatta D, Bendix C, Brunkard JO, Cohn MM, Tung J, Sun H, Kumar P & Baker B (2012) MicroRNA regulation of plant innate immune receptors. Proc. Natl. Acad. Sci. 109: 1790–1795 Available at: http://www.pnas.org/cgi/doi/10.1073/pnas.1118282109

63. Liu Y, Zhai H, Zhao K, Wu B & Wang X (2012) Two suppressors of RNA silencing encoded by cereal-infecting members of the family Luteoviridae. J. Gen. Virol. 93: 1825–1830

64. López-Márquez D, Del-Espino Á, López-Pagán N, Rodríguez-Negrete EA, Rubio-Somoza I, Ruiz-Albert J, Bejarano ER & Beuzón CR (2020) MiRNA and phasiRNAs-mediated regulation of TIR-NBS-LRR defense genes in Arabidopsis thaliana. bioRxiv: 2020.03.02.972620

65. Ma X, Nicole MC, Meteignier LV, Hong N, Wang G & Moffett P (2015) Different roles for RNA silencing and RNA processing components in virus recovery and virus-induced gene silencing in plants. J. Exp. Bot. 66: 919–932

66. Mallory AC & Vaucheret H (2009) ARGONAUTE 1 homeostasis invokes the coordinate action of the microRNA and siRNA pathways. EMBO Rep. 10: 521–526

67. Mangwende T, Wang ML, Borth W, Hu J, Moore PH, Mirkov TE & Albert HH (2009) The P0 gene of Sugarcane yellow leaf virus encodes an RNA silencing suppressor with unique activities. Virology 384: 38–50 Available at: http://dx.doi.org/10.1016/j.virol.2008.10.034

68. Matsuda Y, Liang G, Zhu Y, Ma F, Nelson RS & Ding B (2002) The Commelina yellow mottle virus promoter drives companion-cell-specific gene expression in multiple organs of transgenic tobacco. Protoplasma 220: 51–58 Available at: http://link.springer.com/10.1007/s00709-002-0027-6

69. Mayo MA & Ziegler-Graff V (1996) Molecular biology of luteoviruses. Adv. Virus Res. 46: 413–460

70. Michaeli S, Clavel M, Lechner E, Viotti C, Wu J, Dubois M, Hacquard T, Derrien B, Izquierdo E, Lecorbeiller M, Bouteiller N, De Cilia J, Ziegler-Graff V, Vaucheret H, Galili G & Genschik P (2019) The viral F-box protein P0 induces an ER-derived autophagy degradation pathway for the clearance of membrane-bound AGO1. Proc. Natl. Acad. Sci. 116: 22872–22883 Available at: http://www.pnas.org/lookup/doi/10.1073/pnas.1912222116

71. Monsion B, Incarbone M, Hleibieh K, Poignavent V, Ghannam A, Dunoyer P, Daeffler L, Tilsner J & Ritzenthaler C (2018) Efficient Detection of Long dsRNA in Vitro and in Vivo Using the dsRNA Binding Domain from FHV B2 Protein. Front. Plant Sci. 9: 1–16 Available at: http://journal.frontiersin.org/article/10.3389/fpls.2018.00070/full

72. Montavon T, Kwon Y, Zimmermann A, Hammann P, Vincent T, Cognat V, Bergdoll M, Michel F & Dunoyer P (2018) Characterization of DCL4 missense alleles provides insights into its ability to process distinct classes of dsRNA substrates. Plant J. 95: 204–218

73. Moran PJ & Thompson GA (2001) Molecular Responses to Aphid Feeding in Arabidopsis in Relation to Plant Defense Pathways. Plant Physiol. 125: 1074–1085 Available at: https://academic.oup.com/plphys/article/125/2/1074-1085/6099770

74. Morel J, Godon C, Mourrain P, Feuerbach F & Proux F (2002) Fertile Hypomorphic ARGONAUTE (ago1) Mutants Impaired in Post-Transcriptional Gene Silencing and Virus Resistance. Plant Cell 14: 629–639

75. Mourrain P, Bé C, Elmayan T, Feuerbach F, Godon C, Morel J-B, Jouette D, Lacombe A-M, Nikic S, Picault N, Ré K, Sanial M, Vo T-A & Vaucheret H (2000) Arabidopsis SGS2 and SGS3 Genes Are Required for Posttranscriptional Gene Silencing and Natural Virus Resistance gene required for PTGS, a possible mechanistic link. Cell 101: 533–542

76. Navarro B, Gisel A, Rodio ME, Delgado S, Flores R & Di Serio F (2012) Small RNAs containing the pathogenic determinant of a chloroplast-replicating viroid guide the degradation of a host mRNA as predicted by RNA silencing. Plant J. 70: 991–1003 Available at: http://doi.wiley.com/10.1111/j.1365-313X.2012.04940.x

77. Pazhouhandeh M, Dieterle M, Marrocco K, Lechner E, Berry B, Brault V, Hemmer O, Kretsch T, Richards KE, Genschik P & Ziegler-Graff V (2006) F-box-like domain in the polerovirus protein P0 is required for silencing suppressor function. Proc. Natl. Acad. Sci. U. S. A. 103: 1994–1999

78. Peragine A (2004) SGS3 and SGS2/SDE1/RDR6 are required for juvenile development and the production of trans-acting siRNAs in Arabidopsis. Genes Dev. 18: 2368–2379 Available at: http://www.genesdev.org/cgi/doi/10.1101/gad.1231804

79. Pfeffer S, Dunoyer P, Heim F, Richards KE & Jonard G (2002) P0 of Beet Western Yellows Virus Is a Suppressor of Posttranscriptional Gene Silencing P0 of Beet Western Yellows Virus Is a Suppressor of Posttranscriptional Gene Silencing. J. Virol. 76: 6815–6824

80. Poulsen C, Vaucheret H & Brodersen P (2013) Lessons on RNA Silencing Mechanisms in Plants from Eukaryotic Argonaute Structures. Plant Cell 25: 22–37 Available at: http://www.plantcell.org/lookup/doi/10.1105/tpc.112.105643

81. Pumplin N & Voinnet O (2013) RNA silencing suppression by plant pathogens: defence, counter-defence and counter-counter-defence. Nat. Rev. Microbiol. 11: 745–760 Available at: http://www.nature.com/articles/nrmicro3120

82. Qu F, Ye X & Morris TJ (2008) Arabidopsis DRB4, AGO1, AGO7, and RDR6 participate in a DCL4-initiated antiviral RNA silencing pathway negatively regulated by DCL1. Proc. Natl. Acad. Sci. 105: 14732–14737 Available at: http://www.pnas.org/cgi/doi/10.1073/pnas.0805760105

83. Ramesh S V, Yogindran S, Gnanasekaran P, Chakraborty S, Winter S & Pappu HR (2021) Virus and Viroid-Derived Small RNAs as Modulators of Host Gene Expression: Molecular Insights Into Pathogenesis. Front. Microbiol. 11: Available at: https://www.frontiersin.org/articles/10.3389/fmicb.2020.614231/full

84. Ramírez F, Ryan DP, Grüning B, Bhardwaj V, Kilpert F, Richter AS, Heyne S, Dündar F & Manke T (2016) deepTools2: a next generation web server for deep-sequencing data analysis. Nucleic Acids Res. 44: W160–W165 Available at: https://academic.oup.com/nar/article-lookup/doi/10.1093/nar/gkw257

85. Ratcliff F, Martin-Hernandez AM & Baulcombe DC (2001) Tobacco rattle virus as a vector for analysis of gene function by silencing. Plant J. 25: 237–245 Available at: http://www.ncbi.nlm.nih.gov/entrez/query.fcgi?cmd=Retrieve&db=PubMed&dopt=Citation&list_uids=11169199

86. Reutenauer A, Ziegler-Graff V, Lot H, Scheidecker D, Guilley H, Richards K & Jonard G (1993) Identification of beet western yellows luteovirus genes implicated in viral replication and particle morphogenesis. Virology 195: 692–699

87. Sadowy E, Maasen A, Juszczuk M, David C, Zagórski-Ostoja W, Gronenborn B & Hulanicka MD (2001) The ORF0 product of Potato leafroll virus is indispensable for virus accumulation. J. Gen. Virol. 82: 1529–1532 Available at: https://www.microbiologyresearch.org/content/journal/jgv/10.1099/0022-1317-82-6-1529

88. Schindelin J, Arganda-Carreras I, Frise E, Kaynig V, Longair M, Pietzsch T, Preibisch S, Rueden C, Saalfeld S, Schmid B, Tinevez J-Y, White DJ, Hartenstein V, Eliceiri K, Tomancak P & Cardona A (2012) Fiji: an open-source platform for biological-image analysis. Nat. Methods 9: 676–682 Available at: http://www.nature.com/articles/nmeth.2019

89. Schott G, Mari-Ordonez A, Himber C, Alioua A, Voinnet O & Dunoyer P (2012) Differential effects of viral silencing suppressors on siRNA and miRNA loading support the existence of two distinct cellular pools of ARGONAUTE1. EMBO J. 31: 2553–2565 Available at: http://dx.doi.org/10.1038/emboj.2012.92

90. Seguin J, Otten P, Baerlocher L, Farinelli L & Pooggin MM (2014) MISIS: A bioinformatics tool to view and analyze maps of small RNAs derived from viruses and genomic loci generating multiple small RNAs. J. Virol. Methods 195: 120–122 Available at: http://dx.doi.org/10.1016/j.jviromet.2013.10.013

91. Shen C, Wei C, Li J, Zhang X, Zhong Q, Li Y, Bai B & Wu Y (2020) Barley yellow dwarf virus-GAV-derived vsiRNAs are involved in the production of wheat leaf yellowing symptoms by targeting chlorophyll synthase. Virol. J. 17: 158 Available at: https://virologyj.biomedcentral.com/articles/10.1186/s12985-020-01434-7

92. Shimura H, Pantaleo V, Ishihara T, Myojo N, Inaba J, Sueda K, Burgyán J & Masuta C (2011) A Viral Satellite RNA Induces Yellow Symptoms on Tobacco by Targeting a Gene Involved in Chlorophyll Biosynthesis using the RNA Silencing Machinery. PLoS Pathog. 7: e1002021 Available at: https://dx.plos.org/10.1371/journal.ppat.1002021

93. Shivaprasad P V., Chen H-M, Patel K, Bond DM, Santos BACM & Baulcombe DC (2012) A MicroRNA Superfamily Regulates Nucleotide Binding Site-Leucine-Rich Repeats and Other mRNAs. Plant Cell 24: 859–874 Available at: http://www.plantcell.org/cgi/doi/10.1105/tpc.111.095380

94. Silva TF, Romanel EA, Andrade RR, Farinelli L, Østerås M, Deluen C, Corrêa RL, Schrago CE & Vaslin MF (2011) Profile of small interfering RNAs from cotton plants infected with the polerovirus Cotton leafroll dwarf virus. BMC Mol. Biol. 12: 40

95. Smith NA, Eamens AL & Wang M-B (2011) Viral Small Interfering RNAs Target Host Genes to Mediate Disease Symptoms in Plants. PLoS Pathog. 7: e1002022 Available at: https://dx.plos.org/10.1371/journal.ppat.1002022

96. Srivastava AC, Ganesan S, Ismail IO & Ayre BG (2009) Effective carbon partitioning driven by exotic phloem-specific regulatory elements fused to the Arabidopsis thaliana AtSUC2 sucrose-proton symporter gene. BMC Plant Biol. 9: 7 Available at: http://bmcplantbiol.biomedcentral.com/articles/10.1186/1471-2229-9-7

97. Sun J, Li L, Wang P, Zhang S & Wu J (2017) Genome-wide characterization, evolution, and expression analysis of the leucine-rich repeat receptor-like protein kinase (LRR-RLK) gene family in Rosaceae genomes. BMC Genomics 18: 763 Available at: http://bmcgenomics.biomedcentral.com/articles/10.1186/s12864-017-4155-y

98. Svozil J, Gruissem W & Baerenfaller K (2015) Proteasome targeting of proteins in Arabidopsis leaf mesophyll, epidermal and vascular tissues. Front. Plant Sci. 6: 1–17 Available at: http://www.frontiersin.org/Plant_Systems_and_Synthetic_Biology/10.3389/fpls.2015.00376/abstract

99. Thompson AR, Doelling JH, Suttangkakul A & Vierstra RD (2005) Autophagic Nutrient Recycling in Arabidopsis Directed by the ATG8 and ATG12 Conjugation Pathways. Plant Physiol. 138: 2097–2110 Available at: http://www.plantphysiol.org/cgi/doi/10.1104/pp.105.060673 [Accessed June 2, 2019]

100. Tilsner J, Linnik O, Wright KM, Bell K, Roberts AG, Lacomme C, Santa Cruz S & Oparka KJ (2012) The TGB1 Movement Protein of Potato virus X Reorganizes Actin and Endomembranes into the X-Body, a Viral Replication Factory. Plant Physiol. 158: 1359–1370 Available at: http://www.plantphysiol.org/lookup/doi/10.1104/pp.111.189605

101. Vazquez F, Vaucheret H, Rajagopalan R, Lepers C, Gasciolli V, Mallory AC, Hilbert J-L, Bartel DP & Crété P (2004) Endogenous trans-Acting siRNAs Regulate the Accumulation of Arabidopsis mRNAs. Mol. Cell 16: 69–79 Available at: https://linkinghub.elsevier.com/retrieve/pii/S1097276504005817

102. Waltz F, Nguyen TT, Arrivé M, Bochler A, Chicher J, Hammann P, Kuhn L, Quadrado M, Mireau H, Yashem Y & Giegé P (2019) Small is big in Arabidopsis mitochondrial ribosome. Nat. Plants 5: SOUS PRESSE Available at: http://dx.doi.org/10.1038/s41477-018-0339-y

103. Wang X-B, Jovel J, Udomporn P, Wang Y, Wu Q, Li W-X, Gasciolli V, Vaucheret H & Ding S-W (2011) The 21-Nucleotide, but Not 22-Nucleotide, Viral Secondary Small Interfering RNAs Direct Potent Antiviral Defense by Two Cooperative Argonautes in Arabidopsis thaliana. Plant Cell 23: 1625–1638 Available at: http://www.plantcell.org/lookup/doi/10.1105/tpc.110.082305

104. Wang XBX-B, Wu Q, Ito T, Cillo F, Li W-XWX, Chen X, Yu JLJ-L & Ding SWS-W (2010) RNAi-mediated viral immunity requires amplification of virus-derived siRNAs in Arabidopsis thaliana. Proc. Natl. Acad. Sci. U. S. A. 107: 484–489

105. Yang Y, Liu T, Shen D, Wang J, Ling X, Hu Z, Chen T, Hu J, Huang J, Yu W, Dou D, Wang MB & Zhang B (2019) Tomato yellow leaf curl virus intergenic siRNAs target a host long noncoding RNA to modulate disease symptoms. PLoS Pathog. 15: 1–22

106. Yang Z & Li Y (2018) Dissection of RNAi-based antiviral immunity in plants. Curr. Opin. Virol. 32: 88–99

107. Zhang XP, Liu DS, Yan T, Fang XD, Dong K, Xu J, Wang Y, Yu JL & Wang XB (2017) Cucumber mosaic virus coat protein modulates the accumulation of 2b protein and antiviral silencing that causes symptom recovery in planta. PLoS Pathog. 13: 1–25

108. Ziegler-Graff V, Brault V, Mutterer J, Simonis M-T, Herrbach E, Guilley H, Richards KE & Jonard G (1996) The Coat Protein of Beet Western Yellows Luteovirus is Essential for Systemic Infection but the Viral Gene Products P29 and P19 are Dispensable for Systemic Infection and Aphid Transmission. Mol. Plant-Microbe Interact. 9: 501–510 Available at: http://www.apsnet.org/publications/mpmi/backissues/Documents/1996Abstracts/Microbe09-501.htm

